# LSH-mediated resolution of R-loops mitigates transcription-replication conflicts to preserve genomic stability in prostate cancer cells

**DOI:** 10.1101/2025.06.04.657812

**Authors:** Kai Ni, Yu Zhang, Kun Zheng, Jianwen Huang, Nuorong Xiong, Rong Liang, Yingxiao Chen, Honglin Jin, Youlong Hai, Wang Li, Jiayi Ma, Ranxing Yang, Guanggai Xia, Yueqing Bai, Lujie Song, Xiaoyong Hu, Zijian Tang, Qiang Fu

**Affiliations:** Department of Urology, Shanghai Sixth People’s Hospital Affiliated to Shanghai Jiao Tong University School of Medicine, 200233, Shanghai, China; College of Biomedicine and Health, Huazhong Agricultural University, Wuhan, Hubei, 430070, China; Department of Urology, Sun Yat-sen Memorial Hospital, Sun Yat-Sen University, Guangzhou, 510289, China; Department of Urology, Shanghai Pudong Hospital, Fudan University Pudong Medical Center, Shanghai, 201399, China; Department of general surgery, Shanghai Sixth People’s Hospital Affiliated to Shanghai Jiao Tong University School of Medicine, 200233, Shanghai, China; Department of Pathology, Shanghai Sixth People’s Hospital Affiliated to Shanghai Jiao Tong University School of Medicine, 200233, Shanghai, China

## Abstract

Prostate cancer cells exhibit heightened transcriptional activity to sustain their aggressive phenotype, yet this creates DNA topological stress, leading to RNA: DNA hybrids (R-loops) and DNA damage that compromise genomic stability. LSH, an SNF2-family chromatin remodeler, is frequently dysregulated in cancers and linked to tumor progression, but its mechanistic role in genome maintenance remains unclear. Here, we demonstrate that LSH resolves R-loops in an ATP-dependent manner, mitigating transcription-replication (TR) conflicts in prostate cancer cells. LSH depletion caused R-loop accumulation due to reduced FANCD2 recruitment, exacerbating DNA damage and impairing Rad51 filament formation. Strikingly, LSH deficiency also increased R-loops at promoters of MYC and E2F target genes, stalling RNA polymerase II elongation and disrupting transcriptional programs critical for proliferation. Our findings reveal LSH is a key guardian of genome integrity, resolving R-loops to prevent TR conflicts and maintaining DNA repair fidelity. By linking LSH to R-loop resolution and transcriptional regulation, this study reveals its dual role in prostate cancer pathogenesis— supporting malignant proliferation while safeguarding genomic stability. These insights nominate LSH as a potential therapeutic target to disrupt oncogenic transcription and induce synthetic lethality in prostate cancer.

## Introduction

Prostate cancer is the third most common malignant tumor worldwide. Particularly among the male population, its incidence ranks second globally, and it is the fifth leading cause of cancer-related deaths in men[1–3]. We have already undertaken a comprehensive and in-depth investigation into the molecular biological mechanisms that underlie malignant prostate cancer. Prostate cancer cells demand an exceptionally high transcription rate to maintain their aggressive phenotype[4, 5]. This elevated transcriptional activity is essential not only for driving unlimited cell proliferation but also for ensuring robust expression of cancer-related genes[5–7]. However, excessive transcriptional activity can lead to considerable DNA topological stress, resulting in the formation of RNA: DNA hybrids (R-loops) and DNA double- strand breaks (DSBs), ultimately triggering genomic instability[8–11]. Studies have demonstrated that R-loops play significant roles in driving tumorigenesis and progression in prostate cancer[12–15].

R-loops are three-stranded hybrid structures composed of RNA: DNA complexes[9]. During DNA replication, when the DNA double helix unwinds to form a replication fork, R-loop structures can easily form if an RNA polymer acts complementarily with one of the DNA strands[8–10]. As a product generated during transcription, various factors participate in the regulation of R-loops, including RNA transcription and processing, chromatin modifications, and DNA damage repair[9, 16]. R-loops are closely associated with DNA damage, transcriptional elongation defects, excessive recombination, and genomic instability[8–11, 17]. Recent studies have indicated that a significant number of R-loop structures exist in cancer cells, serving as a major barrier to the progression of the DNA replication fork and causing replication stress[8, 9, 18]. Particularly, R-loops formation during the initial stages of transcription can have genotoxic effects[9–11]. In rapidly proliferating cancer cells, conflicts between DNA replication and transcription represent one of the greatest threats to genomic stability[19, 20].

At the intersection of chromatin structure and function, chromatin remodelers are increasingly recognized as critical players in cancer research. Recent studies have identified several chromatin remodeling proteins that regulate R-loop formation, such as SWI/SNF, INO80, and ATRX[9, 12, 21, 22]. Lymphoid Specific Helicase (LSH), a member of the SNF2 family of ATP-dependent chromatin remodeling proteins[23–25]. LSH mutation leads to Immunodeficiency, Centromeric instability, and Facial dysmorphism syndrome type-4 (ICF-4), which is characterized by immunodeficiency, instability of chromosome centromeres, and abnormal DNA methylation[25–28]. LSH plays a role in regulating DNA methylation, histone modifications, histone variants, and nucleosome composition and positioning, which are critical components of the epigenetic landscape[23, 29–31]. LSH also participates in and regulates DNA damage repair pathways through various mechanisms, which are crucial for maintaining genomic stability and the synthesis of nascent DNA at the replication fork[23, 26, 32]. The loss of LSH can affect genomic instability, senescence, and sensitivity to tumor chemotherapy, all of which are closely linked to DNA damage repair[23, 33–35].

LSH in T-cell lymphoma plays a crucial role in reducing unscheduled R- loops[34], while its Arabidopsis homolog, DDM1, is involved in mediating R- loop resolution[36]. Furthermore, telomeres in ICF syndrome cells are susceptible to DNA damage due to the increased accumulation of RNA: DNA hybrids[37]. In our study, we demonstrated that the loss of LSH in prostate cancer cells increases R-loops and R-loop-dependent DNA breaks, as well as transcription-replication (T-R) conflicts. Furthermore, we elucidated that LSH eliminates R-loop structures through FANCD2 in prostate cancer cells, thereby alleviating transcriptional barriers, promoting gene transcription, and facilitating the elongation of RNA polymerase II (RNAPII). Our research reveals a novel function of LSH in helping to resolve R-loop mediated T-R conflicts, thereby maintaining genomic stability and playing a significant role in driving tumorigenesis and progression in prostate cancer cells. This research is crucial for identifying novel therapeutic targets and advancing the development of more effective treatment strategies.

## Results

### 1. LSH deficiency increases genome instability and impairs cell proliferation in prostate cancer cells

To investigate the role of human LSH protein in maintaining genome stability and tumorigenic phenotype, we developed a single-vector lentiviral system with doxycycline-inducible shRNA. This Tet-on LSH-shRNA system enables efficient and controllable knockdown of the LSH gene in prostate cancer PC3 cell line (**Figure 1A**). Treatment with doxycycline for five days (DOX 5d) resulted in a substantial reduction in LSH mRNA levels, decreasing by over 90% compared to untreated cells (Dox_0d) (**Figure S1A**). This reduction was further validated by a corresponding decrease in LSH protein levels, as shown through Western blotting (**Figure 1B**) and immunofluorescence analysis (**Figure S1B**). Intriguingly, five days after doxycycline washout, LSH expression was fully rescued (RESC) to normal levels at both mRNA (**Figure S1A**) and protein levels **(Figure 1B and Figure S1B**). To begin, we investigated whether LSH depletion would induce genome instability in PC3 cells. Our findings showed an increase in anaphase bridges and micronuclei in Tet-on LSH-shRNA PC3 cells treated with doxycycline, and this instability was mitigated upon restoring LSH expression following a five-day washout period (**Figure 1C-D**). We then investigated the role of LSH in the DNA damage response and maintenance of genome integrity in PC3 cells. Following doxycycline treatment, LSH deletion in DOX_5d PC3 cells led to an accumulation of DNA breaks, as indicated by γH2AX immunostaining (**Figure 1E**) and by the results of the alkaline comet assay (**Figure 1F**), in contrast to DOX_0d cells. Upon doxycycline removal, re-expression of LSH in RESC cells restored DNA damage levels to normal (**Figure 1E-F**). In summary, our results demonstrate that LSH plays a crucial role in maintaining genomic stability and preventing DNA damage in PC3 cells.

**Figure 1.**
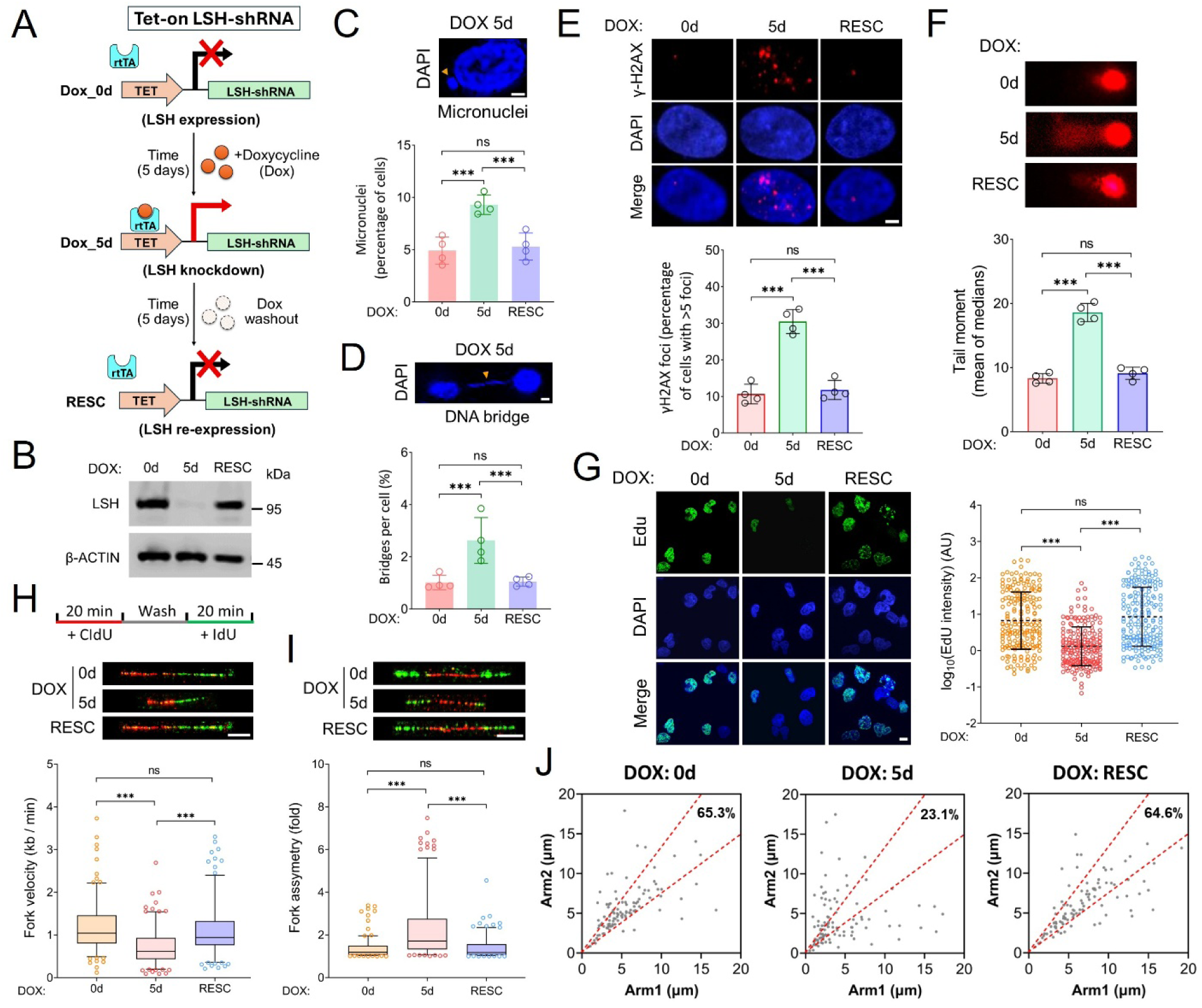
Analysis of DNA damage and genome instability in LSH knockdown PC3 cells using Tet-on LSH-shRNA system. **A** Schematic of the Tet-on LSH-shRNA system for regulating LSH expression in PC3 cells. Treatment with doxycycline for five days (DOX_5d) resulted in a significant reduction in LSH expression compared to cells that did not receive doxycycline treatment (DOX_0d). Following a five-day washout period, LSH expression was fully rescued to normal level (RESC). **B** Doxycycline- inducible knockdown of LSH was assessed using Western blotting to track alterations in protein levels, with endogenous β-ACTIN serving as the loading control. **C-D** Representative images and the percentage of cells with micronuclei (**C**) or DNA bridge (**D**) in Tet-on LSH-shRNA PC3 cells treated as described in panel A. The scale bar represents 5 μm. Data are presented as mean ± s.d. (n = 4 biologically independent experiments). ***p < 0.001; n.s. indicates not significant, as determined by the two-tailed Mann–Whitney test. **E** Representative images of γ-H2AX immunostaining and bar graph show the percentage of PC3 cells containing more than five γ-H2AX foci, treated as described in panel A. The scale bar represents 5 μm. Data are presented as mean ± s.d. (n = 4 biologically independent experiments). ***p < 0.001; n.s. indicates not significant, as determined by the two-tailed Mann– Whitney test. **F** Representative images of the alkaline comet assay and quantification of tail moment in PC3 cells, treated as described in panel A. The scale bar represents 5 μm. Data are presented as mean ± s.d. (n = 4 biologically independent experiments). ***p < 0.001; n.s. indicates not significant, as determined by the two-tailed Mann–Whitney test. **G** Representative images of Edu staining and quantification of nuclear Edu signal intensity in PC3 cells, treated as described in panel A. The scale bar represents 10 μm. Signal intensity data are depicted as a scatter plot with mean ± s.d. (n = 4 biologically independent experiments). ***p < 0.001; n.s. indicates not significant, as determined by the two-tailed Mann–Whitney test. AU denotes arbitrary units. **H-I** Quantification of replication fork velocity (**H**) and fork asymmetry (**I**) is presented using Tukey-style box plots. Schematic diagram of the DNA fiber assay and representative images of stretched DNA fibers in PC3 cells treated as described in panel A. Cells were sequentially labeled at the indicated time points with two different thymidine analogues— CldU (red) and IdU (green). The scale bar represents 10 μm. The central line of a Tukey-style box plot indicates the median value, while the boxes and whiskers represent the 25th to 75th percentiles and the 5th to 95th percentiles, respectively. Outliers are displayed as individual points. Data are representative of n = 4 biologically independent experiments. ***p < 0.001; n.s. indicates not significant, as determined by the One-way ANOVA with Tukey’s multiple comparison test. **J** Scatter plots depicting the distances covered by right-moving and left-moving bi-directional replication forks following the CldU pulse in PC3 cells treated as described in panel H. The central areas, delineated by red lines, represent bi-directional forks with a length difference of less than 25%. The percentage of symmetric forks is also indicated. Data are representative of n = 4 biologically independent experiments. Source data are provided as a Source Data file.

Moreover, LSH plays a crucial role in safeguarding nascent DNA at stalled replication forks, with LSH-mediated DNA breaks occurring primarily due to replication fork stalling[26]. To investigate the impact of LSH depletion on DNA replication dynamics, we measured 5-ethynyl-2′-deoxyuridine (EdU) incorporation as an indicator of DNA synthesis. Our observations revealed a reduction in EdU incorporation in LSH-depleted cells compared to cells without doxycycline treatment (**Figure 1G)**. Additionally, the percentage of EdU-positive cells was reduced following LSH depletion (**Figure S1C**). Both effects were significantly rescued by re-expression of LSH in RESC cells (**Figure 1G and Figure S1C**). To further investigate the mechanism by which LSH prevents DNA damage formation at the replication fork, we conducted DNA fiber analysis. This technique allows for the examination of genome- wide replication fork dynamics at the single-molecule level using double pulse labeling with two sequential thymidine analogs, 5-iodo-2′-deoxyuridine (IdU) and 5-chloro-2′-deoxyuridine (CldU). Our analysis revealed significant decreases in replication fork velocity (**Figure 1H**) and increases in replication fork asymmetry (**Figure 1I**) in LSH-deficient cells (DOX_5d) compared to control cells (DOX_0d). Notably, re-expressing LSH (RESC) mitigated these effects, restoring fork velocity and asymmetry to the levels closer to normal (**Figure 1H-I**). To differentiate between slower fork progression and increased fork stalling in LSH-deficient PC3 cells, we examined the progression of sister replication forks. In control cells (DOX_0d), most sister forks advanced at similar rates from a common origin, resulting in symmetrical IdU/CldU incorporation patterns (65%). In contrast, LSH- depleted cells (DOX_5d) exhibited only 23% symmetrical fork progression, with a greater than two-fold difference in DNA synthesis rates between sister forks compared to controls (**Figure 1J**). This finding suggests a higher frequency of fork stalling in LSH-deficient PC3 cells. Importantly, re- expression of LSH in RESC cells restored fork symmetry (**Figure 1J**), strongly indicating that LSH deficiency leads to increased replication fork stalling. Overall, these results suggest that LSH prevents DNA damage formation at the replication fork, thereby mitigating DNA breaks and contributing to genomic stability.

Depletion of LSH induces DNA damage and compromises genomic integrity in PC3 cells, suggesting that LSH is essential for maintaining genomic stability under physiological conditions. In its absence, unrepaired DNA damage may trigger apoptosis and inhibit cell growth. To evaluate apoptosis levels, TUNEL assay results showed that LSH deficiency leads to significantly higher apoptosis compared to cells without doxycycline treatment, while LSH re-expression restores apoptosis to normal levels (**Figure S1D**). For cell growth assessment, a colony formation assay revealed a marked inhibition of PC3 cell growth following LSH deletion. Notably, LSH restoration after doxycycline washout rescues cell growth (**Figure S1E**). These results are consistent with the CCK-8 assay, which also demonstrated reduced cell viability in the absence of LSH (**Figure S1F**).

Furthermore, the doubling time of LSH-deficient cells increased by over 10 hours compared to wild-type PC3 cells, with LSH re-expression returning the rate to normal, as indicated by proliferation curves from Tet-on LSH-shRNA PC3 cells (**Figure S1G**). Therefore, the absence of LSH in PC3 prostate cancer cells induces unrepaired DNA damage, leading to cell apoptosis and inhibited cell growth.

### 2. LSH suppresses R-loop accumulation and promotes DNA replication

Considering that DDM1, the Arabidopsis homolog of LSH, is involved in mediating R-loop resolution and maintaining genome stability[36], we investigated whether the depletion of human LSH would result in an increase in R-loops in prostate cancer cells. Immunofluorescence analysis of R-loops using the S9.6 anti-DNA–RNA monoclonal antibody revealed a significant increase in the nuclear S9.6 signal in LSH-depleted PC3 cells after five days of doxycycline treatment (DOX_5d), compared to control cells without doxycycline treatment (DOX_0d) (**Figure 2A**). Additionally, in the rescue experiment, five days of doxycycline washout restored R-loop levels through LSH re-expression (RESC) (**Figure 2A**). The results indicate that LSH plays a role in regulating the accumulation of R-loops in the PC3 cells. To further validate the results, we overexpressed wild-type (WT) RNASEH1 protein, tagged with V5, which is known to suppress R-loop formation[9, 38, 39]. In parallel, we overexpressed mutant RNASEH1 variants, including the WKKD and D210N mutants. The WKKD mutation disrupts both the binding and catalytic activities of RNASEH1, whereas D210N selectively abolishes catalytic activity while preserving binding capability[38, 39]. We employed the WKKD mutant as a control for comparison with WT RNASEH1, while the D210N mutant was utilized in subsequent experiments. The overexpression of these proteins was confirmed by western blot (**Figure 2B**) and immunofluorescence (**Figure S2A**) using V5 antibody. Immunostaining of R- loops in WT RNASEH1-overexpressing cells revealed a significant decrease in R-loop levels in LSH-depleted PC3 cells, with or without doxycycline treatment. In contrast, R-loop levels were markedly increased in WKKD mutant cells lacking LSH expression after doxycycline treatment (**Figure 2C**). To substantiate our findings, we further employed another prostate cancer cell line, LNCaP. We utilized lentiviral vectors to knock down (KD) LSH and introduced WT RNASEH1-V5 overexpression (KD+RNH1) in the cells. The expression of these proteins was validated through western blot analysis (**Figure S3A**). Similarly, the immunostaining of R-loops in LSH KD cells showed significantly increased signals compared to LSH WT cells (**Figure S3B**). Furthermore, R-loop levels were significantly reduced in RNASEH1- overexpressing (KD+RNH1) cells relative to LSH KD cells (**Figure S3B**). Therefore, these results revealed that LSH also suppresses R-loop formation in LNCaP cells. To assess whether LSH depletion induces increased DNA damage due to R-loop accumulation, we conducted γ-H2AX staining and alkaline comet assays in Tet-on LSH-shRNA PC3 cells overexpressing WT or WKKD RNASEH1. Tet-on PC3 cells overexpressing WT RNASEH1 led to a significant decrease in both γ-H2AX foci formation (**Figure 2D**) and DNA damage as measured by tail moment in alkaline comet assays (**Figure 2E**), irrespective of doxycycline treatment. In contrast, LSH KD (DOX_5d) cells overexpressing WKKD mutant exhibited increased γ-H2AX staining (**Figure 2D**) and higher tail moments in the comet assay (**Figure 2E**) compared to cells not treated with doxycycline (DOX_0d). Furthermore, LSH knockdown in LNCaP cells resulted in elevated γ-H2AX foci formation (**Figure S3C**) and increased DNA damage as measured by comet assay tail moments (**Figure S3D**). In contrast, overexpression of WT RNASEH1 significantly reduces both γ-H2AX staining and tail moments in LSH-deficient LNCaP cells (**Figure S3C-D**). Consistent with these findings, we observed an increase in micronuclei (**Figure S2B**) and anaphase bridges (**Figure S2C**) in LSH- depleted PC3 cells overexpressing WKKD RNASEH1 mutant compared to those without doxycycline treatment. Notably, both increases were suppressed by the overexpression of WT RNASEH1 (**Figure S2B-C**). In LNCaP cells, we also observed a higher percentage of micronuclei and anaphase bridges in LSH knockdown cells, both of which could be rescued by overexpressing WT RNASEH1 (**Figure S3E**). Thus, DNA damage observed in LSH-deficient PC3 and LNCaP cells is dependent on R-loop accumulation.

**Figure 2.**
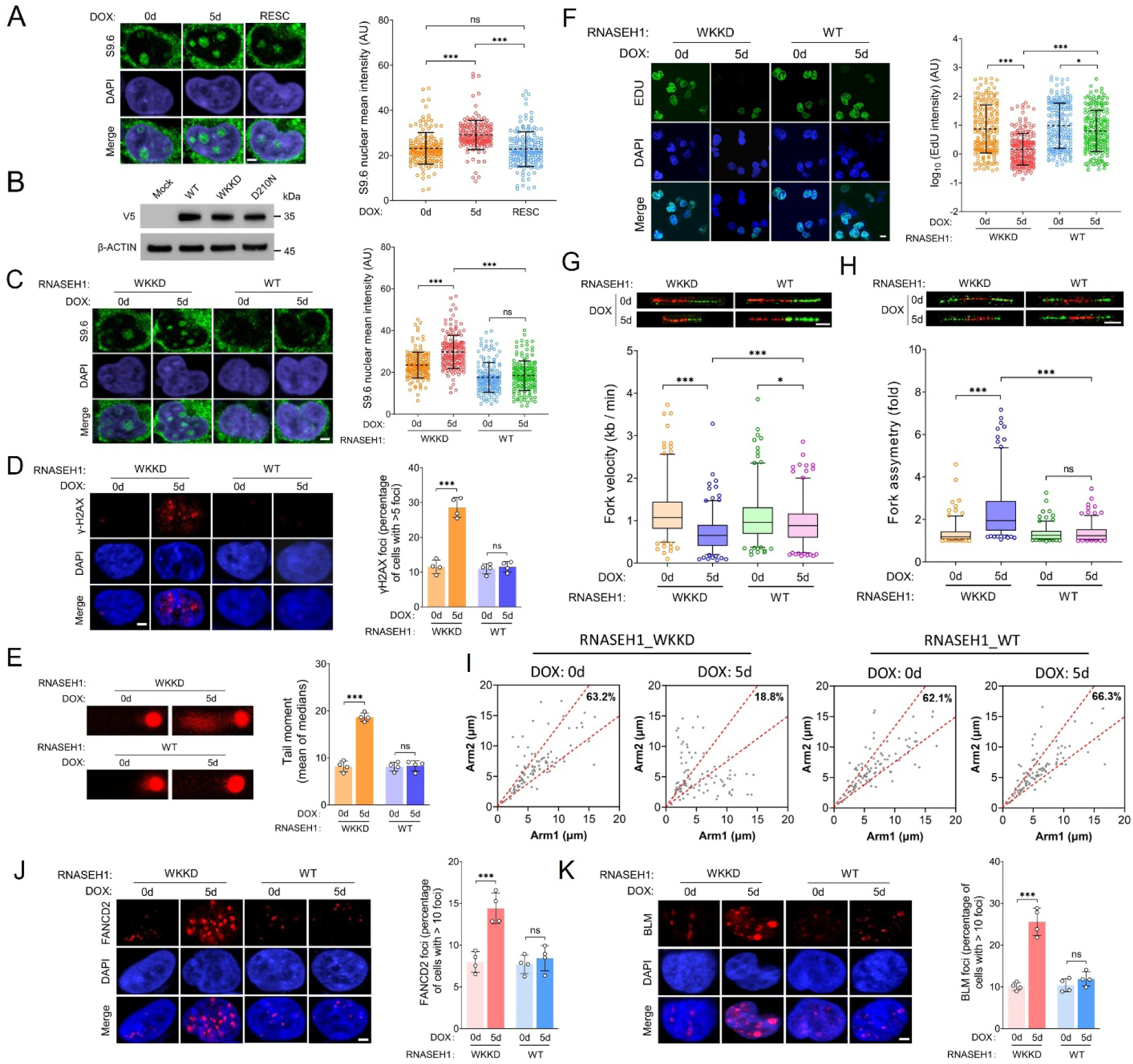
Effects of LSH knockdown on R-loop accumulation and DNA replication dynamics in PC3 cells. **A** Representative S9.6 immunostaining images showing R-loop levels in PC3 Tet-on LSH-shRNA cells under different conditions: untreated (Dox_0d), doxycycline-treated for five days (Dox_5d), and after a five-day doxycycline washout period (RESC). The scale bar represents 5 μm. Signal intensity data are depicted as a scatter plot with mean ± s.d. (n = 4 biologically independent experiments). ***p < 0.001; n.s. indicates not significant, as determined by the two-tailed Mann– Whitney test. AU denotes arbitrary units. **B** Western blot analysis of PC3 Tet- on LSH-shRNA cells demonstrated the successful overexpression of both wild-type (WT) RNASEH1 and its WKKD/D210N mutant variants, all of which were tagged with V5. Detection was performed using an anti-V5 antibody, with β-Actin serving as the loading control. **C** Representative images of S9.6 immunostaining were utilized to assess R-loop levels in DOX_0d and DOX_5d cells overexpressing either WT or WKKD RNASEH1. The scale bar represents 5 μm. Signal intensity data are depicted as a scatter plot with mean ± s.d. (n = 4 biologically independent experiments). ***p < 0.001; n.s. indicates not significant, as determined by the two-tailed Mann–Whitney test. AU denotes arbitrary units. **D** Representative images of γ-H2AX immunostaining were obtained, alongside the percentage of cells exhibiting more than five γ-H2AX foci in DOX_0d and DOX_5d cells overexpressing either WT or WKKD RNASEH1. The scale bar represents 5 μm. Data are presented as mean ± s.d. (n = 4 biologically independent experiments). ***p < 0.001; n.s. indicates not significant, as determined by the two-tailed Mann– Whitney test. **E** Representative images of the alkaline comet assay, along with quantification of the tail moment, were obtained from DOX_0d and DOX_5d cells overexpressing either WT or WKKD RNASEH1. Data are presented as mean ± s.d. (n = 4 biologically independent experiments). ***p < 0.001; n.s. indicates not significant, as determined by the two-tailed Mann– Whitney test. **F** Representative images of EdU staining and the quantification of nuclear EdU signal intensity were obtained from DOX_0d and DOX_5d cells overexpressing either WT or WKKD RNASEH1. The scale bar represents 10 μm. Signal intensity data are depicted as a scatter plot with mean ± s.d. (n = 4 biologically independent experiments). *p < 0.05; ***p < 0.001, as determined by the two-tailed Mann–Whitney test. AU denotes arbitrary units. **G-H** Quantification of replication fork velocity (**G**) and fork asymmetry (**H**) is presented using Tukey-style box plots. Representative images depict stretched DNA fibers from DOX_0d and DOX_5d cells overexpressing either WT or WKKD RNASEH1. The cells were sequentially labeled with CldU (red) and IdU (green). The scale bar represents 10 μm. The central line of a Tukey-style box plot indicates the median value, while the boxes and whiskers represent the 25th to 75th percentiles and the 5th to 95th percentiles, respectively. Outliers are displayed as individual points. Data are representative of n = 4 biologically independent experiments. *p < 0.05; ***p < 0.001; n.s. indicates not significant, as determined by the One- way ANOVA with Tukey’s multiple comparison test. **I** Scatter plots depicting the distances covered by right-moving and left-moving bi-directional replication forks following the CldU pulse in DOX_0d and DOX_5d cells overexpressing either WT or WKKD RNASEH1. The central areas, delineated by red lines, represent bi-directional forks with a length difference of less than 25%. The percentage of symmetric forks is also indicated. Data are representative of n = 4 biologically independent experiments. **J-K** Representative images of FANCD2 (**J**) and BLM (**K**) immunostaining along with the percentage of cells exhibiting more than ten foci are presented for DOX_0d and DOX_5d cells overexpressing either WT or WKKD RNASEH1. The scale bar represents 5 μm. Data are presented as a bar graph with mean ± s.d. (n = 4 biologically independent experiments). ***p < 0.001; n.s. indicates not significant, as determined by the two-tailed Mann–Whitney test. Source data are provided as a Source Data file.

R-loop-mediated DNA breaks primarily occur because of replication fork stalling, as R-loops present a barrier to fork progression[9, 40]. Our previous findings highlight the critical role of LSH in protecting nascent DNA at stalled replication forks[26]. To investigate whether R-loops impair DNA replication as a primary mechanism underlying DNA damage in LSH-deficient cells, we conducted further analysis. First, the quantification of nuclear EdU signal intensity (**Figure 2F**) and the percentage of EdU-positive (**Figure S2D**) cells were significantly reduced in LSH knockdown cells overexpressing the WKKD RNASEH1 compared to control cells not treated with doxycycline. Notably, both phenotypes were partially but significantly rescued by the overexpression of WT RNASEH1 (**Figure 2F and Figure S2D**). Additionally, we found that nuclear EdU signal intensity was significantly reduced in LSH KD LNCaP cells compared to LSH WT cells. This reduction could be also restored by overexpressing WT RNASEH1 (**Figure S3F**). Furthermore, DNA fiber analysis to observe DNA-replication progression revealed significant decreases in replication fork velocity (**Figure 2G**) and increases in replication fork asymmetry (**Figure 2H**) in LSH deficient cells overexpressing WKKD RNASEH1 versus the control cells without doxycycline treatment. Importantly, both effects were suppressed by the overexpression of WT RNASEH1, restoring fork velocity and asymmetry to levels closer to those of the control cells (**Figure 2G-H**). Likewise, in LNCaP cells, both fork velocity and symmetry are compromised in the absence of LSH; however, these deficits were restored by overexpressing WT RNASEH1 (**Figure S3G-H**). We also examined the progression of sister replication forks. In Tet-on LSH shRNA PC3 cells without doxycycline treatment and overexpressing WKKD RNASEH1, most sister forks progressed at similar rates from a common origin, displaying symmetrical IdU/CldU incorporation patterns. However, LSH KD cells exhibited significantly higher asymmetrical fork progression compared to control cells (**Figure 2I**). Intriguingly, overexpression of WT RNASEH1 in LSH KD cells restored fork symmetry to levels observed in control cells (**Figure 2I**). We also analyzed the rates of sister forks in LNCaP cells. Consistent with the previous results, LSH KD causes increased asymmetrical fork progression, which can be rescued in KD+RNH1 cells (**Figure S3I**). To confirm that replication forks were stalled at R-loops, as indicated by the asymmetry of the forks, we conducted a further investigation into the presence of FANCD2 and BLM foci, which are known to accumulate at sites of replication fork stalling[41–43]. The percentage of cells displaying FANCD2 foci increased from approximately 7.5% in the control group to 14.5% in the LSH KD cells following the overexpression of WKKD RNASEH1 (**Figure 2J**). Similarly, the proportion of cells with BLM foci increased from approximately 10% in the control cells to 26% in LSH KD cells (**Figure 2K**).

Upon overexpression of WT RNASEH1, the percentage of cells exhibiting FANCD2 and BLM foci significantly decreased in LSH KD cells, reaching levels comparable to those of the control cells (**Figure 2J-K**). Overall, these findings indicate that R-loop accumulation in LSH KD cells leads to replication fork stalling and disrupts replication fork progression in prostate cancer cells.

Our research findings reveal that the absence of LSH results in persistent DNA damage, which correlates with a decrease in cellular proliferation and the induction of apoptosis (**Figure S1D-G**). We are intrigued by the possibility that attenuating R-loop accumulation through LSH might mitigate these phenotypic effects. In our analysis of cell growth, cell viability, as determined by the CCK-8 assay (**Figure S2E**), and the doubling time, as measured by the cell proliferation curve (**Figure S2F**), demonstrated that the overexpression of WT RNASEH1 can effectively restore cell growth in prostate cancer PC3 cells lacking LSH. In contrast, PC3 cells overexpressing WKKD RNASEH1 were unable to restore cell growth following LSH depletion. The results of the colony formation assay further corroborate these findings (**Figure S2G**). Similarly, in LNCaP cells, we found that cell viability decreased, and doubling time increased in LSH KD cells, both of which were restored in KD+RNH1 cells (**Figure S3J-K**). Regarding cell apoptosis, our TUNEL assay data revealed that overexpression of WT RNaseH1 could reverse the elevated apoptosis levels associated with LSH deficiency. Conversely, overexpression of WKKD RNASEH1 did not exhibit a similar rescue effect on cell apoptosis (**Figure S2H**). We also observed similar TUNEL results in LNCaP cells, showing that LSH deficiency leads to increased apoptosis, which can be rescued by overexpression of WT RNASEH1 (**Figure S3L**). To further validate the role of LSH in vivo, we performed a tumorigenesis assay by subcutaneously injecting LNCaP cells into M-NSG immunodeficient mice. Tumor volume was measured every 2–3 days. Upon reaching the experimental endpoint, all mice were euthanized, and tumors were excised for further analysis (**Figure S3M**). Notably, LSH KD cells exhibited significantly reduced tumor volume (**Figure S3N**) and tumor weight (**Figure S3O**) compared to LSH WT cells. However, these effects were rescued by overexpression RNASEH1, which restored tumor growth parameters to near-normal levels. Altogether, our results suggest that the absence of LSH leads to an increase accumulation of R-loops, consequently causing replication fork stalls and DNA breaks, which ultimately impairs cell growth and aggravates apoptosis in prostate cancer cells.

### 3. Genome-wide analysis of R-loop accumulation upon LSH deficiency

To further elucidate the role of LSH in suppressing R-loop formation, we conducted the cleavage under targets and tagmentation (CUT&Tag) assay using the S9.6 antibody to map genome-wide R-loop accumulation. Due to the influence of reaction time and drug concentration on doxycycline-induced gene knockdown, as well as the potential for certain non-specific effects, these factors may affect the results of whole-genome sequencing. Therefore, we utilized wild-type (WT) and LSH knockout (KO) PC3 cells to create a genome-wide R-loop map. Additionally, we included KO cells with overexpression of WT RNASEH1 (KO+RNH1) as a control group. The PC3 KO cell line was established using the CRISPR-Cas9 method, and stable overexpression of RNH1 was achieved via lentiviral infection. The protein expression was confirmed through western blot analysis (**Figure 3A**). Before conducting whole-genome sequencing, we first assessed the phenotypes of WT, KO and KO + RNH1 cells. Consistent with prior findings, immunostaining with the S9.6 antibody demonstrated that LSH KO results in an increased formation of R-loops compared to LSH WT cells. In contrast, KO+RNH1 cells displayed a significant reduction in R-loop levels (**Figure S4A**). Moreover, LSH KO also resulted in an accumulation of DNA breaks, as evidenced by the results of γH2AX immunostaining and the alkaline comet assay, when compared to WT cells. In contrast, KO + RNH1 cells showed a significant reduction in DNA breaks (**Figure S4B-C**). Similarly, we also observed a higher percentage of micronuclei and anaphase bridges in LSH KO cells, both of which could be rescued in KO+RNH1 cells (**Figure S4D**). Additionally, nuclear EdU signal intensity was significantly reduced in LSH KO cells compared to WT cells. This reduction could be partially rescued in KO + RNH1 cells (**Figure S4E**). Furthermore, both fork velocity and symmetry were compromised in the absence of LSH, but these deficits were restored in KO+RNH1 cells (**Figure S4F**). We also observed a decrease in cell viability (**Figure S4G**) and an increase in doubling time (**Figure S4H**) in LSH KO cells, both of which were rescued by the overexpression of RNASEH1. Additionally, the TUNEL assay revealed that LSH deficiency resulted in increased apoptosis, which was mitigated in KO+RNH1 cells (**Figure S4I**). Therefore, our results indicated that the characteristics of WT, KO and KO+RNH1 cells were consistent with our previous findings, which could be utilized to map genome-wide R-loop accumulation in PC3 cells.

**Figure 3.**
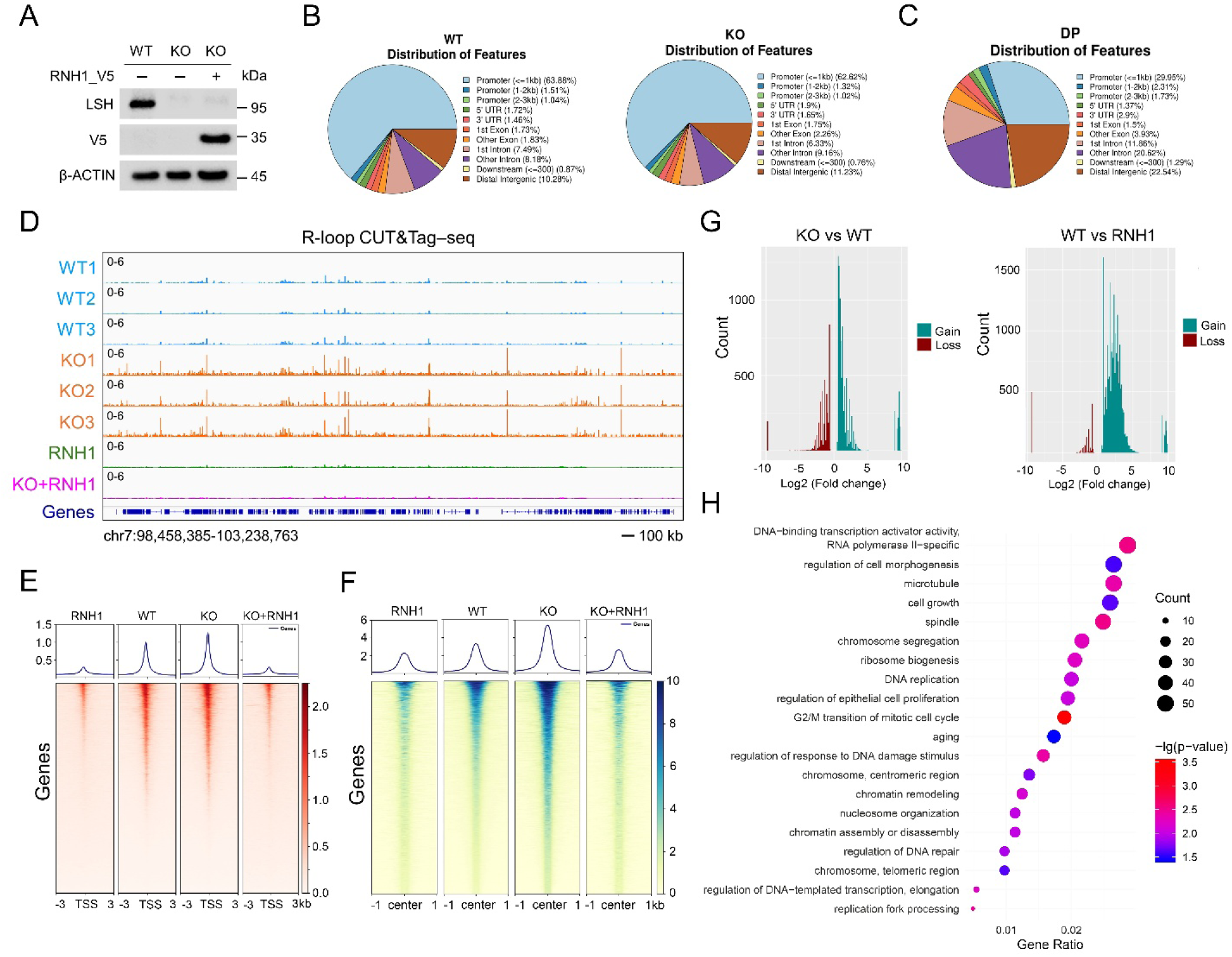
Genome-wide profiling demonstrates that LSH plays a crucial role in modulating R-loop accumulation in PC3 cells. **A** Western blot analysis confirmed the knockout (KO) of LSH in PC3 cells and the overexpression of wild-type RNH1_V5 in LSH KO cells (KO+RNH1). LSH was detected with an anti-LSH antibody, RNH1_V5 was identified using an anti-V5 antibody, and β-Actin was used as the loading control. **B** The genomic distribution of R-loop CUT&Tag peaks is presented for LSH WT and KO cells. UTR refers to the untranslated region. The data are representative of n = 3 biologically independent experiments. **C** The differences in the genomic distribution of R-loop CUT&Tag peaks between LSH WT and KO cells are illustrated. UTR refers to the untranslated region. The data are representative of n = 3 biologically independent experiments. **D** Genome browser view of R-loop CUT&Tag data representing three independent biological replicates of LSH WT and KO cells. Samples treated with the RNH1 enzyme (RNH1) and KO+RNH1 cells were utilized as negative control groups. **E** The average R-loop CUT&Tag signals are shown over 6 kb regions centered on the transcription start site (TSS) in LSH WT and KO cells. Samples treated with the RNH1 enzyme (RNH1) and KO+RNH1 cells served as negative control groups to confirm the specificity of the detected R-loop signals. **F** The average R-loop CUT&Tag read density and heatmap for LSH WT and KO cells are presented within a 1 kb window around the center of peak regions. Samples treated with the RNH1 enzyme (RNH1) and KO+RNH1 cells were used as negative control groups to confirm the specificity of the detected R-loop signals. **G** The counts and fold changes of R-loop CUT&Tag peak gains (green) and losses (red) are presented for LSH KO cells compared to LSH WT cells. Additionally, the comparison between the WT group and the RNH1-treated group was carried out to confirm the specificity of the detected differential peaks. **H** Pathway analysis of genes exhibiting R-loop gains at the promoter region was conducted, comparing LSH KO and WT cells. The data are representative of n = 3 biologically independent experiments. Source data are provided as a Source Data file.

Subsequently, we employed these cells to generate a genome-wide R- loop map. Our genomic distribution analysis revealed a similar pattern of R- loop signals in both LSH WT and KO PC3 cells (**Figure 3B**). The differences in the genomic distribution of R-loop CUT&Tag peaks were evident on a whole genomic scale, particularly in the promoter regions (**Figure 3C**). The genome browser view revealed areas of increased R-loop accumulation in LSH KO PC3 cells compared to LSH WT cells, based on three biological replicates from three independently derived PC3 cell samples (**Figure 3D**). Samples treated with RNASEH1 (RNH1) enzyme and KO+RNH1 cells served as control groups. These results indicate widespread changes at some genomic locations in the absence of LSH. Notably, the transcriptional start site-related region, rather than the gene body, exhibited a higher R-loop signal intensity in LSH KO PC3 cells (**Figure 3E, Figure S5A**). Further analysis of R-loop signals across different regions revealed that R-loop signals in LSH KO cells were significantly stronger than those in LSH WT cells at the peak regions (**Figure 3F**), indicating substantial R-loop accumulation in PC3 cells lacking LSH. Additionally, we analyzed the count and fold change of R-loop CUT&Tag peak gains (green) and losses (red) between LSH WT and KO PC3 cells. The results indicate that LSH KO cells exhibited a greater number of gains and fewer losses compared to WT cells (**Figure 3G**). Meanwhile, a comparison between LSH WT group and RNH1- treated group was conducted to validate the specificity of the identified differential peaks (**Figure 3G**). Therefore, our CUT&Tag-seq data indicates that LSH can regulate the accumulation of R-loops on a genome-wide scale. Next, to illustrate the molecular pathways of genes regulated by LSH in relation to R-loop gains at the promoter region, a pathway analysis was conducted and presented using a bubble chart that compares LSH KO and WT PC3 cells. The findings revealed that R-loop gains induced by LSH knockout are significantly associated with regions related to DNA-binding transcription activator activity (RNA polymerase II - specific), cell morphogenesis, cell cycle, cell growth, chromosome reorganization, DNA replication, and DNA repair, among others (**Figure 3H**), consistent with previously reported functions of LSH[24–26, 29, 32, 34, 44]. Additionally, we obtained similar results when conducting a pathway analysis of R-loop gains across the entire genome (**Figure S5B**). We further confirmed the regulation of R- loop accumulation by LSH at known R-loop loci and repetitive sequences in PC3 cells, employing CUT-tag qPCR analysis, with SNRPN and EGR1 loci serving as negative controls. Consistent with the CUT-tag sequencing results, we observed that R-loop formation was enhanced following LSH knockout at known R-loop loci and repetitive sequences (**Figure S5C-D**). Furthermore, overexpression of WT RNASEH1 significantly reduced the elevated R-loop levels resulting from LSH knockout (**Figure S5C-D**). Finally, we confirmed that DNA damage is induced by the formation of R-loops in the context of LSH deficiency, as demonstrated by γ-H2AX ChIP-qPCR analysis conducted on the known R-loop loci and repetitive sequences (**Figure S5E-F**).

### 4. LSH interacts with R-loops, and its depletion results in diminished Rad51 filament formation induced by R-loops formation

LSH has been demonstrated to suppress the accumulation of R-loops, reduce DNA damage level, and enhance DNA replication in prostate cancer cells. However, its interaction with R-loop-forming sites has not yet been validated. We established Tet-on LSH-shRNA PC3 cells that overexpress either wild-type or the catalytically inactive human RNASEH1 enzyme (RNASEH1-WT-V5 and RNASEH1-D210N-V5, respectively), as confirmed by Western blot analysis (**Figure 4A**). The RNH1-D210N mutant can recognize and bind to RNA: DNA hybrids; however, unlike the wild-type enzyme, it is unable to cleave the RNA moiety within the hybrid[38, 39]. Overexpression of this mutant slows down R-loop resolution in cells, leading to increased R-loop stability and elevated basal levels[38, 39]. Thus, to investigate the association of LSH with R-loop sites, we first performed a proximity ligation assay (PLA) using a rabbit antibody against LSH and a mouse monoclonal antibody (S9.6) that targets RNA: DNA hybrids in the established cells mentioned above. We observed very few PLA foci in PC3 cells overexpressing WT RNASEH1, irrespective of doxycycline treatment (**Figure 4B**). In contrast, we found a significant increase of PLA foci in PC3 Tet-on LSH-shRNA cells overexpressing D210N RNASEH1 without doxycycline treatment, suggesting that LSH is within the proximity of R-loops (**Figure 4B**). This result was confirmed by the observation that PLA signal staining was significantly reduced in LSH knockdown cells after doxycycline treatment, which ensured the specificity of the PLA staining **(Figure 4B**). Additionally, we assessed the formation of PLA foci in PC3 cells that were transiently transfected with siRNAs targeting RNASEH1 (RNH1) or Senataxin (SETX), another RNA: DNA helicase involved in R-loops resolution[9, 45]. We verified the knockdown of both proteins by western blotting (**Figure 4C**) and observed a subsequent increase in RNA: DNA hybrids through immunostaining with the S9.6 antibody (**Figure S6A**). Next, we performed S9.6/LSH PLA staining in PC3 cells transfected with RNH1 or SETX siRNA. Either RNH1 or SETX deficiency resulted in a significant increase in PLA foci compared to the control cells (**Figure 4D**). Overall, our data confirmed that LSH interacts with R-loops and is associated with R-loop sites formation. To further investigate whether LSH binds to R-loops, we employed R-loop immunoprecipitation (IP) method, which enables the identification of RNA: DNA hybrid-interacting proteins in PC3 cells (**Figure 4E**). LSH was significantly more enriched in RNA: DNA hybrid IP samples compared to the nuclear LAMIN B1 protein and IgG IP samples in PC3 cells overexpressing the catalytically inactive RNH1-D210N variant, relative to RNH1-WT expressing cells (**Figure 4E**). We also aimed to investigate whether LSH can bind to various structures of RNA: DNA hybrids. To achieve this, we purified LSH from insect cells using the previously published method[26] and conducted in-vitro binding assays (**Figure 4F**). We constructed different R-loops and RNA: DNA hybrids with or without 5’-RNA overhangs. Our results demonstrated that the LSH-WT protein exhibits strong binding affinity for all forms of R-loops and RNA: DNA hybrids, as confirmed by the electrophoretic mobility shift assay (EMSA) (**Figure 4G**). Therefore, our findings suggest that LSH binds to R-loops under physiological conditions.

**Figure 4.**
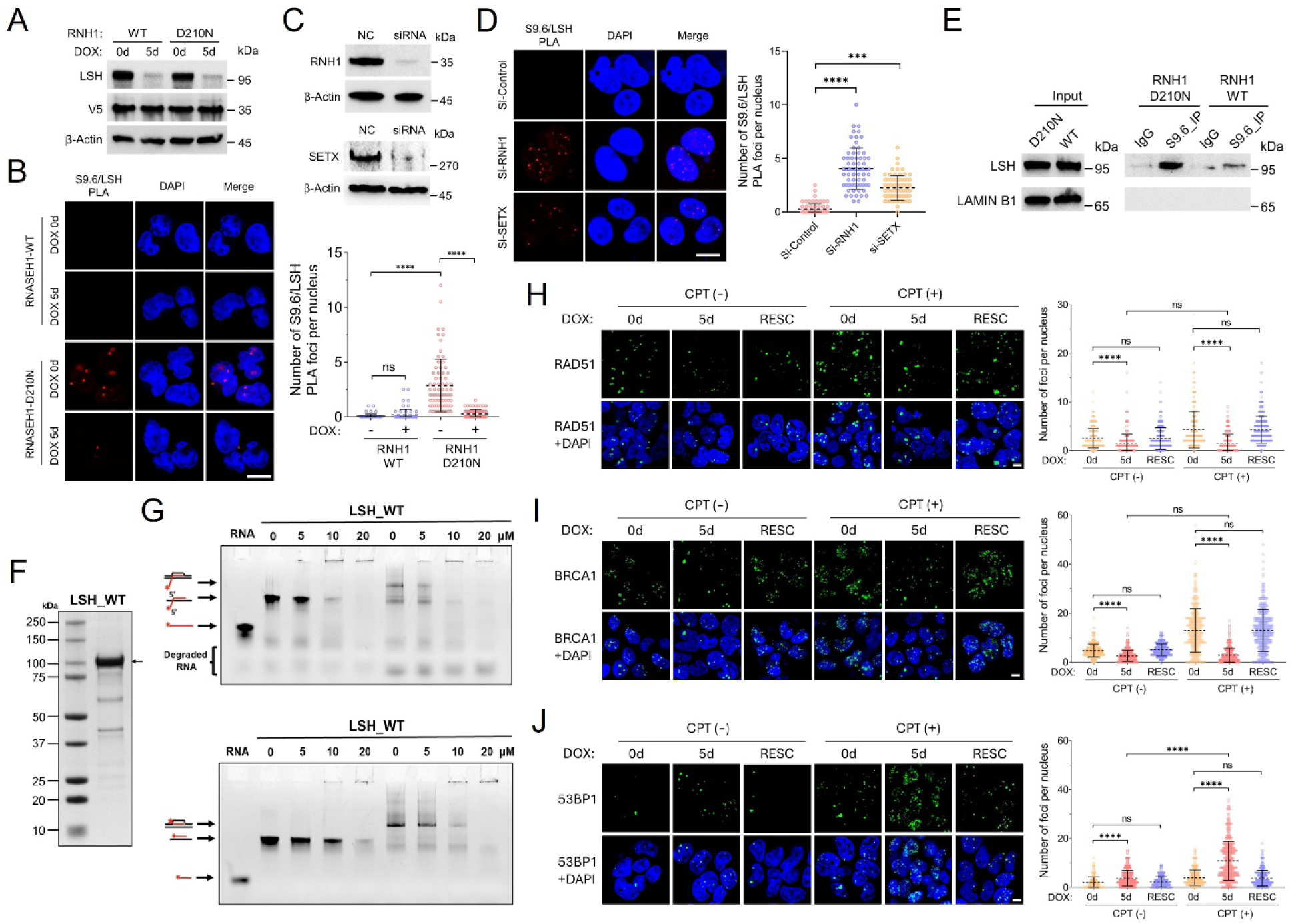
LSH associates with R-loops, and its depletion impairs Rad51 filament formation at R-loop-induced DNA damage. **A** Western blot analysis was conducted on PC3 Tet-on LSH-shRNA cells overexpressing either WT or D210N RNASEH1, both of which were tagged with V5. The cells were left untreated (DOX_0d) or treated with doxycycline for five days (DOX_5d). β-Actin was used as the loading control. **B** Representative images of S9.6/LSH proximity ligation assay (PLA) in DOX_0d and DOX_5d cells overexpressing either WT or D210N RNASEH1. PLA foci (red) indicate the association of the LSH antibody within a 40 nm distance of the S9.6 antibody. The scale bar represents 10 μm. The quantification of PLA foci per nucleus for each experimental condition is presented as a scatter plot with mean ± s.d. (n = 3 biologically independent experiments). ****p < 0.0001; n.s. indicates not significant, as determined by the two-tailed Mann–Whitney test. **C** Western blot analysis was performed to evaluate RNH1 or SETX knockdown in PC3 cells following siRNA transfection, with cells transfected with negative control (NC) siRNA serving as the control group. β-Actin was used as the loading control. **D** Representative images of S9.6/LSH PLA analysis in PC3 cells transfected with siRNA targeting RNH1 or SETX. The scale bar represents 10 μm. The quantification of PLA foci per nucleus for each experimental condition is presented as a scatter plot with mean ± s.d. (n = 3 biologically independent experiments). ****p < 0.0001, as determined by the two-tailed Mann–Whitney test. **E** Western blot analysis of an RNA/DNA hybrid immunoprecipitation (IP) experiment was performed using the S9.6 antibody in PC3 cells overexpressing either WT or D210N RNASEH1. Both input and IP fractions were probed with LSH and lamin B1 antibodies, with LAMIN B1 serving as a negative control. Inputs are displayed on the left, and the S9.6-immunoprecipitated samples are shown on the right. The IgG Lane corresponds to a control IP using an IgG antibody. **F** Coomassie-stained SDS-PAGE gel of purified recombinant LSH_WT protein. **G** Electrophoretic mobility shift assay (EMSA) was conducted to analyze the binding of 0, 5, 10, and 20 μM of purified recombinant LSH_WT protein to 200 nM R-loops or RNA: DNA hybrids, both with and without 5’- RNA overhangs. The RNA 5’-terminus was labeled with 6-FAM fluorescence; DNA is depicted in black, while RNA is shown in red. The samples were loaded onto a 15% polyacrylamide gel and visualized using the Cy2 channel. **H-J** Representative immunofluorescence images depicting RAD51 (**H**), BRCA1 (**I**), and 53BP1 (**J**) foci in PC3 Tet-on LSH-shRNA cells treated with DMSO or with CPT (10 μM for 2 hours). The cells were left untreated (Dox_0d), treated with doxycycline for five days (Dox_5d), or subjected to a five-day doxycycline washout period (RESC). The scale bar represents 10 μm. The quantification of the number of RAD51, BRCA1, and 53BP1 foci per nucleus for each experimental condition is presented as a scatter plot with mean ± s.d. (n = 3 biologically independent experiments). ****p < 0.0001; n.s. indicates not significant, as determined by the two-tailed Mann–Whitney test. Source data are provided as a Source Data file.

Considering that certain chromatin remodelers have been reported to be associated with the resolution of R-loops[9], and given that LSH’s Arabidopsis homolog, DDM1, plays a role in mediating R-loop accumulation[36], we aimed to investigate the potential role of human LSH in removing R-loops and RNA: DNA hybrids in vitro. First, we performed adenosine triphosphate (ATP) hydrolysis assay to evaluate the adenosine triphosphatase (ATPase) activity of recombinant LSH, utilizing either double- stranded DNA (dsDNA) or nucleosomes as substrates (**Figure S7A**), along with the addition of different ATP analogs. The results indicated that the ATPase activity of recombinant LSH was stimulated by both dsDNA and nucleosomes (**Figure S7B**). This stimulation was found to be dependent on ATP hydrolysis, but not on its substitutes, including adenosine diphosphate (ADP), guanosine triphosphate (GTP), and nonhydrolyzable source of ATP (ATP-γ-S) (**Figure S7B**). Moreover, the ATPase activity of recombinant LSH was also stimulated by various R-loops and RNA: DNA hybrids, although this stimulation was lower than that observed with double-stranded DNA (dsDNA) and nucleosomes (**Figure S7C**). Subsequently, we aimed to investigate whether human LSH could unwind R-loops or RNA: DNA hybrids in an ATP- dependent manner across a range of LSH concentrations, from low to high (0 to 20 μM). However, we found no evidence that LSH could unwind R-loops or RNA: DNA hybrids in vitro, regardless of whether the concentration of LSH was low or high (**Figure S7D-E**). This suggests that LSH may interact with other components to regulate R-loop accumulation through either direct or indirect pathways.

LSH deficiency will affect the loading of Rad51, BRCA1 and 53BP1[26], thereby disrupting the balance of DNA repair pathways. Since LSH interacts with R-loops, we aimed to further investigate whether the elevated levels of R-loops in LSH-depleted cells would compromise DNA repair pathways. Camptothecin (CPT) is known to induce R-loop accumulation by inhibiting topoisomerase 1 (TOP1) at active promoters, which subsequently leads to DNA damage[46]. Our results demonstrated that higher levels of S9.6 immunostaining signals were observed in DOX_0d PC3 cells treated with CPT. Furthermore, following doxycycline treatment, LSH-depleted cells (DOX_5d) that were incubated with CPT exhibited an even greater increase in S9.6 immunostaining signals. Notably, these signals could be rescued by washing out doxycycline (RESC) (**Figure S6B**). Simultaneously, the results of γ-H2AX staining revealed changes that corresponded with the accumulation of R-loops (**Figure S6C**). We also verified this finding in another prostate cancer cell line, LNCaP. Similarly, CPT treatment resulted in elevated levels of R-loops and γ-H2AX, with LSH-KD LNCaP cells exhibiting the highest levels of both R-loops and γ-H2AX staining following CPT treatment (**Figure S6D-E**). Subsequently, we investigated the DNA repair pathway checkpoints, including RAD51, BRCA1 and 53BP1 through immunostaining in Tet-on shLSH PC3 cells, both with and without CPT treatment. The results indicated that LSH-deficient cells (DOX_5d) treated with CPT displayed the lowest number of RAD51 and BRCA1 foci per nucleus among the various experimental conditions, reflecting the greatest accumulation of R-loops (**Figure 4H-I**). Conversely, following CPT treatment, the number of 53BP1 foci per nucleus was significantly elevated in the LSH knockdown cells (DOX_5d) compared to control (DOX_0d) and RESC cells (**Figure 4K**). We also observed a similar phenomenon in LNCaP cells: RAD51 and BRCA1 immunostaining signals were the highest in LSH WT cells incubated with CPT (**Figure S6F-G**), while 53BP1 immunostaining signal was the highest in LSH KD cells following CPT treatment (**Figure S6H**). In summary, these results indicate that LSH interacts with R-loops, as demonstrated by our in vivo and in vitro experiments. In the context of R-loop-induced DNA damage, LSH depletion impairs Rad51 filament formation and disrupts DNA repair pathway choice by reciprocally regulating BRCA1 displacement and 53BP1 enrichment at damage sites.

### 5. The ATP binding site is required for LSH-mediated R-loops resolution

The ATP binding site is critical for the proposed chromatin remodeling activity[47, 48]. A point mutation in human LSH protein at this site, which alters a conserved lysine at position 254 to arginine (K254A), as observed in SNF2 factors, completely abolishes its ability to hydrolyze ATP[24, 49] (**Figure 5A**). As a result, LSH protein loses its chromatin remodeling activity. To elucidate the molecular function of the ATP binding domain of LSH protein at regulating R-loops formation, we designed LSH siRNA targeting the noncoding region to silence endogenous LSH expression (**Figure 5B**). Subsequently, we overexpressed exogenous LSH (K254A)-Flag, which carries a point mutation at the ATP binding site, as well as LSH (WT)-Flag, in PC3 cells (**Figure 5B**). Western-blot analysis showed that PC3 cells overexpressing LSH (WT)-Flag and LSH (K254A)-Flag displayed comparable levels of exogenous LSH expression following LSH siRNA treatment, similar to the endogenous LSH expression observed in WT cells (**Figure 5B**). Therefore, we can explore the molecular function of the ATP binding site of LSH in regulating R-loop formation and genomic instability by utilizing these cells.

**Figure 5.**
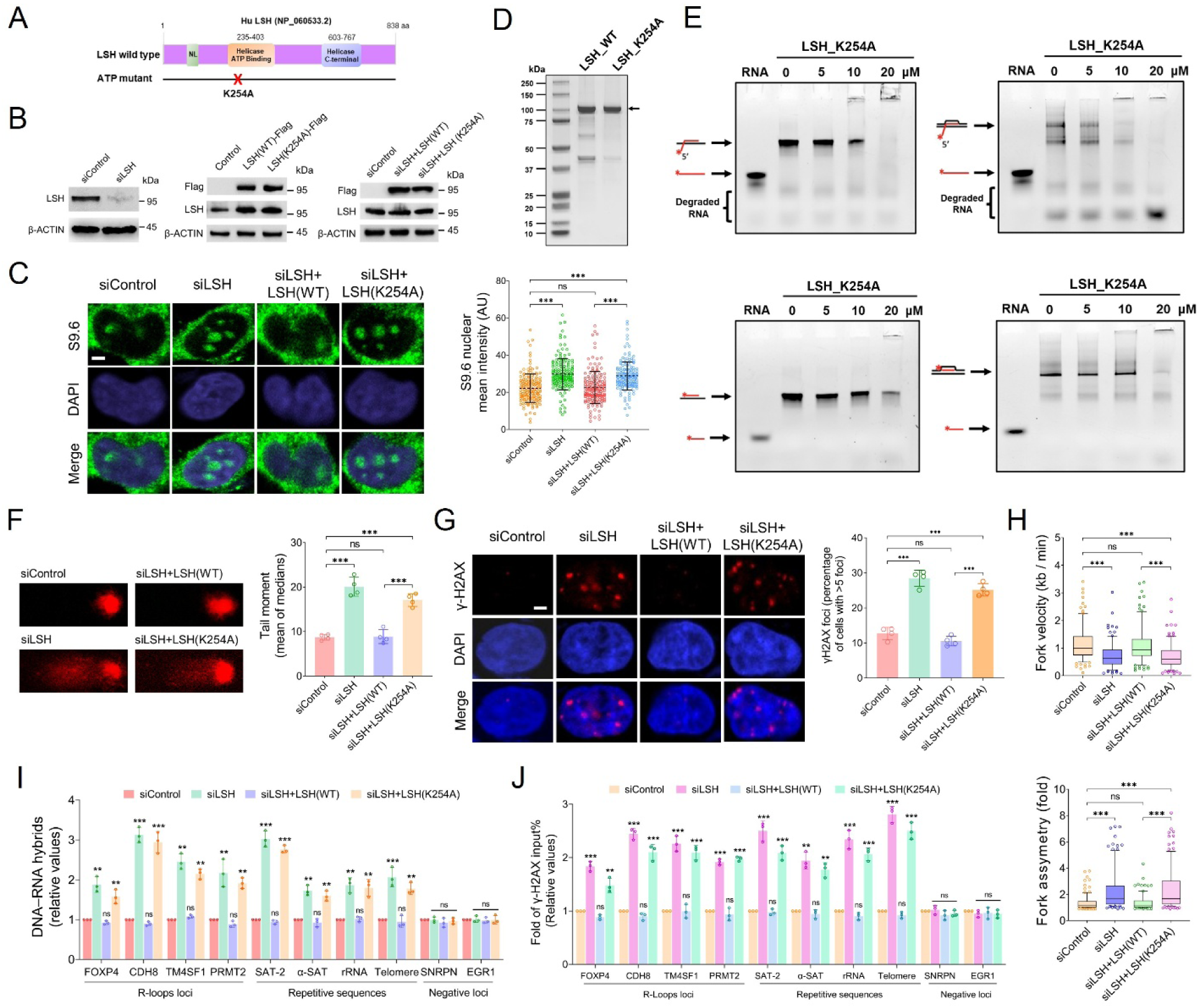
The ATP binding site is essential for LSH-mediated resolution of R-loops. **A** Schematic representation of a human LSH mutation at the ATP binding site, illustrating a point mutation that replaces lysine (K) at amino acid 254 with alanine (A). **B** LSH 3’ UTR siRNA was designed to knock down endogenous LSH expression in PC3 cells (left panel), while these cells overexpressed either exogenous LSH(WT)-Flag or LSH(K254A)-Flag (middle panel). Following transient transfection with LSH 3’ UTR siRNA, western blot analysis (right panel) of PC3 cells indicated that the expression of exogenous LSH(WT)-Flag or LSH(K254A)-Flag protein was comparable to endogenous LSH expression, with β-Actin serving as the loading control. **C** Representative images of S9.6 immunostaining were used to evaluate the R-loop levels in PC3 cells treated as described in panel B. The scale bar represents 5 μm. Data are presented as a scatter plot with mean ± s.d. (*n* = 3 biologically independent experiments). ***p < 0.001; n.s. indicates not significant, as determined by the two-tailed Mann–Whitney test. **D** Coomassie-stained SDS-PAGE gel of purified recombinant LSH_WT and LSH_K254A proteins. **E** Electrophoretic mobility shift assays (EMSA) were conducted to analyze the binding of purified mutant LSH_K254A recombinant protein at concentrations of 0, 5, 10, and 20 μM to 200 nM R- loops or RNA: DNA hybrids, with or without 5’-RNA overhangs. The RNA 5’- terminus was labeled with 6-FAM fluorescence, with DNA represented in black and RNA in red. The samples were loaded onto a 15% polyacrylamide gel and visualized using the Cy2 channel. **F** Representative images of the alkaline comet assay along with quantification of tail moment in PC3 cells treated as outlined in panel B. Data are presented as a bar graph with mean ± s.d. (*n* = 3 biologically independent experiments). ***p < 0.001; n.s. indicates not significant, as determined by the two-tailed Mann–Whitney test. **G** Representative images of γ-H2AX immunostaining, complemented by a bar graph, illustrate the percentage of PC3 cells that contain more than five γ-H2AX foci, as treated in panel B. The scale bar represents 5 μm. Data are presented as a bar graph with mean ± s.d. (*n* = 3 biologically independent experiments). ***p < 0.001; n.s. indicates not significant, as determined by the two-tailed Mann–Whitney test. **H** Quantification of replication fork velocity and asymmetry in PC3 cells treated as described in panel B. The data are presented as a Tukey-style box plot, where the center line represents the median value, and the boxes and whiskers indicate the 25th to 75th percentiles and the 5th to 95th percentiles, respectively. Outliers are represented as individual points. The data are representative of n = 3 biologically independent experiments. ***p < 0.001; n.s. indicates not significant, as determined by the two-tailed Mann–Whitney test. **I** DNA-RNA hybrids CUT&Tag-qPCR analysis at R-loop loci and repetitive sequences in PC3 cells treated as described in panel B. Signal values were normalized to the controls and plotted as mean ± s.d. (n = 3 biologically independent experiments). **p < 0.01; ***p < 0.001; n.s. indicates not significant, as determined by the One-way ANOVA with Tukey’s multiple comparison test. **J** ChIP–qPCR analysis of γ-H2AX at R-loops loci and repetitive sequences in PC3 cells treated as described in panel B. Signal values were normalized to the controls and plotted as mean ± s.d. (n = 3 biologically independent experiments). **p < 0.01; ***p < 0.001; n.s. indicates not significant, as determined by the two-tailed Mann–Whitney test. Source data are provided as a Source Data file.

First, we investigated whether LSH-regulated R-loop formation depends on the ATP domain in PC3 cells. Our results revealed that the nuclear S9.6 signal was significantly increased in cells treated with LSH siRNA. This signal returned to control levels in the siLSH+LSH (WT) cells, but not in the siLSH+LSH (K254A) cells (**Figure 5C**). Additionally, we generated LSH (K254A) recombinant protein to evaluate the binding activity of the ATP mutant LSH (K254A) through EMSA assays (**Figure 5D**). In contrast to the LSH (WT) protein, the LSH (K254A) protein exhibited no ATPase activity in the presence of dsDNA, nucleosomes, R-loops, or RNA: DNA hybrids (**Figure S8A-B**). Our results of EMSA assays demonstrated that LSH (K254A) protein, which has a mutation at the ATP binding domain, also exhibited a high level of binding activity with various R-loops or RNA: DNA hybrids (**Figure 5E**). Moreover, we investigated whether LSH (K254A) mutation leads to an accumulation of R-loops in conjunction with genomic instability. The alkaline comet assay and γH2AX immunostaining revealed that the level of DNA damage was significantly higher in cells treated with LSH siRNA compared to the control cells, and this increase in DNA damage was mitigated in the siLSH+LSH (WT) cells (**Figure 5F-G**). In contrast, the siLSH+LSH (K254A) cells continued to exhibit significantly higher DNA damage (**Figure 5F-G**). In addition, we performed DNA fiber analysis to monitor the progression of DNA replication. Our findings revealed a significant reduction in replication fork velocity, along with an increase in replication fork asymmetry, in PC3 cells treated with LSH siRNA (**Figure 5H**). These replication defects were effectively rescued in the siLSH+LSH (WT) cells, but not in the siLSH+LSH (K254A) cells (**Figure 5H**). We also employed more sensitive techniques, specifically CUT-tag qPCR and ChIP- qPCR, to further validate our findings by evaluating the levels of R-loops and γ-H2AX. The results confirmed our previous findings, demonstrating a significant increase in both R-loop and γ-H2AX levels at known R-loop loci and repetitive sequences in cells treated with LSH siRNA (**Figure 5I-J**). These levels were significantly decreased in the siLSH+LSH (WT) cells, whereas they remained obviously elevated in the siLSH+LSH (K254A) cells (**Figure 5I-J**). Finally, the results from the CCK-8 assay (**Figure S8C**), doubling time measurements (**Figure S8D**), and TUNEL assays (**Figure S8E-F**) indicated that the ATP binding domain of LSH is essential for cell viability and apoptosis in PC3 cells.

In summary, our experimental data underscores the pivotal role of the ATP binding domain of LSH in the accumulation of R-loop, which is closely associated with the maintenance of genomic stability and the regulation of cell proliferation and apoptosis in prostate cancer cells.

### 6. LSH resolves R-loops-mediated transcription–replication conflicts through FANCD2 at stalled replication forks

R-loop homeostasis is maintained through three primary mechanisms involving proteins that prevent nascent RNA from re-hybridizing with DNA (such as the RNA-binding factors THOC1 and UAP56), proteins that resolve R-loops (including RNASEH1 and SETX), and DNA repair factors from the Fanconi anemia (FA) pathway[21, 50]. To investigate the mechanism by which LSH regulates R-loop homeostasis, we performed double depletion of LSH along with a representative factor from each of these mechanisms (UAP56, THOC1, FANCD2, and SETX) in PC3 cells. The efficiency of the depletion was confirmed through Western blot analysis (**Figure S9A**). We then assessed R-loop levels and DNA damage using IF staining with S9.6 and γH2AX antibodies, respectively. As anticipated, single depletion of either factor led to an increase in R-loops and DNA breaks (**Figure 6A** and **Figure S9B**). Co-depletion of LSH along with THOC1, UAP56, and SETX resulted in a further elevation of both R-loops and DNA breaks (**Figure 6A** and **Figure S9B**). In contrast, co-depletion of LSH with FANCD2 did not result in significant changes in R-loops and DNA breaks compared to the single depletions (**Figure 6A** and **Figure S9B**). This suggests that LSH operates within the same mechanistic framework as FANCD2 to prevent the accumulation of R-loops and the associated DNA damage in PC3 cells.

**Figure 6.**
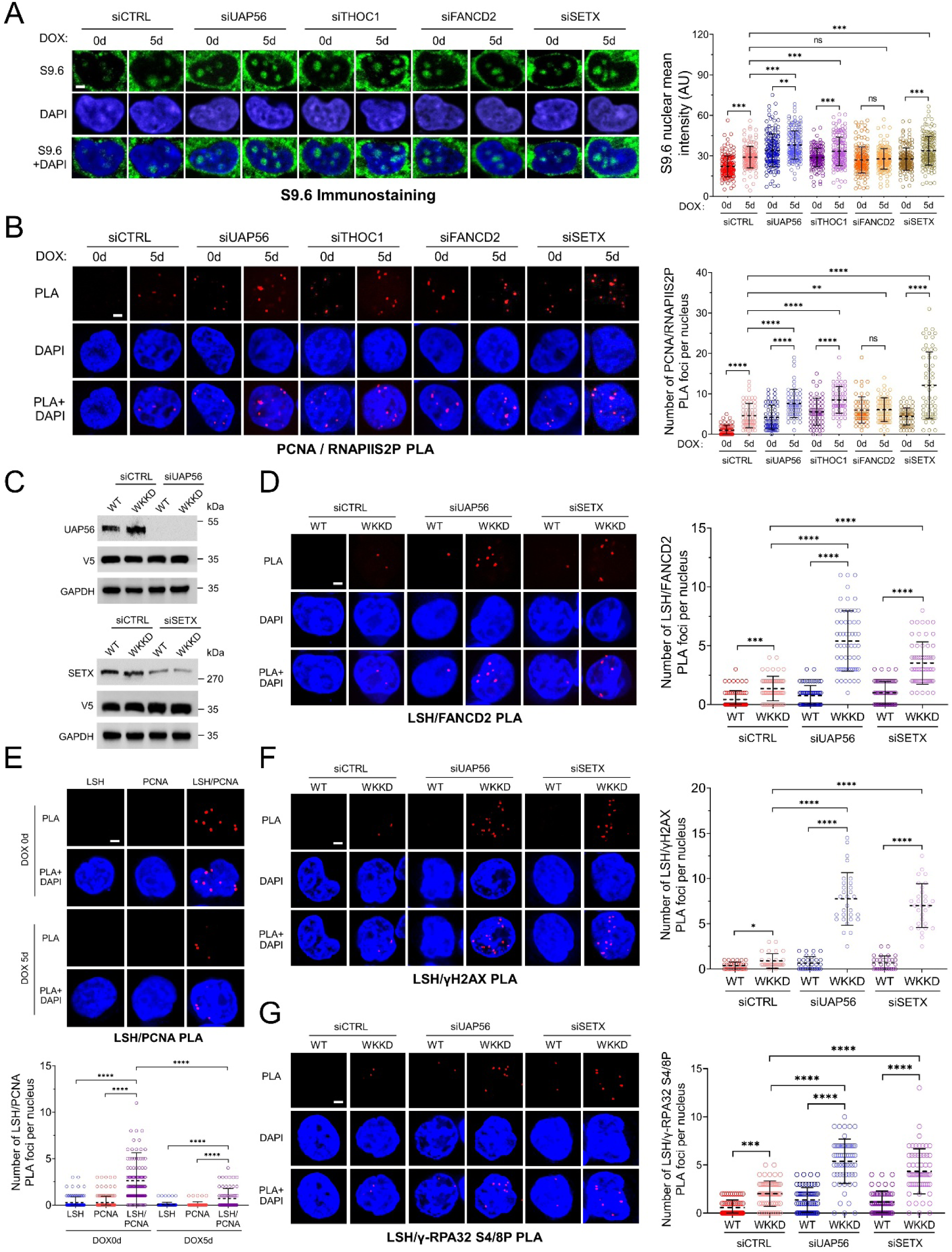
LSH interacts with FANCD2 to resolve transcription- replication conflicts mediated by R-loops at stalled replication forks. A Representative images of S9.6 immunostaining to evaluate R-loop levels in PC3 Tet-on LSH-shRNA cells following transient transfection with siRNAs targeting UAP56, THOC1, FANCD2, or SETX. The cells were left untreated (DOX_0d) or treated with doxycycline for five days (DOX_5d). The scale bar represents 5 μm. Signal intensity data are depicted as a scatter plot with mean ± s.d. (n = 3 biologically independent experiments). ** p < 0.01; ***p < 0.001; n.s. indicates not significant, as determined by the two-tailed Mann– Whitney test. AU denotes arbitrary units. **B** Representative images of PCNA/RNAPIIS2P PLA in cells treated as described in panel A. The scale bar represents 5 μm. Quantification of the number of PLA foci per nucleus for each experimental condition is presented as a scatter plot with mean ± s.d. (n = 3 biologically independent experiments). ** p < 0.01; ****p < 0.0001; n.s. indicates not significant, as determined by the two-tailed Mann–Whitney test. **C** Western blot analysis was performed in PC3 Tet-on LSH-shRNA cells overexpressing either wild-type (WT) or WKKD RNASEH1, following transient transfection with siRNAs targeting UAP56 or SETX. GAPDH was used as the loading control. **D** Representative images of LSH/FANCD2 PLA in cells treated as described in panel C. The scale bar represents 5 μm. Quantification of the number of PLA foci per nucleus for each experimental condition is presented as a scatter plot with mean ± s.d. (n = 3 biologically independent experiments). ***p < 0.001; ****p < 0.0001, as determined by the two-tailed Mann–Whitney test. **E** Representative images of LSH/PCNA PLA in PC3 Tet-on LSH-shRNA cells, which were left untreated (DOX_0d) or treated with doxycycline for five days (DOX_5d). Negative controls, utilizing only LSH or PCNA antibodies, were performed to verify the specificity of the detected PLA signals. The scale bar represents 5 μm. Quantification of the number of PLA foci per nucleus for each experimental condition is presented as a scatter plot with mean ± s.d. (n = 3 biologically independent experiments). ****p < 0.0001, as determined by the two-tailed Mann–Whitney test. **F** Representative images of LSH/γH2AX PLA in cells treated as described in panel C. The scale bar represents 5 μm. Quantification of the number of PLA foci per nucleus for each experimental condition is presented as a scatter plot with mean ± s.d. (n = 3 biologically independent experiments). *p < 0.05; ****p < 0.0001, as determined by the two-tailed Mann–Whitney test. **G** Representative images of LSH/γ-RPA32 S4/S8P PLA in cells treated as described in panel C. The scale bar represents 5 μm. Quantification of the number of PLA foci per nucleus for each experimental condition is presented as a scatter plot with mean ± s.d. (n = 3 biologically independent experiments). ***p < 0.001; ****p < 0.0001, as determined by the two-tailed Mann–Whitney test. Source data are provided as a Source Data file.

Considering that the increase of R-loops in cells primarily occurs during T-R conflicts, and given LSH has close association with transcription and replication fork progression (**Figure 1H-J**, **Figure 2G-I**, **Figure 5H**, **Figure S3G-I**, and **Figure S4F**), we aimed to investigate whether LSH’s role in R-loop protection is linked to its potential function in managing T-R conflicts in PC3 cells. Additionally, we sought to determine whether LSH functionally interacts with FANCD2, which serves as a biomarker for replication fork stalls[41–43]. Firstly, to evaluate T-R conflicts, we conducted PLA experiments using proliferating cell nuclear antigen (PCNA), a key component of the replisome for identifying replication sites[51, 52], alongside the elongating form of RNA polymerase II phosphorylated at Ser2 (RNAPII-S2P)[53, 54]. Our results indicated that the depletion of LSH substantially elevated the number of PCNA/RNAPII-S2P PLA foci (**Figure 6B**). Additionally, we observed a synergistic increase in PLA foci when LSH was co-depleted with UAP56, THOC1, or SETX, but not when co-depleted with FANCD2 (**Figure 6B**). These findings suggest that LSH plays a role in resolving R-loop-mediated T-R conflicts together with FANCD2 in PC3 cells. Secondly, considering that LSH and FANCD2 are implicated in the same pathway to prevent R-loop formation and resolve T-R conflicts (**Figure 6A-B** and **Figure S6B**), we proceeded to evaluate the colocalization of LSH with FANCD2 in an R-loop- dependent manner through PLA using anti-LSH and anti-FANCD2 antibodies. Thus, we utilized siRNA targeting UAP56 and SETX to trigger R-loop formation (**Figure 6A** and **Figure S6A**) and employed PC3 cells overexpressing RNASEH1-WT or RNASEH1-WKKD to confirm that the LSH/FANCD2 interaction was R-loop accumulation dependent. The efficiency of siRNA in knocking down UAP56 and SETX in the RNASEH1-V5 overexpressing cells was confirmed through Western-blot analysis (**Figure 6C**). The LSH/FANCD2 PLA-positive foci were notably enriched in PC3 cells (**Figure 6D**). Specifically, the number of positive PLA foci per cell was significantly increased when R-loop-mediated T-R conflicts were induced by the depletion of UAP56 or SETX in the RNASEH1-WKKD overexpressing cells, which lack R-loop binding and catalytic functions, thus serving as negative control cells (**Figure 6D**). In contrast, the number of LSH/FANCD2 PLA foci was significantly reduced in the RNASEH1-WT overexpressing cells (**Figure 6D**). Therefore, these findings suggest that LSH and FANCD2 are closely interacting with each other in a manner dependent on R-loop formation.

Finally, considering previous findings that R-loop-mediated replication fork stalling is caused by LSH depletion, we wondered whether LSH is enriched at replication sites. To investigate this, we performed PLA between LSH and PCNA. Indeed, the application of anti-LSH and anti-PCNA antibodies produced nuclear PLA foci (**Figure 6E**), reinforcing the conclusion that LSH is enriched at sites of replication fork stalling. Furthermore, to assess whether the association between LSH and FANCD2 reflects replication fork blocks that could lead to double-strand breaks (DSBs), we performed PLA between LSH and markers of either DNA breakage (γH2AX)[55] or replication fork stalling (RPA phosphorylated at Ser4/8, RPA- S4/8P)[56]. Consistently, we observed positive PLA foci for both LSH/γH2AX and LSH/RPA32-S4/8P interactions in PC3 cells (**Figure 6F-G**). Notably, the PLA foci were significantly increased in PC3 cells with UAP56 or SETX depletion upon overexpression of RNASEH1-WKKD, and they disappeared when RNASEH1-WT was overexpressed, thereby confirming the specificity of the PLA signal (**Figure 6F-G**). Therefore, these findings indicate that LSH plays a crucial role in resolving R-loop-mediated T-R conflicts and in addressing DNA damage via FANCD2 at stalled replication forks.

### 7. LSH-mediated R-loops correlate with MYC and E2F target genes expression by modulating RNA Pol II occupancy at promoter regions

Previous studies have highlighted the role of LSH in genome maintenance; however, the significance of LSH’s remodeling properties in regulating gene expression in cancer remains unclear. It is known that the aberrant accumulation of R-loops leads to the stacking of RNAPII during elongation, which is a primary source of genomic instability[9, 10, 50]. Therefore, we hypothesized that LSH may regulate gene transcription by promoting RNAPII elongation through the alleviation of R-loop accumulation, thereby reducing potential obstacles to the transcription machinery. We conducted a transcriptomic analysis using bulk RNA sequencing in LSH WT, LSH KO, and KO+RNH1 PC3 cells. Principal component analysis (PCA) and unbiased clustering analysis revealed distinct expression patterns among the different cell types (**Figure S10A-B**). The RNA-seq results indicate that 471 overlapping genes are downregulated in LSH KO cells compared to LSH WT cells; however, these genes are restored in KO+RNH1 cells (**Figure 7A**). This suggests that these genes are regulated by the effects of LSH on gene transcription through the modulation of R-loop formation. Furthermore, we performed pathway analysis of the overlapping genes, with a primary focus on cancer signaling pathways, transcriptional activity, cellular biological processes (**Figure 7B**), as well as cell development and differentiation, cell metabolism, and cell structure (**Figure S7C**). The heat maps display the changes in relative expression levels of the overlapping genes in LSH WT, LSH KO, and KO+RNH1 PC3 cells, as outlined in the pathway analysis (**Figure 7C**, **Figure S10D**). The R-loop CUT&Tag-seq analysis revealed that these overlapping genes also displayed a significantly higher R-loop signal intensity at the transcriptional start site region, rather than within the gene body, in LSH KO PC3 cells compared to both LSH WT and KO+RNH1 PC3 cells (**Figure 7D**, **Figure S10E**). Samples treated with the RNH1 enzyme served as a negative control to confirm the specificity of the detected R-loop signals (**Figure 7D**, **Figure S10E**). Based on these findings, we have concluded that LSH may regulate the transcription of target genes by alleviating the accumulation of R-loops.

**Figure 7.**
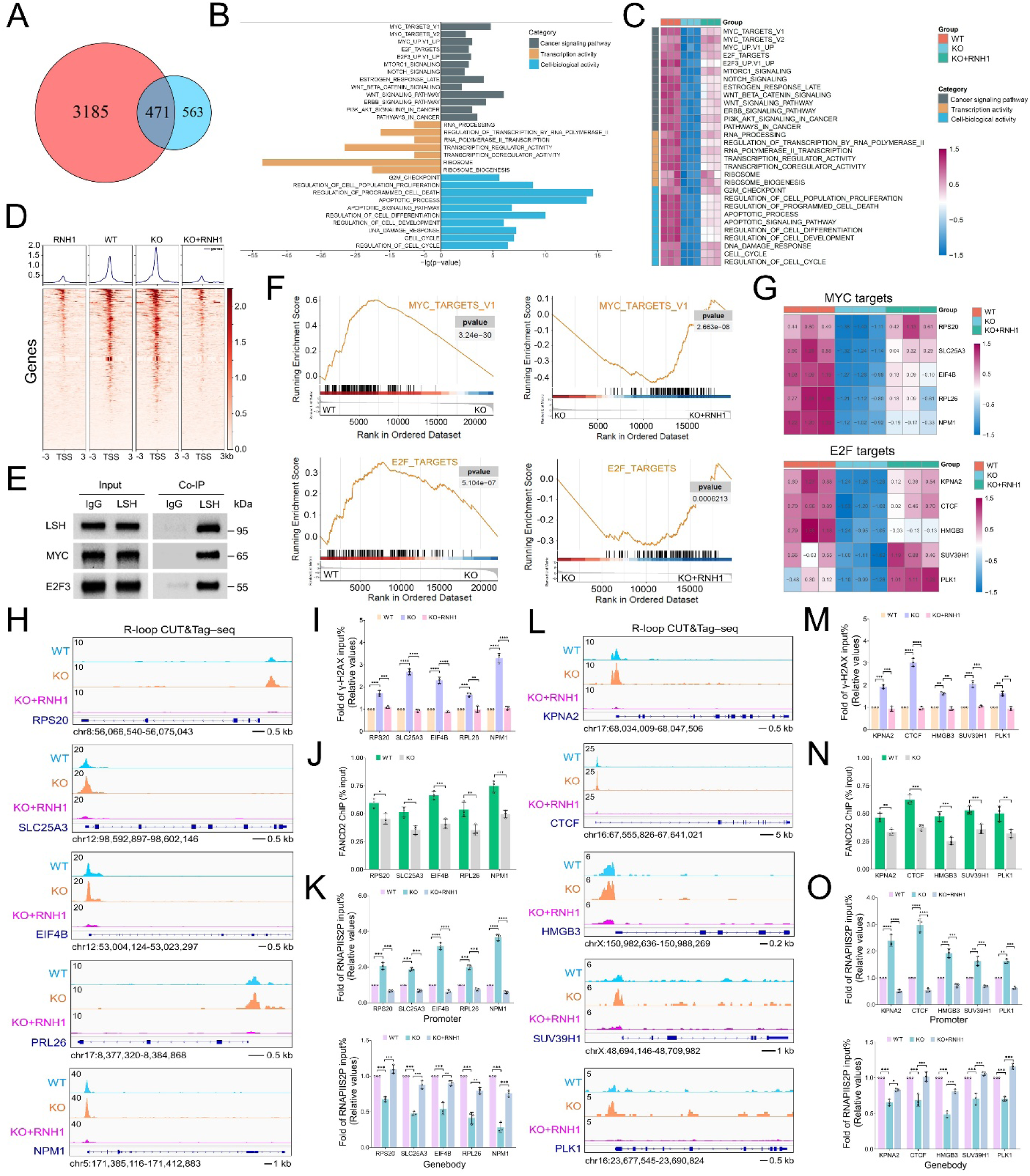
LSH-mediated R-loop accumulation is linked to MYC and E2F target genes expression through the modulation of RNA Polymerase II occupancy at promoter regions. **A** The Venn diagram illustrates the number of overlapping genes (n = 471) that are downregulated in LSH KO cells (KO vs. WT, red) and restored in KO+RNH1 cells (blue). Data are representative of n = 3 biologically independent experiments. **B** Pathway analysis of the overlapping genes was conducted, with a primary focus on cancer signaling pathways, transcriptional activity, and cellular biological processes. Data are representative of n = 3 biologically independent experiments. **C** Heat map displaying the relative expression levels of overlapping genes in WT, KO, and KO+RNH1 PC3 cells, as outlined in the pathway analysis from panel B. Data are representative of n = 3 biologically independent experiments. **D** The average R-loop CUT&Tag signals over 6 kb regions centered on the TSS of overlapping genes were analyzed in WT, KO, and KO+RNH1 PC3 cells. Samples treated with the RNH1 enzyme served as a negative control to confirm the specificity of the detected R-loop signals. Data are representative of n = 3 biologically independent experiments. TSS denotes transcription start site. **E** Co-IP experiments were performed in PC3 cells to evaluate the interaction between LSH and MYC as well as E2F3 proteins. Whole-cell lysates were collected and subjected to immunoprecipitation using either an anti-LSH antibody or an IgG control. Immunoblotting was subsequently carried out with anti-MYC and anti-E2F3 antibodies. Input samples are indicated, and the data represent n = 3 biologically independent experiments. **F** Enrichment plots for MYC and E2F targets were generated to compare WT and KO cells and further validated by the comparison of KO cells with KO+RNH1 cells. Data are representative of n = 3 biologically independent experiments. **G** The heat map illustrates the relative expression levels of five selected MYC and E2F target genes among the overlapping genes in WT, KO, and KO+RNH1 PC3 cells. Data are representative of n = 3 biologically independent experiments. **H** Genome browser snapshots of CUT&Tag-seq data depict R-loop accumulation at the selected MYC target genes, as illustrated in panel G, for WT, KO, and KO+RNH1 PC3 cells. **I** ChIP–qPCR analysis of γ-H2AX at the selected MYC target genes, as illustrated in panel G, for WT, KO, and KO+RNH1 PC3 cells. Values were normalized to the WT group and presented as a bar graph with mean ± s.d. (n = 3 biologically independent experiments). **p < 0.01; ***p < 0.001; ****p < 0.0001, as determined by the One-way ANOVA with Tukey’s multiple comparison test. **J** ChIP–qPCR analysis of FANCD2 at the selected MYC target genes, as shown in panel G, for WT and KO PC3 cells. Data are presented as a bar graph with mean ± s.d. (n = 3 biologically independent experiments). *p < 0.05; **p < 0.01; ***p < 0.001, as determined by the two- tailed Mann–Whitney test. **K** ChIP–qPCR analysis of RNAPIIS2P at the promoter and gene body regions of the selected MYC target genes, as shown in panel G, for WT, KO, and KO+RNH1 PC3 cells. Values were normalized to the WT group and presented as a bar graph with mean ± s.d. (n = 3 biologically independent experiments). **p < 0.01; ***p < 0.001; ****p < 0.0001, as determined by the One-way ANOVA with Tukey’s multiple comparison test. **L** Genome browser snapshots of CUT&Tag-seq data depict R-loop accumulation at the selected E2F target genes, as illustrated in panel G, for WT, KO, and KO+RNH1 PC3 cells. **M** ChIP–qPCR analysis of γ-H2AX at the selected E2F target genes, as illustrated in panel G, for WT, KO, and KO+RNH1 PC3 cells. Values were normalized to the WT group and presented as a bar graph with mean ± s.d. (n = 3 biologically independent experiments). **p < 0.01; ***p < 0.001; ****p < 0.0001, as determined by the One-way ANOVA with Tukey’s multiple comparison test. **N** ChIP–qPCR analysis of FANCD2 at the selected E2F target genes, as shown in panel G, for WT and KO PC3 cells. Data are presented as a bar graph with mean ±s.d. (n = 3 biologically independent experiments). **p < 0.01; ***p < 0.001, as determined by the two-tailed Mann–Whitney test. **O** ChIP–qPCR analysis of RNAPIIS2P at the promoter and gene body regions of the selected E2F target genes, as shown in panel G, for WT, KO, and KO+RNH1 PC3 cells. Values were normalized to the WT group and presented as a bar graph with mean ± s.d. (n = 3 biologically independent experiments). *p < 0.05; **p < 0.01; ***p < 0.001; ****p < 0.0001, as determined by the One-way ANOVA with Tukey’s multiple comparison test. Source data are provided as a Source Data file.

It has been reported that LSH interacts with the key oncogenic transcription factors MYC and E2F to regulate the expression of target genes that are critical for the proliferation and migration of cancer cells[35, 57]. In our research, we have also found that LSH plays a regulatory role in the expression of MYC and E2F target genes. Therefore, we aim to investigate the underlying molecular mechanisms. Our Co-IP experiments confirmed that LSH exhibits strong interactions with the MYC and E2F3 proteins, which aligns with previously reported findings[35, 57] (**Figure 7E**). Enrichment plots confirmed that MYC and E2F target genes were markedly enriched in LSH WT cells compared to LSH KO cells. This enrichment was effectively restored in KO+RNH1 cells through the overexpression of RNASEH1 (**Figure 7F**). Subsequently, we selected five overlapping MYC and E2F target genes to assess R-loop accumulation and RNAPII enrichment during transcriptional elongation. In line with the previous results, the heat map showed that the expression of these selected genes was significantly reduced in LSH KO PC3 cells, whereas it was restored in KO+RNH1 cells (**Figure 7G**). This observation was further validated by RT-qPCR (**Figure S10F**). Additionally, the expression changes of the selected MYC and E2F target genes were confirmed in LSH WT, LSH KD, and KD+RNH1 LNCaP cells (**Figure S11A**, **Figure S11H**). It has been reported that LSH interacts with MYC and E2F proteins to regulate the expression of target genes at their promoter regions[35]. We aimed to detect the enrichment of LSH, MYC, and E2F at the promoter regions of the selected overlapping genes. Our results indicated that LSH enrichment was significantly lower in LSH KO PC3 cells compared to LSH WT cells (**Figure S10G**). Consistent with previously reported findings, our results showed that the enrichment of MYC and E2F at the promoter regions was obviously decreased in LSH_KO PC3 cells (**Figure S10H**). Moreover, we found that R-loop accumulation was significantly higher at these promoter regions in LSH KO PC3 cells as detected by R-loop CUT&Tag-seq (**Figure 7H**, **Figure 7L**). This finding was validated by R-loop CUT&Tag-qPCR (**Figure S10I**). Furthermore, in line with the previous findings, we observed that the level of γH2AX was significantly higher, while the level of FANCD2 was lower at the promoter regions in LSH KO PC3 cells relative to LSH WT cells (**Figure 7I-J**, **Figure 7M-N**). Notably, the elevated γH2AX levels were normalized in KO+RNH1 cells (**Figure 7I**, **Figure 7M**). Similarly, at the promoter regions of the selected genes, we observed coordinated changes in the enrichments of LSH, MYC, and E2F, along with R-loop accumulation and variations in γH2AX and FANCD2 levels in LSH WT, LSH KD, and KD+RNH1 LNCaP cells (**Figure S11B-F**, **Figure S11I-M**). Finally, we aimed to investigate whether LSH deficiency-induced R- loop accumulation and genomic instability affect RNAPII enrichment during transcriptional elongation. Using CUT&Tag-qPCR, we found that the levels of phosphorylated serine 2 RNA polymerase II (RNAPIIS2P), a hallmark of active transcriptional elongation[58], were significantly higher at the promoter regions of the selected MYC and E2F target genes in LSH_KO PC3 cells compared to LSH_WT cells. Conversely, RNAPIIS2P levels were markedly reduced at the gene body regions (**Figure 7K**, **Figure 7O**). Intriguingly, in KO+RNH1 PC3 cells, RNAPIIS2P levels were effectively restored to levels comparable to those in wild-type cells in both the promoter and gene body regions (**Figure 7K**, **Figure 7O**). These findings were further corroborated in LSH_WT, LSH_KD, and KD+RNH1 LNCaP cells (**Figure S11G**, **Figure S11N**). Therefore, our findings suggest that LSH can promote RNAPII elongation by alleviating R-loop accumulation at the promoter regions of MYC and E2F target genes, thereby reducing potential obstacles to the transcription machinery in the prostate cancer cells.

### 8. Clinical relevance of LSH expression and R-loops accumulation in prostate cancer

We have identified LSH as a critical factor mediating R-loop accumulation, DNA damage, and the modulation of proliferation and apoptotic pathways in prostate cancer cells. Consequently, we examined the clinical relevance of LSH expression in prostate cancer (PCa), with a particular focus on its association with R-loop accumulation, DNA damage response, and patient outcomes.

Firstly, using bulk RNA-seq data from The Cancer Genome Atlas - Prostate Adenocarcinoma (TCGA-PRAD), we evaluated LSH expressions in subgroups of prostate cancer patients categorized by Gleason score, clinical stage, and lymph node metastasis. Our results indicated that LSH expression was significantly higher in prostate cancer tissues compared to normal tissues and showed a positive correlation with elevated Gleason scores, advanced pathological stages, and increased prostate-specific antigen (PSA) levels (**Figure 8A, Figure S12A**). Kaplan-Meier analysis demonstrated that high LSH expression was significantly associated with shorter overall survival (OS), progression-free interval (PFI), and disease-specific survival (DSS), underscoring its critical role in the progression of prostate cancer (**Figure 8B, Figure S12B**). Next, we computed the R-loop and DNA damage repair (DDR) scores to evaluate the potential role of LSH in regulating R-loop accumulation, DNA damage, and genomic instability within the prostate cancer database. In detail, a higher R-loop score indicates a greater enrichment of R-loop formation[59], while a higher DDR score signifies a stronger capacity for DNA damage response[60, 61]. Overall, the data presented in the columns indicate that higher levels of LSH are significantly associated with lower R-loop score (RS) and higher DDR scores (DS) (**Figure 8C**). This finding is consistent with the Gene Set Enrichment Analysis (GSEA), which revealed that R-loop regulatory genes and DDR related genes were significantly enriched in patients with higher LSH levels (**Figure 8D**). Furthermore, LSH levels are closely associated with various clinicopathological features in prostate cancer patients. Specifically, LSH levels exhibit a positive correlation with the Gleason score (GS), tumor purity (TP), pathological T stage (pT), and pathological N stage (pN) (**Figure 8C**). This suggests that LSH may play a crucial role in the DNA damage response and the regulation of R-loop enrichment during the progression of prostate cancer. We further performed a Pearson correlation analysis among LSH levels, R-loop scores, and DDR scores using the prostate cancer database. The results revealed that LSH levels are negatively correlated with R-loop scores and positively correlated with DDR scores (**Figure 8E**). Furthermore, R-loop scores also exhibit a negative correlation with DDR scores (**Figure 8E**). Patients were categorized into low and high groups based on LSH levels, R-loop scores, and Gleason scores. Consistent with the previous findings, we observed that patients in the high LSH group had lower R-loop scores and higher DDR scores, while those in the high R-loop score group exhibited lower DDR scores (**Figure 8F**). Meanwhile, patients with advanced prostate cancer (high Gleason score > 7) demonstrated lower R-loop scores and higher DDR scores (**Figure S12C**). Moreover, single-cell RNA sequencing (scRNA-seq) data revealed a heterogeneous distribution of LSH levels and R-loop scores in prostate cancer cells, highlighting distinct clustering patterns specific to LSH and R-loops (**Figure 8G**). Notably, a subset of tumor cells with the highest LSH expression formed an independent cluster and exhibited the lowest corresponding R-loop scores (**Figure 8G**). To further validate these correlations not only in prostate tumor tissue but also in prostate cancer cell lines, we used LSH WT, LSH knockout (LSH_KO) and LSH_KO with RNH1 overexpression (KO+RNH1) PC3 cell lines. The results suggest that the knockout of LSH results in increased R-loop scores, which can be partially restored through the overexpression of RNH1 (**Figure S12D**). Additionally, LSH_KO cells exhibited reduced enrichment of R-loop regulatory genes compared to LSH_WT cells, while the overexpression of RNH1 was able to partially restore the enrichment of these genes (**Figure S12D**). These findings highlight the potential regulatory role of LSH in modulating R-loops formation and the mechanisms of DNA damage response in prostate cancer.

**Figure 8.**
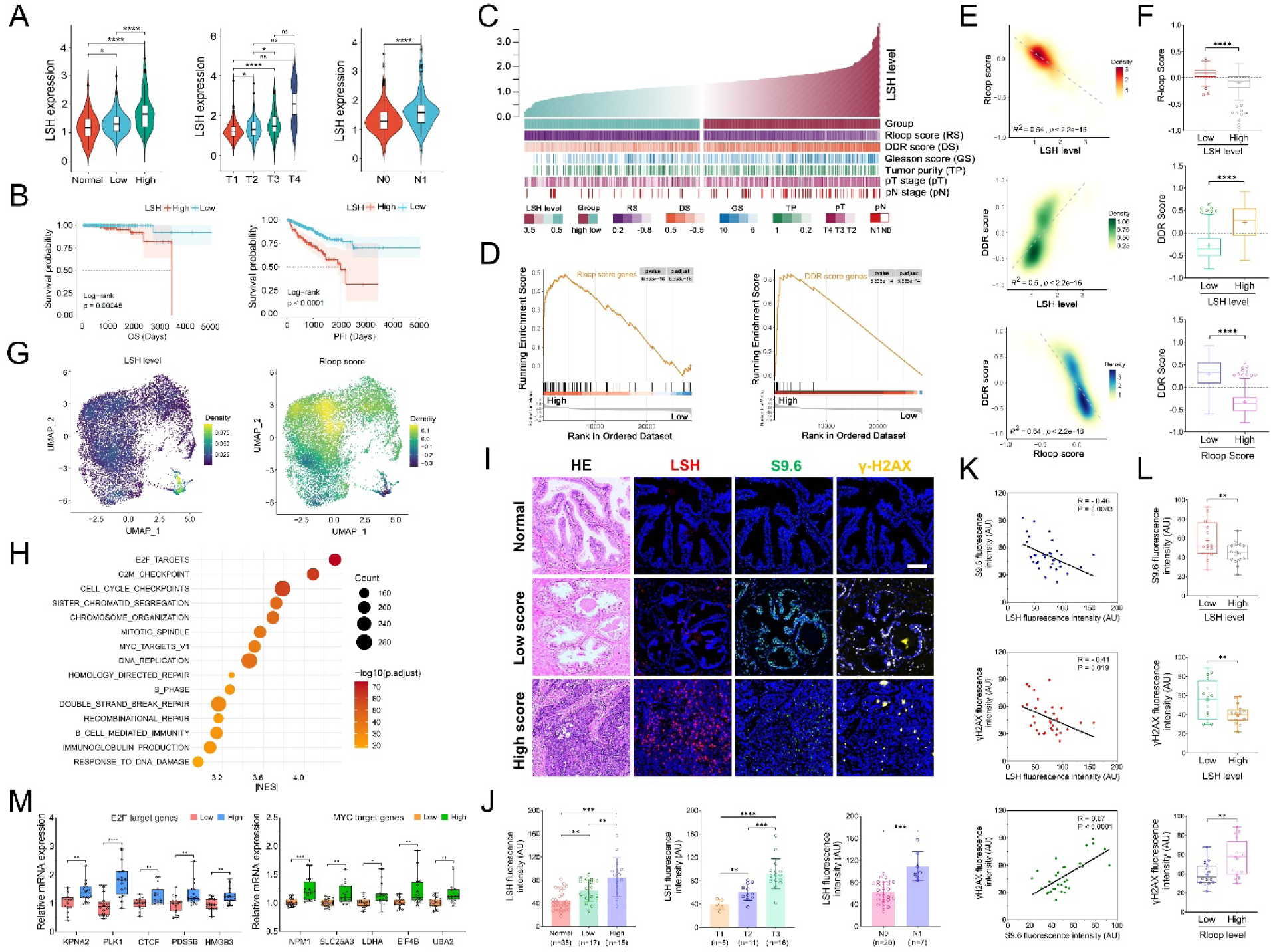
Clinical relevance of LSH level in relation to R-loops accumulation in prostate cancer. **A** Comparisons of LSH level are presented using violin plots across subgroups based on Gleason score (low vs. high, left), clinical stage (T, middle), and lymph node involvement (N, right), utilizing prostate cancer TCGA RNA-seq data. Endpoints depict minimum and maximum values, quartiles depicted by thin black lines, median depicted by thick black lines. *p < 0.05; ****p < 0.0001; ns indicates not significant, as determined by the One-way ANOVA with Tukey’s multiple comparison test or the two-tailed Student’s t test. **B** High LSH expression was significantly associated with a shorter survival rate, as indicated by overall survival (OS) and progression-free interval (PFI) data. The p-values are shown in the panel. **C** The associations between LSH levels and R-loop scores (RS), DDR scores (DS), and clinicopathological features of prostate cancer are presented. The columns represent samples arranged according to LSH levels, ranked from low to high (top row). The subsequent rows display the corresponding R-loop scores, DDR scores, and clinicopathological features associated with varying LSH levels. **D** Enrichment plots for genes related to R-loop and DDR scores were generated. Patients were classified into high and low LSH expression groups for analysis. **E** Pearson correlation analyses were conducted to assess the relationships between LSH levels, R-loop scores, and DDR scores. The R^2^ values and p-values are shown in the panel. **F** Prostate cancer patients with higher LSH levels exhibited lower R-loop scores and higher DDR scores, while patients with higher R-loop scores had lower DDR scores. The data are presented as a Tukey-style box plot, where the center line represents the median value, and the boxes and whiskers indicate the 25th to 75th percentiles and the 5th to 95th percentiles, respectively. Outliers are represented as individual points. ****p < 0.0001, as determined by the two- tailed Student’s t test. **G** Characterization of LSH expression and R-loop score patterns in prostate cancer cells was conducted using the single-cell RNA-seq dataset GSE141445[88]. **H** A bubble chart illustrated the pathway analysis of prostate cancer RNA-seq data exhibiting concurrently high LSH levels and low R-loop scores. **I** Representative images of H&E staining and immunofluorescence staining for LSH, S9.6, and γ-H2AX in prostate cancer specimens collected from real-world sources, along with adjacent normal tissue. The prostate cancer samples were categorized into high and low Gleason score groups. The scale bar denotes 100 μm. **J** Quantification of LSH immunofluorescence staining in real-world prostate cancer samples, stratified by Gleason score (low vs. high, left), clinical stage (T, middle), and lymph node (N, right). Data are presented as a bar graph with mean ± s.d. **p < 0.01; ***p < 0.001; ****p < 0.0001, as determined by the two-tailed Mann–Whitney test. AU denotes arbitrary units. **K** Pearson correlation coefficients for fluorescence intensity were calculated among the immunostaining results of LSH, S9.6, and γ-H2AX in real-world prostate cancer samples. The R values and p-values are shown in the panel. AU denotes arbitrary units. **L** Prostate cancer samples from real-world patients with elevated LSH levels exhibited reduced S9.6 and γ-H2AX staining, whereas samples with increased R-loop levels showed heightened γ-H2AX staining. The data are presented as a Tukey-style box plot, where the center line represents the median value, and the boxes and whiskers indicate the 25th to 75th percentiles and the 5th to 95th percentiles, respectively. Outliers are represented as individual points. **p < 0.01, as determined by the two- tailed Student’s t test. AU denotes arbitrary units. **M** RT-qPCR analysis was conducted on the selected E2F and MYC target genes in real-world prostate cancer patients, who were classified into low and high LSH expression groups. The mRNA levels of E2F and MYC target genes are displayed in a box plot, where the center line indicates the median, and the boxes and whiskers represent the 25th to 75th percentiles and the 5th to 95th percentiles, respectively. *p < 0.05, **p < 0.01, ***p < 0.001; ****p < 0.0001, as determined by the two-tailed Mann–Whitney test. Source data are provided as a Source Data file.

To illustrate the molecular pathway regulated by LSH-induced R-loop formation, a pathway analysis of RNA-seq data is presented using a bubble chart, derived from prostate cancer patients exhibiting concurrently high levels of LSH and low R-loop scores (**Figure 8H**). The findings align with our previous results, showing that LSH-induced R-loop formation is significantly associated with the expression of E2F and MYC target genes, as well as pathways related to DNA replication and DNA damage repair (**Figure 8H**). Additionally, it involves genes associated with cell cycle regulation, chromosome reorganization, and cellular immune response, among others, as previously reported[25, 27, 29, 32, 62] (**Figure 8H**). The GSEA results also suggest that prostate cancer patients with higher LSH level were enriched of E2F and MYC target genes (**Figure S12E**), and Pearson correlation analysis demonstrated that LSH level was positively related with the expression of R-loop suppressor FANCD2, DNA damage repair checkpoints (BRCA1, RAD51), proliferation marker Ki67, as well as E2F and MYC target genes (**Figure S12F**). We also analyzed the patterns of gene co-mutation and mutual exclusivity in patients with low and high levels of LSH. The results indicate that a greater number of mutations occurred in the high LSH group (**Figure S12G**). Each group exhibited a distinct set of top mutated genes, with significant differences in mutation rates for shared genes between the groups (**Figure S12H**). Furthermore, patients with high LSH expression showed a significant co-occurrence with mutations in the tumor suppressor protein TP53 (**Figure S12I**). These findings illustrate that the abnormally high expression of LSH is closely associated with tumor aggressiveness, possibly owing to its important role in regulating R-loop formation.

To validate our findings regarding the critical role of LSH in R-loop formation, we collected real-world prostate cancer samples from the Department of Urology at Shanghai Sixth People’s Hospital. Representative images of H&E staining and immunostaining for LSH, S9.6, and γ-H2AX using prostate cancer specimens revealed distinct expression patterns between the low and high Gleason score groups, with adjacent normal tissues serving as a control group (**Figure 8I**). Specifically, the intensity of LSH immunofluorescence is significantly higher in patients with a high Gleason score (> 7), advanced pathological T stage, and lymph node metastasis (**Figure 8J**). This finding is consistent with the LSH levels measured by RT-qPCR (**Figure S13A**). Furthermore, there is a strong correlation between the LSH levels measured by immunofluorescence and RT-qPCR, as analyzed using Pearson correlation (**Figure S13B**). We also analyzed R-loops accumulation and DNA damage levels using S9.6 and γ- H2AX staining in real-world samples (**Figure 8I**). The intensity of S9.6 and γ-H2AX immunostaining was lower in patients with a high Gleason score (> 7), advanced pathological T stage, and lymph node metastasis (**Figure S13C-D**). Additionally, we conducted an analysis of the Pearson correlation coefficients among these parameters using real-world samples. The findings revealed that LSH exhibits a negative correlation with both R-loop and γ- H2AX levels, whereas R-loop levels display a positive correlation with γ- H2AX levels (**Figure 8K**). Meanwhile, we categorized the samples based on LSH and R-loop levels. We found that the low LSH group exhibited elevated R-loop and γ-H2AX levels, while the low R-loop group displayed reduced γ- H2AX levels (**Figure 8L**). Therefore, these results support our findings that LSH can downregulate R-loop formation and DNA damage in prostate cancer using real-world samples. Finally, we conducted RT-qPCR analysis of selected E2F and MYC target genes in real-world prostate cancer samples (**Figure 8M**), along with RNA-seq analysis from TCGA-PRAD (**Figure S13E**). Both analyses consistently revealed that E2F and MYC target genes were significantly elevated in samples with high LSH levels (**Figure 8M**, **Figure S13E**). This finding aligns with the pathway enrichment results (**Figure 8H**). Given that E2F and MYC targets are recognized for their critical roles in cancer cell proliferation and tumorigenesis, these results reinforce our findings that LSH promotes cell proliferation in prostate cancer cells by modulating E2F and MYC target pathways through the regulation of R-loop formation.

Overall, this comprehensive clinical investigation reveals that high levels of LSH are associated with reduced R-loops accumulation and enhanced DNA damage response, which are significantly linked to a higher degree of malignancy and poorer prognosis in prostate cancer.

## Discussion

Genome integrity is perpetually challenged by the very processes essential for its function. Critical cellular activities such as transcription, DNA replication, and repair mechanisms frequently occur concurrently within shared genomic regions, creating topological conflicts and torsional stress in the DNA molecule[63, 64]. These competing demands predispose the genome to structural instability, increasing the likelihood of DNA breaks and lesions. Transcription is widely recognized as a key contributor to genome instability. The process of DNA unwinding by RNA polymerases during transcriptional elongation generates significant torsional stress, which in turn promotes widespread DNA supercoiling throughout the genome[63–66]. Notably, the heightened transcriptional activity required to sustain rapid cancer cell proliferation induces profound mechanical strain on the DNA helix, including severe distortions and torsional stress, presenting a substantial challenge to maintaining genome stability[19, 67]. Three-stranded RNA–DNA hybrids, termed R-loops, form between the nascent RNA and template DNA upstream of RNA polymerases[9]. These structures can impede transcriptional elongation and compromise chromatin stability[8, 9]. In contexts requiring high transcriptional outputs, such as in rapidly proliferating cancer cells, RNA polymerase alone often fails to traverse gene bodies efficiently due to R- loop-induced structural barriers[8–10]. Consequently, auxiliary mechanisms are likely necessary to facilitate polymerase progression through nucleosomes and prevent aberrant transcriptional arrest. Transcriptionally active genes are predominantly localized within nucleosome-deprived regions, which enable the binding of RNA polymerase and transcription factors (TFs)[65, 68]. According to the prevailing model, pioneering TFs recruit ATP-dependent chromatin remodelers to displace or reposition nucleosomes, thereby granting RNA polymerase access to DNA[69, 70]. This rapid and tightly regulated cycle of nucleosome eviction and reloading is critical for the precise regulation of transcriptional programs, ensuring proper cellular identity and function[70, 71]. However, disruptions in this dynamic equilibrium, commonly observed in diseases such as cancer, lead to aberrant chromatin states that promote tumorigenesis[72, 73]. In this context, faithful chromatin remodeling activity is essential for orchestrating the spatiotemporal regulation of transcription while maintaining chromatin architecture[72–74].

LSH, a member of the SNF2 family of chromatin-remodeling proteins, plays critical roles in epigenetic regulation and genome stability[23, 24, 26]. Like many ATP-dependent chromatin remodelers, LSH is frequently dysregulated in cancer, with aberrant expression serving as a hallmark of both solid tumors and hematological malignancies[23, 34, 35, 75]. Over-expression of LSH promotes tumorigenesis and malignant progression, underscoring its oncogenic potential[23, 35, 75, 76]. Mechanistically, LSH contributes to chromatin instability; however, the precise molecular pathways underlying its role in genomic instability are still not well understood in cancer cells. Emerging evidence indicates that LSH is essential for DNA damage response and safeguards nascent DNA at stalled replication forks, thereby modulating tumor cell proliferation and apoptosis[23, 26, 33, 44]. Our results align with and extend previous findings by demonstrating that LSH deficiency in prostate cancer cells not only significantly elevates DNA damage levels but also compromises its protective function at replication forks, where it normally safeguards nascent DNA (**Figure 1**). Consistent with its role in genome maintenance, we found that LSH knockout impairs cellular proliferation while promoting apoptosis. Most importantly, our study provides novel mechanistic insights by establishing LSH as a critical resolver of R-loop structures in prostate cancer cells (**Figure 2**). Through genome-wide S9.6 CUT&Tag sequencing analysis, we further identify LSH as a global regulator of R-loop homeostasis (**Figure 3 and Figure S5**), a function that underlies its ability to maintain genomic integrity, mitigate DNA damage accumulation, and consequently modulate key cellular processes including proliferation and apoptosis (**Figure S1-4**).

Considering the pivotal role of LSH in tumorigenesis, elucidating the precise molecular mechanisms by which LSH resolves R-loops is crucial for understanding prostate cancer pathogenesis and developing novel therapeutic strategies. While LSH has been implicated in R-loop resolution, the precise molecular mechanisms underlying this process remain incompletely characterized and are likely multifaceted in nature. Earlier work found that LSH functions as a critical safeguard against R-loop-induced genomic instability, specifically by mitigating RNA-DNA hybrid accumulation and associated DNA damage at pericentromeric regions in ICF syndrome cells[77]. Alternatively, DDM1, the LSH homolog in Arabidopsis, can directly eliminate co-transcriptional R-loops, as demonstrated by in-vitro experiments[36]. In our study, through comprehensive in-vivo and in-vitro experiments, we have conclusively demonstrated the functional interaction between LSH and R-loops (**Figure 4**). Our findings reveal that the ATP- binding domain within LSH plays a crucial role in this process - mutations in this domain lead to abnormal R-loop accumulation, resulting in DNA damage and genomic instability (**Figure 5 and Figure S8**). Notably, unlike its plant ortholog DDM1, our in-vitro assays did not detect direct R-loops helicase activity in LSH (**Figure S7**). Through systematic investigation of alternative R-loop regulatory mechanisms, we discovered that LSH predominantly localizes replication forks and transcription-replication conflict sites (**Figure 6**). Furthermore, we identified a novel functional partnership between LSH and FANCD2 protein in modulating R-loops accumulation, thereby preventing R-loop-associated DNA damage and double-strand breaks (**Figure 6 and Figure S9**). The genomic instability induced by aberrant R- loops manifests through dual threats: replication fork destabilization and exacerbated nucleolytic DNA fragmentation, both functionally intertwined with chromatin architectural rearrangements. Previous studies have demonstrated that LSH plays a crucial role in safeguarding nascent DNA strands from degradation by facilitating RAD51 filament formation[26]. Importantly, impaired RAD51 loading has been shown to disrupt the delicate balance between BRCA1 and 53BP1 accumulation at stalled replication forks, thereby compromising the proper selection of DNA repair pathways that are essential for maintaining genomic stability during replication stress[26, 78, 79]. Our data demonstrated that the absence of LSH leads to defective RAD51 filament formation at R-loop-induced lesions, disrupting the canonical DNA repair pathway choice through reciprocal regulation of BRCA1 displacement and 53BP1 enrichment at damage sites (**Figure 4 and Figure S6**). Collectively, our results reveal LSH as a critical guardian of genome stability. It interacts with FANCD2 to resolve R-loops accumulation and nucleates RAD51 filament assembly, while also balancing BRCA1 and 53BP1 loading, thereby establishing a comprehensive DNA repair pathway for R-loop-induced genomic damage.

While previous studies have established LSH’s involvement in genome maintenance, the mechanistic link between its chromatin remodeling activity and gene expression regulation in cancer remains poorly understood. Emerging evidence now demonstrates that LSH directly and multifunctionally participates in transcriptional regulation. LSH was reported to mediate chromatin reorganization to modulate RNAPII and transcription factor (TF) binding, thereby fine-tuning cell identity-related transcriptional programs[34, 35, 80, 81]. During periods of high transcriptional demand, particularly in rapidly proliferating cancer cells, elongating RNAPII encounters significant structural barriers imposed by R-loop accumulation[9, 50, 82]. These three-stranded nucleic acid structures physically obstruct RNAPII progression through gene bodies. Consequently, successful transcriptional elongation necessitates auxiliary factors that facilitate RNAPII navigation through nucleosome landscapes while preventing aberrant transcriptional arrest[9]. Notably, our findings reveal a novel role for LSH in resolving transcription-replication conflict by eliminating promoter-proximal R-loops at oncogenic and cellular fitness genes, thereby enabling uninterrupted transcriptional elongation (**Figure 7 and Figure S11**). Specifically, LSH exerts a significant influence on transcriptional regulation in the genome by modulating R-loop formation at the promoter regions. It is critically involved in tumor-related signaling pathways, transcriptional activity, as well as cell cycle and DNA damage repair pathways (**Figure 7)**. This dual functionality not only ensures their stable expression but also safeguards genomic integrity by preventing transcription-associated structural instability. Among these, LSH interacts with oncogenic TFs, such as MYC and E2F (**Figure 7)**. TFs play a pivotal role in this regulatory network, not only through their ability to recognize specific DNA sequences and modulate local chromatin accessibility, but also by orchestrating the recruitment of chromatin remodelers that ultimately govern the three-dimensional topological architecture of the genome[71]. Our findings reveal that LSH can regulate the expression of downstream target genes of MYC and E2F by controlling R-loop accumulation at the promoter regions (**Figure 7 and Figure S10)**. The underlying molecular mechanism may involve the increased local enrichment of R-loops at promoters, which impedes the elongation mechanism of RNAPII during transcription, thereby leading to DNA damage and reduced FANCD2 accumulation (**Figure 7 and Figure S11)**. This also suggests that targeting LSH inhibition through the MYC and E2F pathways could provide a novel therapeutic strategy for prostate cancer. Finally, through analyses of public databases and collected clinical samples, we further confirmed that LSH regulates R-loop formation and significantly affects the expression of downstream target genes of MYC and E2F (**Figure 8 and Figure S12-13**). Elevated LSH expression markedly influences the progression of prostate cancer and patient prognosis (**Figure 8 and Figure S12-13**).

Overall, our study aims to elucidate the precise molecular mechanisms through which LSH modulates R-loop dynamics, thereby affecting DNA repair pathways and genomic instability in prostate cancer cells. Additionally, we further examine its functional implications in tumor proliferation and evaluate its potential for clinical translation. Our results furnished compelling evidence indicating that LSH assumes a pivotal and critical role in expediting R-loops resolution in an ATP-dependent manner, which might contribute to resolve R-loop-mediated transcription–replication conflicts in prostate cancer cells. Loss of LSH led to increased accumulation of R-loops by decreasing the recruitment of FANCD2, which in turn resulted in the accumulation of DNA damage and an impairment of Rad51 filament formation. Therefore, LSH makes a significant contribution to the preservation of DNA integrity and genome stability in prostate cancer cells. Furthermore, LSH deficiency led to the accumulation of R-loops at the promoter regions of MYC and E2F target genes, thereby inhibiting RNAPII elongation during the transcriptional process. Consequently, this disruption may have profound and far-reaching implications for the proliferation of prostate cancer cells and the effectiveness of clinical treatments. In summary, our work highlights the importance of LSH in resolving R-loop structures within chromatin, demonstrating its critical role in preserving genome integrity and supporting cell proliferation in prostate cancer cells, which is crucial for understanding prostate cancer pathogenesis and developing novel therapeutic strategies.

## Methods

### Cell culture

The cell lines utilized in this study included the androgen-independent prostate cancer cell line PC3 (ATCC), the androgen-dependent prostate cancer cell line LNCaP (ATCC), and the human embryonic kidney epithelial cell line HEK293T (ATCC). PC3 cell line was grown in F-12K medium (HyClone, USA) supplemented with 10% Fetal Bovine Serum (ExCell Bio, China) and 50 U/ml Pen/Strep. LNCaP cell line was cultured in RPMI 1640 medium (HyClone, USA) supplemented with 10% Fetal Bovine Serum (Biological Industrie, Israel) and 50 U/ml Pen/Strep. 293T cell line was grown in high-glucose DMEM medium (HyClone, USA) supplemented with 10% FBS (RMBIO, USA) and 50 U/ml Pen/Strep. All cells were cultured at 37 °C in a humidified incubator containing 5% CO2.

### Cell treatment

The LSH-shRNA sequence (5’-GATCAAGAGAGAAGGTCATTA-3’) was cloned into a Tet-on puromycin-resistant plasmid (Addgene, #21915). Lentivirus was produced by transfecting HEK293T cells with the Tet-on LSH- shRNA vector using Lipofectamine 2000 (Invitrogen), alongside co- transfection with the packaging vectors psPAX2 (Addgene, #12260) and pMD2.G (Addgene, #12259). Concentrated lentiviral supernatants by Lenti-XTM Concentrator (Takara, Japan) were used to infect PC3 cells with polybrene (8 μg/ml). Cells were incubated overnight before the virus was removed, followed by puromycin selection (2 μg/mL) to establish stably transfected PC3 cells with the lentiviral construct. LSH knockdown was induced in vitro by treating cells with 100 ng/mL doxycycline for five days, followed by a five-day washout period to restore LSH expression. To achieve LSH knockdown in LNCaP cells, following lentivirus production and infection, cells were selected with puromycin (2 μg/mL) to establish a stable cell line. Knockdown of LSH was analyzed and monitored by RT-qPCR and western blotting. For siRNA-mediated knockdown of the indicated genes, siRNAs were synthesized following standard protocols. Cells were transfected with siRNAs using Lipofectamine RNAiMAX (Invitrogen, USA) according to the manufacturer’s instructions, and analysis was conducted 48–72 hours post-transfection. Knockdown efficiency was assessed by Western blotting. The siRNA sequences are listed in **Supplementary Table 1**.

LSH knockout PC3 cells were generated using the CRISPR/Cas9 technology. Briefly, a Cas9 sgRNA targeting the exon 11 of the LSH gene, using the guide RNA sequence 5’-TATGAAGTGCCGTCTAATCA-3’, was cloned into the lentiCRISPRv2 vector (Addgene #52961) and subsequently transformed into Stbl3 competent cells. The guide RNA was designed and validated by Feng Zhang’s laboratory at the Broad Institute to specifically target the human LSH gene[83]. The lentiCRISPRv2-sgLSH was packaged into lentiviral particles by co-transfecting the packaging plasmids psPAX2 (Addgene, #12260) and pVSVG (Addgene, #8454) into HEK293T cells. Viral particles were collected, infected into PC3 cells and selected using puromycin (2 μg/mL) to generate stable cell line. Knockout efficiency was validated in the bulk cell population by western blotting.

To generate stable cell lines overexpressing either wild-type or mutant RNASEH1 or LSH, cells were transfected with RNASEH1_WT-V5 (Addgene #111906), RNASEH1_WKKD-V5 (Addgene #111905), RNASEH1_D210N-V5 (Addgene #111904)[38], LSH_WT, or LSH_K254A vectors[24]. Following transfection, cells were either sorted by flow cytometry or clonally selected in media containing hygromycin (100 mg/mL) or puromycin (2 μg/mL). Protein overexpression levels were validated by Western blotting.

### Collection of real-world prostate tumor samples

Following the receipt of patient consent and approval from the institutional research ethics committee, we collected 32 surgical resection samples of prostate tumors along with 35 adjacent normal tissues, which served as a control group. These samples were obtained from patients admitted to the Department of Urology at Shanghai Sixth People’s Hospital.

Detailed patient information is provided in **Supplementary Table 2**. For all resected tumor samples, we engaged pathology experts to ensure accurate pathological diagnoses for the patients. All samples were collected within 5 minutes post-excision and were immediately placed in liquid nitrogen for preservation.

### Cell viability assay

For the cell viability assay, PC3 and LNCaP cells were seeded at a density of 5,000 cells per well in 96-well plates. The cells were incubated at 37°C with 5% CO₂ for 24, 48, and 72 hours. To assess cell viability, the 10% Cell Counting Kit-8 (CCK8) reagent was added to each well, and the plates were incubated at 37°C in the dark for 1 hour. The assay was conducted following the manufacturer’s instructions (Glpbio Technology, USA). Absorbance was measured at 450 nm using a CLARIOstar Microplate Reader (BMG Labtech). Absorbance values at different time points were plotted to compare cell proliferation rates.

### Colony formation assay

For the colony formation assay, PC3 and LNCaP cells were seeded in 6-well plates at a density of 2,000 cells per well to minimize contact between colonies. Following treatment as indicated, the cells were incubated at 37 °C with 5% CO₂ for 10 to 14 days. Colonies were then fixed with 4% paraformaldehyde for 20 minutes and stained with 0.5% (w/v) crystal violet in 25% methanol for 15 minutes. Colony images were analyzed and quantified using ImageJ v.2 software.

### In vivo tumor formation assay

All animal experiments were approved by the Animal Care and Research Committee of Shanghai Sixth People’s Hospital (Approval No. 2021-0908) and conducted in accordance with institutional ethical guidelines. Briefly, 4-week-old male NOD-Prkdcscid Il2rgem1/Smoc (NSG) mice, obtained from the Shanghai Model Organisms Center, were subcutaneously inoculated in the right flank with 2 × 10⁶ LNCaP cells. After allowing two weeks for tumor establishment post-inoculation, tumor growth was assessed every 2-3 days through caliper measurements, and tumor volumes were determined using the standard formula (length × width²)/2. Mice were humanely euthanized when tumors reached a diameter of 1.5 cm or upon reaching predefined experimental endpoints. Subsequently, excised tumors were weighed and prepared for further analysis.

### Immunoblotting

A whole-cell extract was prepared using 2 × 10^6^ cells. Cells were centrifuged at 200 × g, 4° for 5 minutes and washed twice with PBS. Next, 100 μL of RIPA lysis buffer was added to resuspend the cell pellet and placed on ice for 60 minutes. Afterwards, cells were centrifuged for 15 minutes at 14,000×g, 4 °C. The supernatant was collected to determine the protein concentration by bicinchoninic acid assay (BCA) assay (Thermo Fisher, USA). Samples were boiled in Laemmli sample buffer (Bio-Rad) at 95 °C for 5min. Equal amounts of protein were loaded onto acrylamide/bis gels and transferred to PVDF membranes after electrophoresis. Following blocking in 5% nonfat milk for 1 h, membranes were incubated at 4 °C overnight in primary antibodies listed in **Supplementary Table 3**. After one hour incubation with HRP-conjugated secondary antibodies at a 1:2500 dilution (as detailed in **Supplementary Table 3**), signal detection was performed using the Amersham ECL Western blotting analysis system and exposed using a chemiluminescence gel imaging system (BIO-RAD, USA).

### Co-immunoprecipitation (Co-IP) analysis

RNA: DNA hybrids Co-IP was carried out as previously described using the S9.6 antibody[84]. Briefly, non-crosslinked nuclei were incubated in RSB buffer (10 mM Tris-HCl, pH 7.5; 200 mM NaCl; 2.5 mM MgCl2) containing 0.2% sodium deoxycholate, 0.1% SDS, 0.05% sodium lauroyl sarcosinate, and 0.5% Triton X-100, and then sonicated for 10 minutes. Samples were subsequently diluted 4-fold in RSB with 0.5% Triton X-100 (RSB+T) prior to immunoprecipitation (IP) with the S9.6 antibody, which was bound to protein A Dynabeads and preblocked with 0.5% BSA/PBS for 2 hours. IgG was used as a negative control. IP was performed in the presence of 0.1 ng of RNase A per microgram of genomic DNA. Beads were washed 4 times with RSB + T and 2 times with RSB. IPs were eluted either in 1x LDS, 100 mM DTT for 10 minutes at 70°C for SDS-PAGE, or in 1% SDS and 0.1 M NaHCO3 for 30 minutes at room temperature for RNA: DNA hybrid slot blot. RNA: DNA hybrid competitors were added during the IP step at a concentration of 1.3 mM. RNA: DNA hybrids were prepared as described with ssDNA (CGGTGTGAATCAGAC) and ssRNA (GUCUGAUUCACACCG). IP samples were then loaded onto acrylamide/bis gels and subjected to western blotting with anti-LSH and anti-lamin B1 antibodies (as listed in **Supplementary Table 3**).

### Immunofluorescence (IF) staining

Cells cultured on chamber slides were fixed with 4% paraformaldehyde for 20 minutes, followed by permeabilization with 0.1% Triton X-100 for 15 minutes. The cells were then blocked with 5% BSA for 30 minutes and incubated overnight at 4°C with the primary antibodies listed in **Supplementary Table 3**. The following day, the slides were incubated with Alexa fluorophore–conjugated secondary antibodies for 1 hour, as detailed in **Supplementary Table 3**. Afterward, the slides were stained with DAPI and imaged using confocal microscopy. Image analysis was performed using ImageJ v. 2 software.

### EdU incorporation and detection

The Click-iT EdU Cell Proliferation Kit for Imaging (Thermo Fisher Scientific, USA) was used to assess DNA replication via EdU incorporation, following the manufacturer’s instructions. Briefly, cells were cultured in complete medium supplemented with 10 μM EdU for 30 minutes. The samples were then fixed, permeabilized, and subjected to the Click-iT reaction according to the manufacturer’s guidelines. Finally, nuclei were stained with DAPI and imaged using confocal microscopy. Image analysis was performed with ImageJ v. 2 software. The nuclear intensity of the entire EdU-positive population and the percentage of cells incorporating EdU were determined, with EdU intensity measured only in cells that had incorporated EdU.

### H&E staining

The collected prostate cancer tissue specimens were fixed in formalin and embedded in paraffin. Prostate cancer sections were cut from paraffin embedded tissue blocks, soaked with 10% formaldehyde, and fixed on slides. The slides were incubated at 42°C overnight. For pathological evaluation, all sections were stained using H&E. Briefly, the slides were placed in xylene twice for 5 minutes each, followed by sequential immersion in 100%, 90%, 80%, and 70% ethanol for 5 minutes each, and then rinsed in distilled water for 5 minutes. Next, the slides were stained with hematoxylin for 5 minutes, rinsed in distilled water for 3 minutes, and differentiated in 1% hydrochloric acid in ethanol for 3 seconds. After washing in distilled water, the slides were treated with 0.8% ammonia for 1 second, rinsed again in distilled water for 5 minutes, and stained with 0.5% eosin for 1 minute. The sections were then dehydrated through graded ethanol (80%, 90%, 95%, and 100%), cleared in xylene twice for 1 minute each, sealed, and imaged under a microscope.

### RT-qPCR analysis

Total RNA was extracted using the RNeasy Plus Mini Kit (Qiagen, Germany), followed by an additional RNase-free DNase treatment to remove any residual DNA. Reverse transcription was carried out using the PrimeScript RT Reagent Kit (Takara). Quantitative PCR was performed on the MyiQ2 Real-Time PCR detection system (Bio-Rad, USA) with iTaq Universal qPCR Mastermix (Bio-Rad). Gene expression levels were normalized to GAPDH as internal control. Three biological replicates were analyzed, and quantitative amplification was performed immediately using the fluorescent qPCR system (Bio-Rad, USA). The primer sequences are provided in **Supplementary Table 4**.

### ChIP-qPCR assay

For the ChIP assay, 1 × 10⁷ cells were cross-linked with 1.42 % formaldehyde in PBS for 15 minutes at room temperature, followed by quenching with 125 mM glycine for 5 minutes. Cells were then scraped, collected by centrifugation at 2000 × g for 5 minutes at 4°C, and lysed in IP buffer (Pierce) containing protease inhibitors, following the manufacturer’s instructions. The nuclear pellet was washed, resuspended in IP buffer, and subjected to sonication on ice to shear chromatin into 200-800 bp DNA fragments. The sheared chromatin was incubated overnight at 4°C with the indicated ChIP-grade antibodies (listed in **Supplementary Table 3**). After reversing the cross-links, the precipitated DNA was suspended in 50 μl of nuclease-free water and purified using the QIAquick PCR Purification Kit (Qiagen, Germany). The purified ChIP DNA was analyzed by qPCR using a MyiQ2 Real-Time PCR Detection System (Bio-Rad, USA). ChIP data were normalized as a percentage of input, and the specific primers used for qPCR are listed in **Supplementary Table 4**.

### DNA fiber analysis

For the DNA fiber assay, cells were seeded in six wells plate at about 20% confluency on the previous day. Cells were first incubated with 50 μM CIdU for 20 min at 37 °C followed by three times wash with 37 °C PBS, and then incubated with 250 μM IdU for 20 min at 37 °C. Cells were harvested and resuspended in cold PBS to 2 × 10^5^ cells/ml and 2.5 μl of resuspended cells were mixed directly with 7.5 μl of lysis buffer (200mM Tris–HCl pH 7.4, 50mM EDTA, 0.5% SDS) on slides. The slides were left horizontally for 8 min at room temperature and then tilted to allow drops to run slowly down the slides, followed by air dry and fixation in 3:1 methanol: acetic acid solution overnight at 4 °C. Slides were rehydrated in PBS and DNA denatured in 2.5M HCl. Slides were washed several times in PBS until pH was back to 7.5. Afterwards slides were blocked in 2% BSA, 0.1% Tween 20 in PBS, and immuno-fluorescence staining performed. Slides were mounted with prolong gold antifade mounting medium (Thermo Fisher Scientifi, USA). Antibodies employed for immunofluorescence were as follows: anti-CIdU (Abcam, ab6326, 1:300), anti-IdU (BD bioscience, #347580, 1:500), goat anti-mouse Alexa Fluor 488-conjugated (Cell Signaling Technology, 4408S, 1:1000), goat anti-rat Alexa Fluor 555-conjugated (Cell Signaling Technology, 4417S, 1:1000). Images were captured using confocal microscopy with a 63× objective and analyzed using ImageJ v.2 software.

### Proximity ligation assay (PLA)

PLA was conducted using the Duolink PLA Technology (Sigma, USA) following the previously established protocols. Samples were first pre-extracted, fixed, permeabilized, and incubated with two primary antibodies (detailed in **Supplementary Table 3**) following the procedure used for immunofluorescence assays. Subsequently, secondary antibody binding, ligation, and amplification were performed according to the manufacturer’s instructions. The PLA reaction was carried out using Duolink in situ PLA probes: anti-rabbit PLUS and anti-mouse MINUS, along with Duolink Detection Reagents Red (Sigma, USA). Nuclei were stained with DAPI, and the samples were mounted in ProLong Gold AntiFade reagent (Invitrogen, USA). Images were then captured using confocal microscopy with a 63× objective lens, and analysis was conducted using ImageJ v.2 software to quantify the number of PLA foci per cell across all conditions.

### Neutral comet assay

For the neutral comet assay, slides were precoated with 1% low-gelling-temperature agarose. Harvested cells were prepared as a single-cell suspension at a concentration of approximately 2 × 10⁴ cells/ml in PBS, then mixed with 1% low-gelling-temperature agarose at 40°C. The cell-agarose mixture was carefully pipetted onto the agarose-precoated slides, ensuring no air bubbles formed. After the agarose solidified, the slides were gently submerged in a neutral lysis solution (N1) containing 2% sarkosyl, 0.5 M EDTA-Na₂, and 0.5 mg/ml proteinase (pH = 8.0) at 4°C. The samples were incubated at 37°C for 18–20 hours in the dark. Following incubation, the slides were removed and washed three times in a rinse buffer (N2) composed of 90 mM Tris, 90 mM boric acid, and 2 mM EDTA-Na₂ (pH = 8.5). The slides were then placed in an electrophoresis chamber filled with fresh N2 solution, and electrophoresis was conducted for 25 minutes at 0.6 V/cm. After electrophoresis, the slides were neutralized in distilled water and stained with 2.5 μg/ml propidium iodide for 20 minutes. Excess stains were removed through several washes, and images were captured using confocal microscopy with a 20× objective lens. At least 50 comet images per slide were analyzed, and tail lengths were measured using ImageJ v.2 software.

### Protein expression and purification

The synthetic full-length cDNA of human LSH (WT) or the LSH (K254A) mutant was subcloned into the pDest-636 vector, which contains N-terminal His6 and MBP tags, using Gateway LR recombination (Thermo Fisher, USA). The reaction mixture was transformed into DH10B cells, and colonies were selected on LB plates containing Ampicillin. To construct the recombinant baculovirus, bacmid DNA was generated, and Tni-FNL cells were transfected following previously described protocols. At 72 hours post-infection, Tni-FNL cells were harvested and lysed in lysis buffer (20 mM HEPES, 300 mM NaCl, 1 mM TCEP, and 1:100 v/v protease inhibitor). The lysate was passed twice through an M-110EH-30 microfluidizer (Microfluidics) at 7000 psi to ensure complete lysis. The lysate was clarified by centrifugation at 100,000 × g for 30 minutes at 4°C. The clarified supernatant was filtered through a 0.45 μm filter and loaded onto a 20 mL IMAC HP column (GE Scientific) pre- equilibrated with LB supplemented with 50 mM imidazole (10% Buffer B) using a Bio-Rad NGC system. Proteins were eluted using a 10-column volume gradient of 10–100% Buffer B (LB + 500 mM imidazole). To remove the His6-MBP solubility tag, 5% (v/v) TEV protease was added to the pooled elution, and the mixture was dialyzed using a 10K MWCO dialysis membrane (Life Technologies). The dialyzed samples were loaded onto IMAC HP columns pre-equilibrated with LB. Bound proteins were eluted using a 10- column volume gradient of 0–10% Buffer B, followed by a 10-column volume gradient of 10–100% Buffer B. The final elution pool was dialyzed into LB and concentrated using a 10K MWCO spin concentrator (Amicon) at 4000 rpm and 4°C. Protein concentration was measured, and the samples were stored at −80°C.

### Electrophoretic mobility shift assay (EMSA)

The oligonucleotides for the EMSA assay were designed based on a previous study[15]. The sequences of these oligonucleotides are listed in **Supplementary Table 5**. The different DNA: RNA hybrids or R-loops were prepared using an annealing buffer containing 10 mM Tris-Cl (pH 7.5), 7 mM MgCl₂, 100 mM NaCl, and 1 mM freshly added DTT. The mixture was heated to 95 °C for 5 minutes, then cooled to 37 °C for 1 hour, and finally allowed to cool slowly to room temperature overnight. The EMSA binding buffer and purified LSH WT or LSH (K254A) protein were pre-treated with RNase Inhibitor at 37°C for 30 minutes. The binding assay was performed under previously established conditions. For the binding assays, the indicated concentrations of LSH were incubated with 0–20 nM substrate in a binding buffer containing 17 mM Tris-Cl (pH 7.8), 12 mM Hepes (pH 7.7), 6 mM MgCl₂, 15 mM NaCl, 55 mM KCl, 0.3 mM EDTA, 5.5% glycerol, 1.25 mM DTT, 0.05% Nonidet P-40, 0.05% Triton X-100, and 0.25 mM phenylmethylsulfonyl fluoride. The binding reactions were incubated at 37 °C for 1 h. Reactions were then stopped by adding a 10-fold molar excess of sheared salmon sperm DNA as a competitor and allowed to terminate for 1 hour. The samples were separated on a 15 % PAGE gel, and the FAM signal was detected using the Cy3 channel on a Bio-Rad Chemidoc imager.

### R-loops or RNA: DNA hybrids unwinding assay

For the unwinding assays, the LSH_WT recombinant protein was mixed with 100 nM R-loops or RNA: DNA hybrids, with the RNA 5’-terminus labeled with 6-FAM fluorescence, in a 10 μL reaction volume of unwinding buffer containing 50 mM NaCl, 10 mM Tris-HCl (pH 7.5), 10 mM MgCl2, 0.1M KCl, 1 mM DTT, and 1 mM ATP, 0.1 μg/μL BSA, and 5% glycerol. The unwinding buffer was pre-incubated with RiboLock RNase Inhibitor at 37°C for 1 hour to prevent RNA degradation. The reaction mixture was incubated at 37°C for 1 hour. For the 95°C unwinding reaction, 100 nM R-loop or RNA: DNA hybrid was directly placed in the unwinding buffer and incubated at 95°C for 5 minutes. The reactions were stopped by adding 1 μL 1% SDS and 1 μL 10 μg/μL Proteinase K, followed by incubation at 37°C for 20 minutes. The products were separated on a 15% PAGE gel, and the FAM signal was detected using the Cy3 channel on a Bio-Rad Chemidoc imager.

### ATPase activity assay

The ATPase activity assay was conducted as previously described with modifications[24]. In brief, nucleosomes were reconstituted through salt dialysis onto a biotinylated 208 bp DNA fragment containing sea urchin 5S rDNA. For the ATPase activity assay, 50 nM of purified LSH WT or LSH (K254A) was mixed with 15 nM dsDNA, nucleosomes, or various R-loops and RNA:DNA hybrids in a 100 μl ATPase buffer (25 mM Tris–HCl, pH 7.5; 50 mM NaCl; 1.5 mM MgCl2; 1 mM ATP; 2% glycerol; 100 μg/ml BSA; 0.5 mM β-mercaptoethanol; and 0.05 mM EDTA) and then incubated at 25°C for 30 minutes. Reactions conducted without cofactors served as controls. ATPase activities were measured using a Colorimetric ATPase Assay Kit (NOVUS) that included PiColorLock Gold reagent and purified Pi-free ATP to eliminate background signals. The ATPase reaction rate was calculated by dividing the concentration of phosphate generated in the samples by the total reaction time. The results were analyzed using GraphPad Prism version 8.0 software.

### R-loop CUT&Tag qPCR and sequencing

R-loop profiles were obtained using the S9.6 CUT&Tag method as previously described[15], utilizing the Active Motif CUT&Tag kit (53165) according to the manufacturer’s protocol. In brief, 1 × 10^6^ cells were harvested and washed twice with ice-cold wash buffer supplemented with protease inhibitors. Concanavalin A beads were then added to the cell suspension and incubated with rotation at room temperature for 10 minutes. Bead-bound cells were separated from the supernatant using a magnetic rack and resuspended in ice-cold antibody buffer containing protease inhibitors. S9.6 antibody (2 μg) (Active Motif, 65683) was added to the bead-bound cells and incubated overnight at 4°C, with negative control samples treated with 10 U of ribonuclease (RNase) H. Following overnight incubation, the cells were treated with a secondary antibody diluted in dig-wash buffer with protease inhibitors and rotated for 1 hour at room temperature. The cells were washed three times with dig-wash buffer before the addition of pA-Tn5 transposomes diluted in Dig-300 buffer with protease inhibitors, which were incubated with the bead-bound cells for 1 hour at room temperature. Excess transposomes were removed by washing the cells three times with Dig-300 buffer. The samples were then tagmented in 40 μl of tagmentation buffer at 37°C for 1 hour. To terminate the tagmentation reaction, 2.3 μl of 0.5 M EDTA, 2.8 μl of 10% SDS, and 0.5 μl of proteinase K (10 μg/μl) were added. Samples were then incubated at 55°C for 1 hour to solubilize DNA fragments before proceeding with column purification. DNA was eluted, and libraries were amplified using Illumina i7/i5 indexed primers. Library cleanup involved an initial 1.1x SPRI bead purification, followed by size selection using 0.5x and 1x Ampure XP beads (Beckman Coulter, A63880). Finally, the CUT&Tag products were next either analyzed using qPCR (detailed in **SupplementaryTable 4**) or sequenced on an Illumina NovaSeq 6000 platform (Frasergen, China).

### R-loop CUT&Tag data processing and analysis

Sequencing data quality was assessed using FastQC (v0.11.8), and low-quality reads were discarded. Residual adapter sequences were removed with Trimmomatic (v0.39). For CUT&TAG-seq analysis, filtered reads were aligned to the human reference genome (GRCh38/hg38) using Bowtie2 (v2.2.4). Track files were generated via the BamCoverage command in deepTools (v3.5.1), while peak calling was performed with MACS2 (v2.2.7.1)[85]. CUT&Tag peak annotation was carried out using the ChIPseeker R package[86], linking peaks to the nearest genes based on GRCh38/hg38 annotations within a transcription start site (TSS) window of ±3 kb[87]. Differential binding analysis was conducted using the DiffBind R package (v2.13.1). Significant peaks (q < 0.05) were merged, and only high-confidence peaks—enriched in at least two of three replicates—were retained for further analysis. Heatmaps and metaplots of specific peaks and protein-coding genes were generated using deepTools. To quantify the R-loop signal in expressed genes, we applied the peak set derived from RNA sequencing data in PC3 cells. Fold change was calculated based on the average normalized coverage (rpkm) across the regions using R/Bioconductor packages, and enrichment around the TSS was visualized with deepTools. R-loop gain peaks were identified using a fold change > 2 and p-value < 0.05. Finally, the enriched peaks were visualized using IGV software.

### RNA sequencing

Total RNA was extracted from LSH_WT, LSH_KO and KO+RNH1 PC3 cells using the RNeasy Plus Mini Kit (Qiagen, Germany) followed by an additional RNase-free DNase treatment to remove any residual DNA, with three biological replicates. RNA integrity was evaluated using the RNA Nano 6000 Assay Kit on the Bioanalyzer 2100 system (Agilent Technologies, USA). Total RNA from each sample was used to prepare RNA libraries using the NEBNext Ultra™ RNA Library Prep Kit for Illumina, following the manufacturer’s protocols. Library quality was assessed using the Agilent Bioanalyzer 2100 system. Raw data, consisting of 150 bp paired-end reads, were generated using the Illumina NovaSeq 6000 platform.

### RNA-seq data processing and analysis

RNA quality was assessed using the Bioanalyzer 2100 system with the RNA Nano 6000 Assay Kit. mRNA was purified from total RNA using magnetic beads coated with poly-T oligos. The purified mRNA was then fragmented using divalent cations at elevated temperature. First- and second-strand cDNA synthesis was performed using random primers, M-MuLV Reverse Transcriptase, DNA Polymerase I, and RNase H. The cDNA ends were polished and adenylated, followed by ligation of an adaptor with a hairpin loop structure to the cDNA. cDNA fragments approximately 370–420 bp in length were selected using the AMPure XP system. PCR amplification was performed using Phusion High-Fidelity DNA polymerase, universal PCR primers, and Index (X) Primer. The PCR products were purified with the AMPure XP system, and library quality was assessed using the Agilent Bioanalyzer 2100 system. Index-coded samples were clustered using the cBot Cluster Generation System and the TruSeq PE Cluster Kit v3- cBot-HS (Illumina). Libraries were then sequenced on the Illumina NovaSeq platform, generating 150 bp paired-end reads. Raw data in FASTQ format were processed using the fastp software. Reads containing adapters, poly- N sequences, or of low quality were removed to obtain clean data. Q20, Q30, and GC content were calculated for quality control. The reference genome and gene model annotation files were downloaded from the genome database. A reference genome index was built using Hisat2 v2.0.5, and clean reads were aligned to the reference genome with the same version. The gene model annotation file was used by Hisat2 to create a splice junction database, improving the accuracy of the mapping results. Mapped reads were assembled in a reference-based manner using StringTie (v1.3.3b). StringTie employs a new network flow algorithm, along with an optional de novo assembly step, to generate and quantify full-length transcripts for each gene locus. The number of reads for each gene was counted using featureCounts v1.5.0-p3. The FPKM for each gene was calculated based on gene length and read counts.

### Single-cell RNA-seq data processing

The GSE141445[88] scRNA-seq dataset was processed uniformly using the Seurat package (v5.1.0)[89]. Cells with fewer than 200 genes, over 20% mitochondrial gene expression, or features present in fewer than three cells were excluded from the analysis. Normalization followed Seurat’s default settings, and 2,000 hypervariable features were selected for downstream analysis. Sample integration and batch effect correction were performed using anchor-based canonical correlation analysis (CCA), followed by data scaling with the ScaleData function. Dimensionality reduction and clustering of the integrated RNA assay were accomplished through principal component analysis (PCA) using the RunPCA function, and Louvain clustering was applied on the first 30 principal components at a resolution of 1. The results were visualized in two-dimensional space using uniform manifold approximation and projection (UMAP). Cluster annotation was performed based on marker genes using the SingleR package (v2.7.4)[90]. Epithelial cells were subsequently isolated for further study, and the density distributions of RloopScore and LSH were plotted with the plot_density function from the Nebulosa package (v1.15.0).

### Differential Expression Analysis

Differential expression (DE) analysis was conducted using the Limma R package on bulk RNA-seq data[91]. Expression data and metadata were imported into R software. The expression values were log2-transformed and transposed to align samples as columns and genes as rows. A DGEList object was then created, and normalization factors were calculated using the calcNormFactors function. Subsequently, model design and transformation were performed. For linear model fitting and hypothesis testing, a linear model was fitted using the lmFit function. Contrasts were defined to compare two groups using the makeContrasts function. Estimated coefficients and standard errors were computed with contrasts.fit, followed by Empirical Bayes moderation using the eBayes function. Finally, differentially expressed genes were identified and ranked using the topTable function.

### Gene Set Enrichment Analysis (GSEA)

Reference gene sets for GSEA analysis, including GO:BP (GO biological process gene sets), GO:CC (GO cellular component gene sets), GO:MF (GO molecular function gene sets), hallmark gene sets, CGP (chemical and genetic pertubations gene sets), KEGG_MEDICUS (KEGG Medicus gene sets), KEGG_LEGACY (KEGG Legacy gene sets), oncogenic gene sets, and Reactome gene sets, were downloaded from the Molecular Signatures Database (MSigDB, http://www.gsea-msigdb.org/gsea/index.jsp)[92]. The Investigate Gene Sets tool (http://www.gsea-msigdb.org/gsea/msigdb/annotate.jsp) was utilized to identify overlapping gene sets within the submitted gene lists in MSigDB. GSEA was performed on pre-ranked differentially expressed (DE) gene lists using the clusterProfiler package (v4.4.4)[93], referencing the downloaded gene sets. Bubble and bar plots were generated using the ggplot2 R package to visualize the analysis results.

### Calculation of R-loop Score and DDR Score

Using the 92 R-loop regulators identified by Zhang et al.[59], we calculated the R-loop Score for both single-cell and bulk samples with the cal_CRDscore function from the CRDscore package (v0.1.0). The DDR Score, based on the 20 core DNA damage repair genes identified by Ding et al.[61], was calculated using the GSVA algorithm (v1.53.23).

### Somatic Mutation and CNV Analysis

We analyzed the genomic characteristics and mutation spectrum of the high and low LSH expression groups by performing somatic mutation and copy number variation (CNV) analyses. Somatic mutation and CNV data from TCGA-PRAD were downloaded, and significantly amplified or deleted genomic regions were identified using GISTIC_2.0 (v6.15.30, https://cloud.genepattern.org/gp/pages/index.jsf). The G-score for each region was calculated based on the amplitude and frequency of variations. Mutation types, frequencies, and CNVs were further analyzed and visualized using the maftools package (v2.12.0)[94]. Somatic interaction plots were generated to illustrate interactions between the top 20 mutated genes, and oncoplots were created to display the top 20 copy number alterations (CNA) in each group.

### Survival Analysis

We conducted survival analysis using the Survival package (v3.4.0) and the Survminer package (v0.4.9) based on the Kaplan-Meier method. Kaplan-Meier survival curves were plotted, and the log-rank test was used to compare survival differences between the high and low LSH expression groups.

### Data availability

The public datasets used in this study were acquired from the Cancer Genome Atlas Program (TCGA, https://portal.gdc.cancer.gov/) and the Gene Expression Omnibus (GEO, https://www.ncbi.nlm.nih.gov/geo/).

### Statistics and reproducibility

All CUT&Tag-qPCR, ChIP-qPCR, and RT-qPCR experiments were performed as independent biological replicates, with each assay conducted at least three times. The results are presented as mean ± SD. The CUT&Tag-seq and RNA-seq experiments involved biological replicates derived from three independent samples. Furthermore, all in vitro EMSA assays and unwinding assays were executed three times using independent enzyme preparations, yielding consistent results, with representative outcomes displayed. Co-IP assays and immunoblots were repeated at least three times with different biological samples, resulting in similar findings. Additionally, all in vitro ATPase activity assays were conducted three times using independent enzyme preparations, and the results are presented as mean ± SD. Micrographs shown are representative of a minimum of six images taken from at least three biological replicates. For Figures. 1F, H-J; 2E, G-I; 5D-F, H; and Supplementary Figures. 1E-G; 2E-G; 3D, G-O; 4C, F-H; 7A, D, E; 8C, D at least three independent biological samples were analyzed, resulting in similar outcomes, with representative results provided. The Pearson correlation test was utilized to assess the correlation between normally distributed variables. The log-rank test was employed to compare survival differences between groups using the Kaplan-Meier method. Statistical analyses were performed using R software (v4.3.1) or GraphPad Prism v. 8 software, employing paired or unpaired two-tailed Student’s t-test or two-tailed Mann–Whitney test for comparisons between two samples, along with one-way ANOVA with Tukey’s analysis for multiple comparisons, unless otherwise stated. All confidence intervals are reported using binomial 95% confidence intervals. A p-value of < 0.05 was considered statistically significant. In all cases, n.s. indicates P > 0.05; * indicates P < 0.05; ** indicates P < 0.01; and *** indicates P < 0.001.

### Reporting summary

Further information on research design is available in the Nature Research Reporting Summary linked to this article.

## Acknowledgements

This work was supported by the National Natural Science Foundation of China (No. 82103260, 32270777, 82370684, 82170694); the 2022 Shanghai Leading Talent Training Program (No. 026); the 2022 Shanghai Rising-Star Program (No. 22QA1407100); the Independent Science and Technology Innovation Fund of Huazhong Agricultural University (No. 2662024SYPY001, 2662024JC005); the Excellent Youth Cultivation Program of Shanghai Sixth People’s Hospital (No. ynyq202204); the Fundamental Research Funds from the Shanghai Sixth People’s Hospital (No. X-2490, ynms202405).

## Author contributions

K.N., L.S., X.H., Z.T., and Q.F. conceived and designed the study. K.N., Y.Z., K.Z., N.X., R.L., Y.C., H.J., Y.H., J.M., and G.X. performed the experiments. K.N., J.H., Y.H., W.L., and R.Y. coordinated clinical sample collection and processing. K.N. and Y.B. conducted histopathological evaluation and slide analysis. K.N. and K.Z. carried out the bioinformatic analysis. K.N., Y.Z., K.Z., Z.T. and Q.F. interpreted the data and wrote the manuscript. All authors reviewed and approved the final manuscript.

