## Supplementary Figures and Tables for "LSH-mediated resolution of R-loops mitigates transcription-replication conflicts to preserve genomic stability in prostate cancer cells"

Ni et al.

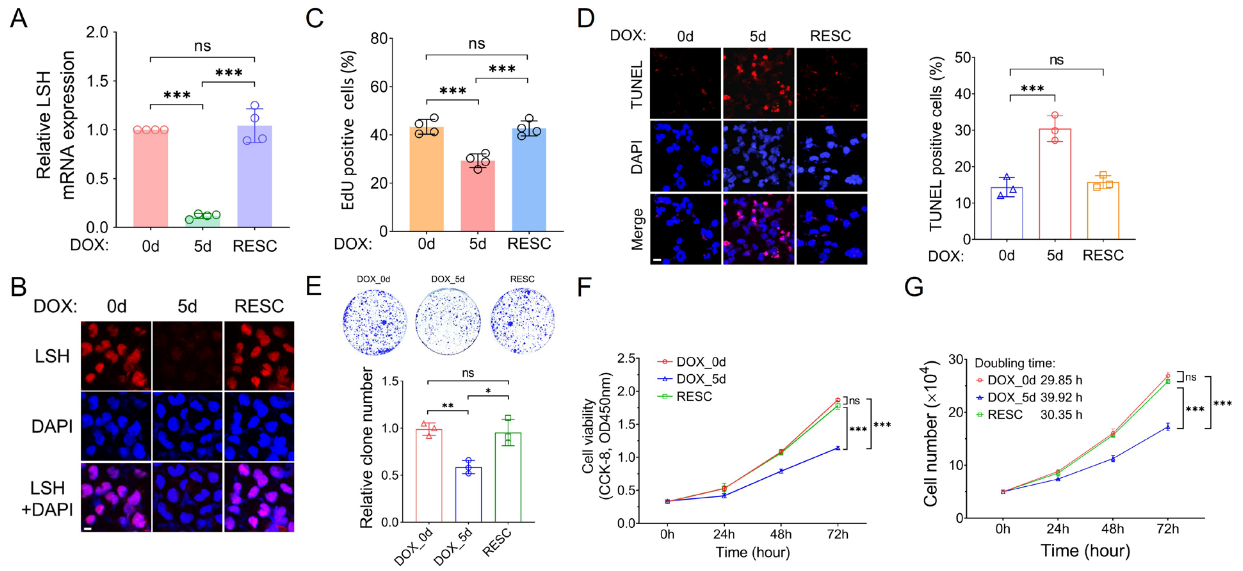


**Supplementary Figure 1: Analysis of cell proliferation and apoptosis in LSH knockdown PC3 cells using Tet-on LSH-shRNA system (corresponding to Figure 1). A** The doxycycline-inducible knockdown of LSH was analyzed using RT-qPCR to monitor changes in mRNA levels. Signal values were normalized to the Dox_0d cells and represented as a bar graph with mean ± s.d. (n = 4 biologically independent experiments). ***p < 0.001; n.s. indicates not significant, as determined by the One-way ANOVA with Tukey's multiple comparison test. **B** Representative images of immunofluorescence were captured to observe changes in LSH expression utilizing the doxycycline-inducible knockdown system in PC3 cells, with DAPI serving as a counterstain (n = 4 biologically independent experiments). The scale bar represents 10 μm. **C** The percentage of EdU-positive (EdU+) was quantified in PC3 Tet-on LSH-shRNA cells and presented as a bar graph with mean ± s.d. (n = 4 biologically independent experiments). The PC3 Tet-on LSH-shRNA cells were left untreated (Dox_0d), treated with doxycycline for five days (Dox_5d), or subjected to a five-day doxycycline washout period (RESC). ***p < 0.001; n.s. indicates not significant, as determined by the One-way ANOVA with Tukey's multiple comparison test. **D** Representative images of apoptotic cells, analyzed by the TUNEL assay (red), with nuclei counterstained by DAPI (blue), are presented for the DOX_0d, DOX_5d cells and RESC cells. The scale bar represents 20 μm. Results are presented as a bar graph with the mean ± s.d. (n = 3 biologically independent experiments). ***p < 0.001; n.s. indicates not significant, as determined by the One-way ANOVA with Tukey's multiple comparison test. **E** Representative images from the colony formation assay using Tet-on LSH-shRNA PC3 cells, treated with or without doxycycline, are presented. Cells were cultured in a 6-well dish for 14 days and subsequently stained with crystal violet to visualize colony growth. The bar graph illustrates the relative number of colonies compared to the DOX_0d group. Data are presented as a bar graph with the mean ± s.d. (n = 3 biologically independent experiments). *p < 0.05; **p < 0.01; n.s. indicates not significant, as determined by the One-way ANOVA with Tukey's multiple comparison test. **F** Cell viability in Tet-on LSH-shRNA PC3 cells, treated with or without doxycycline, was assessed using the CCK-8 assay at 24, 48, and 72 hours. Data are presented as a line graph with the mean ± s.d. (n = 3 biologically independent experiments). ***p < 0.001; n.s. indicates not significant, as determined by the One-way ANOVA with Tukey's multiple comparison test. **G** Proliferation curves of Tet-on LSH-shRNA PC3 cells, treated with or without doxycycline, were monitored over a 72-hour period. A total of fifty thousand cells were seeded and cultured, with cell counts recorded at 24, 48, and 72 hours. Data are presented as a line graph with the mean ± s.d. (n = 3 biologically independent experiments). The doubling time was calculated for all groups. ***p < 0.001; n.s. indicates not significant, as determined by the One-way ANOVA with Tukey's multiple comparison test. Source data are provided as a Source Data file.

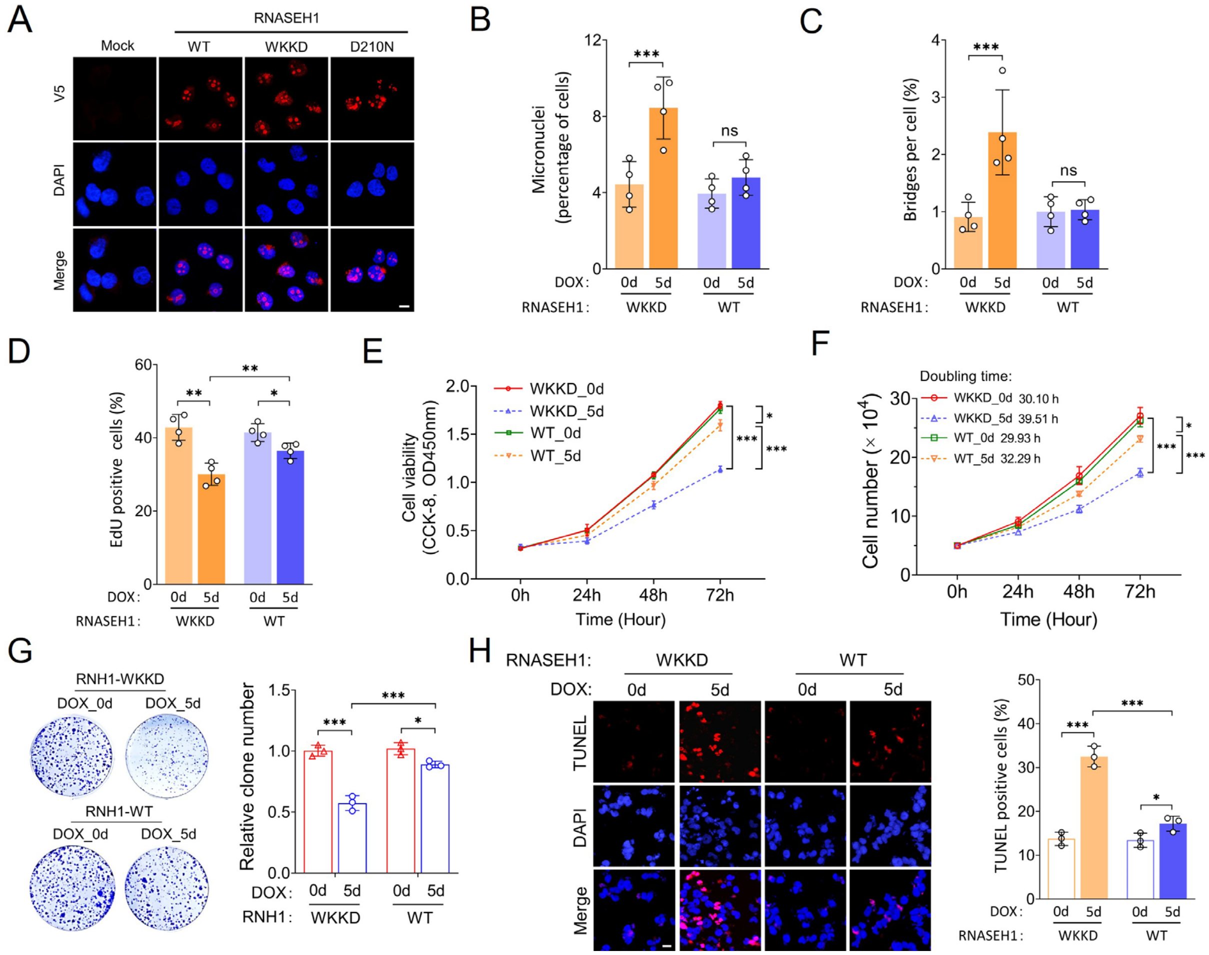


**Supplementary Figure 2: Effects of R-loop accumulation induced by LSH deficiency on cell proliferation and apoptosis in PC3 cells (corresponding to Figure 2). A** Immunofluorescence analysis of PC3 Tet-on LSH-shRNA cells revealed the localization of overexpressed wild-type (WT) and mutant (D210N and WKKD) RNASEH1, all tagged with V5. V5 is shown in red, DAPI in blue; The scale bar represents 10 μm. **B-C** Percentage of cells presenting micronuclei (**B**) and DNA bridge (**C**) in PC3 Tet-on LSH-shRNA cells overexpressing WT or WKKD RNASEH1. Cells were either left untreated (DOX: 0d) or treated with doxycycline for five days (DOX: 5d). Data are presented as a bar graph with mean ± s.d. (n = 4 biologically independent experiments). ***p < 0.001; n.s. indicates not significant, as determined by the two-tailed Mann–Whitney test. **D** The percentage of EdU-positive (EdU+) PC3 Tet-on LSH-shRNA cells overexpressing WT or WKKD RNASEH1, treated with or without doxycycline, is quantified and presented as a bar graph showing the mean ± s.d. (n = 4 biologically independent experiments). *p < 0.05; **p < 0.01, as determined by the two-tailed Mann–Whitney test. **E** Cell viability in Tet-on LSH-shRNA PC3 cells overexpressing WT or WKKD RNASEH1, treated with or without doxycycline, was assessed using the CCK-8 assay at 24, 48, and 72 hours. Data are presented as a curve chart with mean ± s.d. (n = 3 biologically independent experiments). *$p < 0.05$; ***$p < 0.001$, as determined by the One-way ANOVA with Tukey's multiple comparison test. **F** Proliferation curves of Tet-on LSH-shRNA PC3 cells overexpressing WT or WKKD RNASEH1, treated with or without doxycycline, were monitored over a 72-hour period. Fifty thousand cells were seeded, cultured, and counted at 24, 48, and 72 hours. Data are presented as a curve chart with mean ± s.d. (n = 3 biologically independent experiments). The doubling time was calculated for all groups. *$p < 0.05$; ***$p < 0.001$, as determined by the One-way ANOVA with Tukey's multiple comparison test. **G** Representative images of the colony formation assay using Tet-on LSH-shRNA PC3 cells overexpressing WT or WKKD RNASEH1, with or without doxycycline treatment. Cells were cultured in a 6-well dish for 14 days and stained with crystal violet to visualize colony growth. The bar graph shows the relative number of colonies compared to the DOX (-) group. Data are presented as the mean ± s.d. (n = 3 biologically independent experiments). *$p < 0.05$; ***$p < 0.001$, as determined by the two-tailed Mann–Whitney test. **H** Representative images of apoptotic cells analyzed by TUNEL assay (red), with nuclei counterstained by DAPI (blue), are shown. Tet-on LSH-shRNA PC3 cells overexpressing WT or WKKD RNASEH1 were treated with or without doxycycline. The scale bar represents 20 μm. Data are presented as a bar graph with mean ± s.d. (n = 3 biologically independent experiments). *$p < 0.05$; ***$p < 0.001$, as determined by the two-tailed Mann–Whitney test. Source data are provided as a Source Data file.

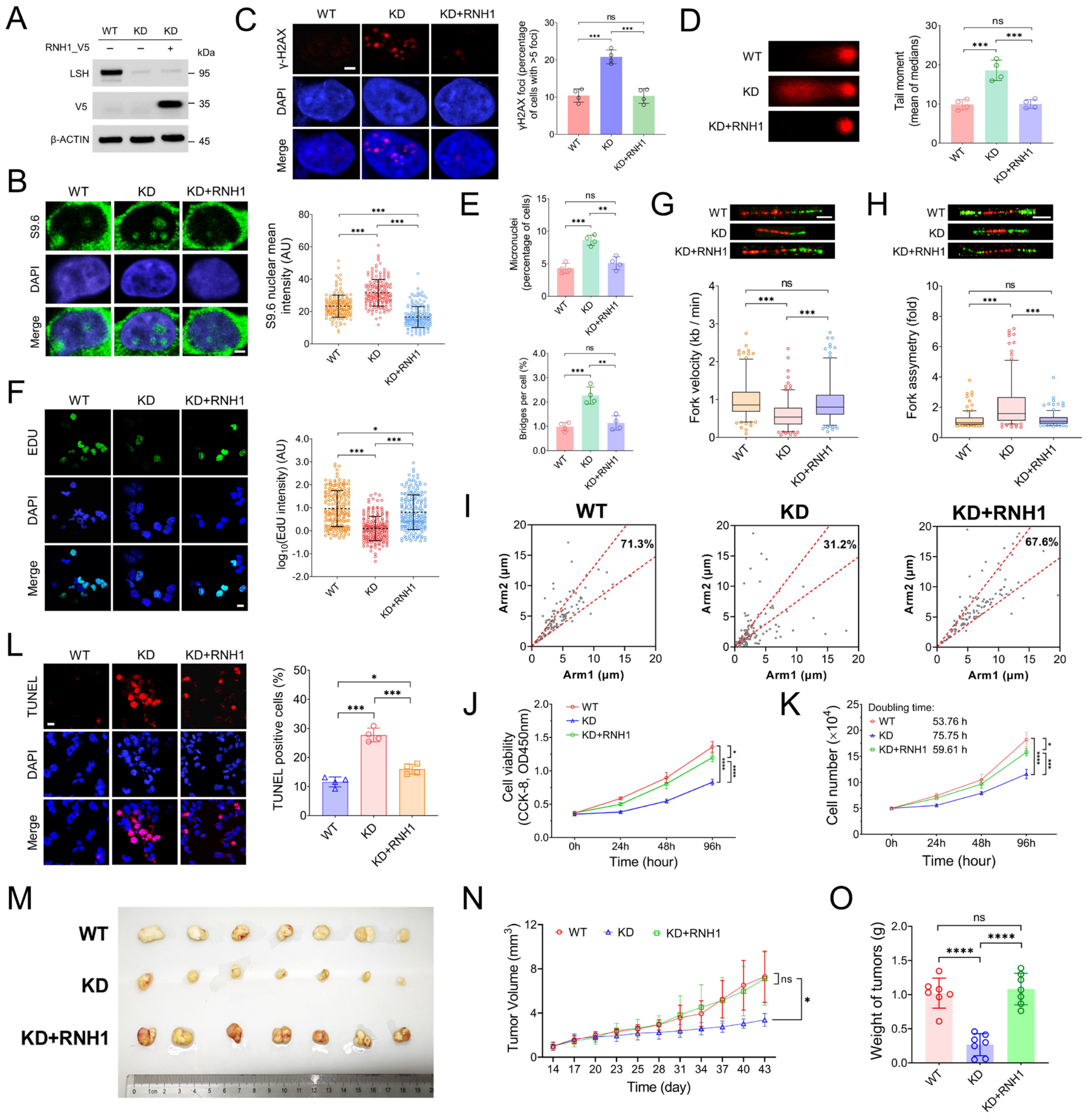


**Supplementary Figure 3: Impact of LSH Depletion on R-Loop accumulation, DNA damage, cell proliferation and apoptosis in LNCaP cells (corresponding to Figure 2). A** Western Blot Analysis of LNCaP cells confirmed LSH knockdown (KD) and overexpression of V5-Tagged RNASEH1_WT (KD+RNH1). LSH was detected using an anti-LSH antibody, RNH1_V5 was detected using an anti-V5 antibody, and β-Actin was used as the loading control. **B** Representative S9.6 immunostaining images evaluating R-loops level in WT, KD and KD+RNH1 LNCaP cells. The scale bar represents 5 μm. Data are presented as a scatter plot with mean ± s.d. ($n$ = 4 biologically independent experiments). ***p < 0.001, as determined by the One-way ANOVA with Tukey's multiple comparison test. AU denotes arbitrary units. **C** Representative images of γ-H2AX immunostaining and the percentage of cells with more than five γ-H2AX foci in WT, KD and KD+RNH1 LNCaP cells. The scale bar represents 5 μm. Data are presented as a bar graph with mean ± s.d. (n = 4 biologically independent experiments). ***p < 0.001; n.s. indicates not significant, as determined by the One-way ANOVA with Tukey's multiple comparison test. **D** Representative images of the alkaline comet assay and quantification of tail moment in WT, KD and KD+RNH1 LNCaP cells. Data are represented as mean ± s.d. (n = 4 biologically independent experiments). ***p < 0.001; n.s. indicates not significant, as determined by the One-way ANOVA with Tukey's multiple comparison test. **E** Percentage of cells presenting micronuclei and DNA bridge occurrence in WT, KD and KD+RNH1 LNCaP cells. Data are presented as a bar graph with mean ± s.d. (n = 4 biologically independent experiments). **p < 0.01; ***p < 0.001; n.s. indicates not significant, as determined by the One-way ANOVA with Tukey's multiple comparison test. **F** Representative images of EdU staining and quantification of nuclear EdU signal intensity in WT, KD, and KD+RNH1 LNCaP cells. The scale bar represents 10 μm. Signal intensity data are presented as a scatter plot with mean ± s.d. (n = 4 biologically independent experiments). *p < 0.05; ***p < 0.001, as determined by the One-way ANOVA with Tukey's multiple comparison test. AU denotes arbitrary units. **G-H** Quantification of replication fork velocity (**G**) and fork asymmetry (**H**) is presented using Tukey-style box plots. Representative images depict stretched DNA fibers from WT, KD and KD+RNH1 LNCaP cells. The cells were sequentially labeled with CldU (red) and IdU (green). The scale bar represents 10 μm. The central line of a Tukey-style box plot indicates the median value, while the boxes and whiskers represent the 25th to 75th percentiles and the 5th to 95th percentiles, respectively. Outliers are displayed as individual points. Data are representative of n = 4 biologically independent experiments. ***p < 0.001; n.s. indicates not significant, as determined by the One-way ANOVA with Tukey's multiple comparison test. **I** Scatter plots of the distances covered by right-moving and left-moving bi-directional replication forks with the CldU pulse in WT, KD and KD+RNH1 LNCaP cells. The central areas delimited with red lines contain bi-directional forks with less than a 25% length difference. The percentage of symmetric forks is indicated. Data are representative of n = 4 biologically independent experiments. **J** Cell viability in WT, KD and KD+RNH1 LNCaP cells was assessed using the CCK-8 assay at 24, 48, and 72 hours. Data are presented as a curve chart with mean ± s.d. (n = 3 biologically independent experiments). *p < 0.05; ****p < 0.0001, as determined by the One-way ANOVA with Tukey's multiple comparison test. **K** Proliferation curves of WT, KD and KD+RNH1 LNCaP cells were monitored over a 72-hour period. Fifty thousand cells were seeded, cultured, and counted at 24, 48, and 72 hours. The doubling time was calculated for all groups. Data are presented as a curve chart with mean ± s.d. (n = 3 biologically independent experiments). *p < 0.05; ***p < 0.001; ****p < 0.0001, as determined by the One-way ANOVA with Tukey's multiple comparison test. **L** Representative images of apoptotic cells analyzed by TUNEL assay (red), with nuclei counterstained by DAPI (blue) in WT, KD and KD+RNH1 LNCaP cells. The scale bar represents 20 μm. Data is presented as a bar graph with mean ± s.d. ($n = 3$ biologically independent experiments). *$p < 0.05$; ***$p < 0.001$, as determined by the One-way ANOVA with Tukey's multiple comparison test. **M** Representative images of tumorigenesis assay by subcutaneously injecting LSH WT, KD and KD+RNH1 LNCaP cells into M-NSG immunodeficient mice ($n = 7$ mice for each group). After 6 weeks, all mice were euthanized, and the tumors were harvested. **N** Two weeks after injecting LSH WT, KD and KD+RNH1 LNCaP cells, tumor volume was measured at 2–3 days intervals. The results are shown as a curve chart with mean ± s.d. ($n = 7$ mice for each group). *$p < 0.05$; n.s. indicates not significant, as determined by the One-way ANOVA with Tukey's multiple comparison test. **O** Upon reaching the experimental endpoint (6 weeks), all mice were euthanized, and the tumors were harvested for tumor weight measurement. Data is presented as a bar graph with mean ± s.d. ($n = 7$ mice for each group). ****$p < 0.0001$; n.s. indicates not significant, as determined by the One-way ANOVA with Tukey's multiple comparison test. Source data are provided as a Source Data file.

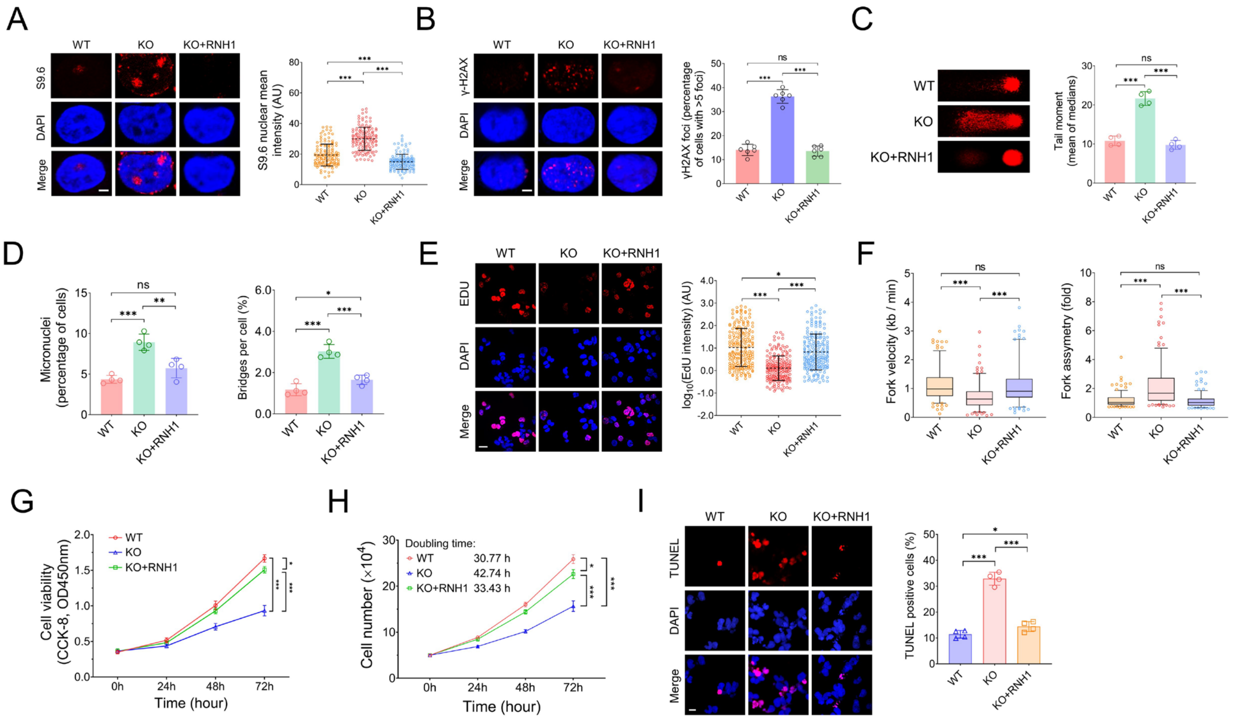


**Supplementary Figure 4: Impacts of LSH knockout on R-loops accumulation, DNA damage, cell proliferation, and apoptosis in PC3 cells (corresponding to Figure 3).** **A** Representative S9.6 immunostaining images evaluating R-loops level in WT, KO and KO+RNH1 PC3 cells. The scale bar represents 5 μm. Data is presented as a scatter plot with mean ± s.d. (n = 4 biologically independent experiments). ***p < 0.001, as determined by the One-way ANOVA with Tukey's multiple comparison test. AU denotes arbitrary units. **B** Representative images of γ-H2AX immunostaining and the percentage of cells with more than five γ-H2AX foci in WT, KO and KO+RNH1 PC3 cells. The scale bar represents 5 μm. Data is presented as a bar graph with mean ± s.d. (n = 6 biologically independent experiments). ***p < 0.001; n.s. indicates not significant, as determined by the One-way ANOVA with Tukey's multiple comparison test. **C** Representative images of the alkaline comet assay and quantification of tail moment in WT, KO and KO+RNH1 PC3 cells. Data are represented as mean ± s.d. (n = 4 biologically independent experiments). ***p < 0.001; n.s. indicates not significant, as determined by the One-way ANOVA with Tukey's multiple comparison test. **D** Percentage of cells presenting micronuclei and DNA bridge occurrence in WT, KO and KO+RNH1 PC3 cells. Data are presented as a bar graph with mean ± s.d. (n = 4 biologically independent experiments). *p < 0.05; **p < 0.01; ***p < 0.001; n.s. indicates not significant, as determined by the One-way ANOVA with Tukey's multiple comparison test. **E** Representative images of EdU staining and quantification of nuclear EdU signal intensity in WT, KO, and KO+RNH1 PC3 cells. The scale bar represents 10 μm. Signal intensity data are presented as a scatter plot with mean ± s.d. (n = 4 biologically independent experiments). *p < 0.05; ***p < 0.001, as determined by the One-way ANOVA with Tukey's multiple comparison test. AU denotes arbitrary units. **F** Quantification of replication fork velocity and asymmetry in WT, KO, and KO+RNH1 PC3 cells. The data are presented as a Tukey-style box plot, where the center line represents the median value, and the boxes and whiskers indicate the 25th to 75th percentiles and the 5th to 95th percentiles, respectively. Outliers are represented as individual points. The data are representative of n = 4 biologically independent experiments. ***p < 0.001; n.s. indicates not significant, as determined by the One-way ANOVA with Tukey's multiple comparison test. **G** Cell viability was assessed using the CCK-8 assay at 24, 48, and 72 hours in WT, KO and KO+RNH1 PC3 cells. Data are presented as a curve chart with mean ± s.d. (n = 4 biologically independent experiments). *p < 0.05; ***p < 0.001, as determined by the One-way ANOVA with Tukey's multiple comparison test. **H** Proliferation curves of WT, KO and KO+RNH1 PC3 cells were monitored over a 72-hour period. Fifty thousand cells were seeded, cultured, and counted at 24, 48, and 72 hours. The doubling time was calculated for all groups. Data are presented as a curve chart with mean ± s.d. (n = 3 biologically independent experiments). *p < 0.05; ***p < 0.001; ***p < 0.001, as determined by the One-way ANOVA with Tukey's multiple comparison test. **I** Representative images of apoptotic cells analyzed by TUNEL assay (red), with nuclei counterstained by DAPI (blue) in WT, KO and KO+RNH1 PC3 cells. The scale bar represents 20 μm. Data are presented as a bar graph with mean ± s.d. (n = 4 biologically independent experiments). *p < 0.05; ***p < 0.001, as determined by the One-way ANOVA with Tukey's multiple comparison test. Source data are provided as a Source Data file.

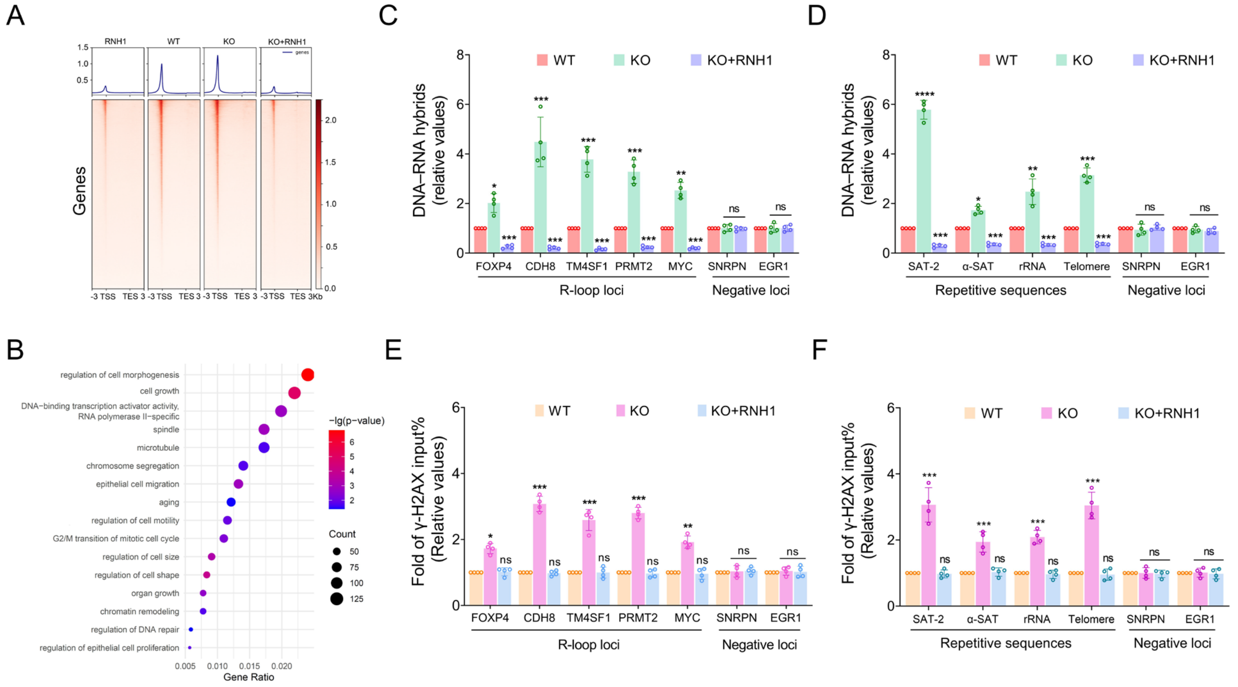


**Supplementary Figure 5: LSH deficiency results in an increased accumulation of R-loops in PC3 cells using R-loop CUT&Tag sequencing (corresponding to figure 3). A** The average R-loop CUT&Tag signals across the gene body region, from the TSS to TES sites, in LSH_WT and LSH_KO PC3 cells. Samples treated with the RNH1 enzyme and KO cells overexpressing RNH1 (KO+RNH1) were used as negative control groups to confirm the specificity of detected R-loop signals. Data are representative of n = 3 biologically independent experiments. TSS, transcription start site. TES, transcription end site. **B** Pathway analysis of genes with R-loop gains comparing LSH_KO and LSH_WT PC3 cells. Data represents n = 3 biologically independent experiments. **C-D** DNA-RNA hybrids CUT&Tag–qPCR analysis at R-loops loci (**C**) and repetitive sequences (**D**) among LSH_WT, LSH_KO and KO+RNH1 PC3 cells. Signal values normalized with respect to LSH_WT cells and plotted as mean ± s.d. (n = 4 biologically independent experiments). *p < 0.05; **p < 0.01; ***p < 0.001; n.s. indicates not significant, as determined by the One-way ANOVA with Tukey's multiple comparison test. **E-F** ChIP–qPCR analysis of γ-H2AX at R-loops loci (**E**) and repetitive sequences (**F**) among LSH_WT, LSH_KO and KO+RNH1 PC3 cells. Signal values normalized with respect to LSH_WT cells and plotted as mean ± s.d. (n = 4 biologically independent experiments). *p < 0.05; **p < 0.01; ***p < 0.001; n.s. indicates not significant, as determined by the One-way ANOVA with Tukey's multiple comparison test. Source data are provided as a Source Data file.

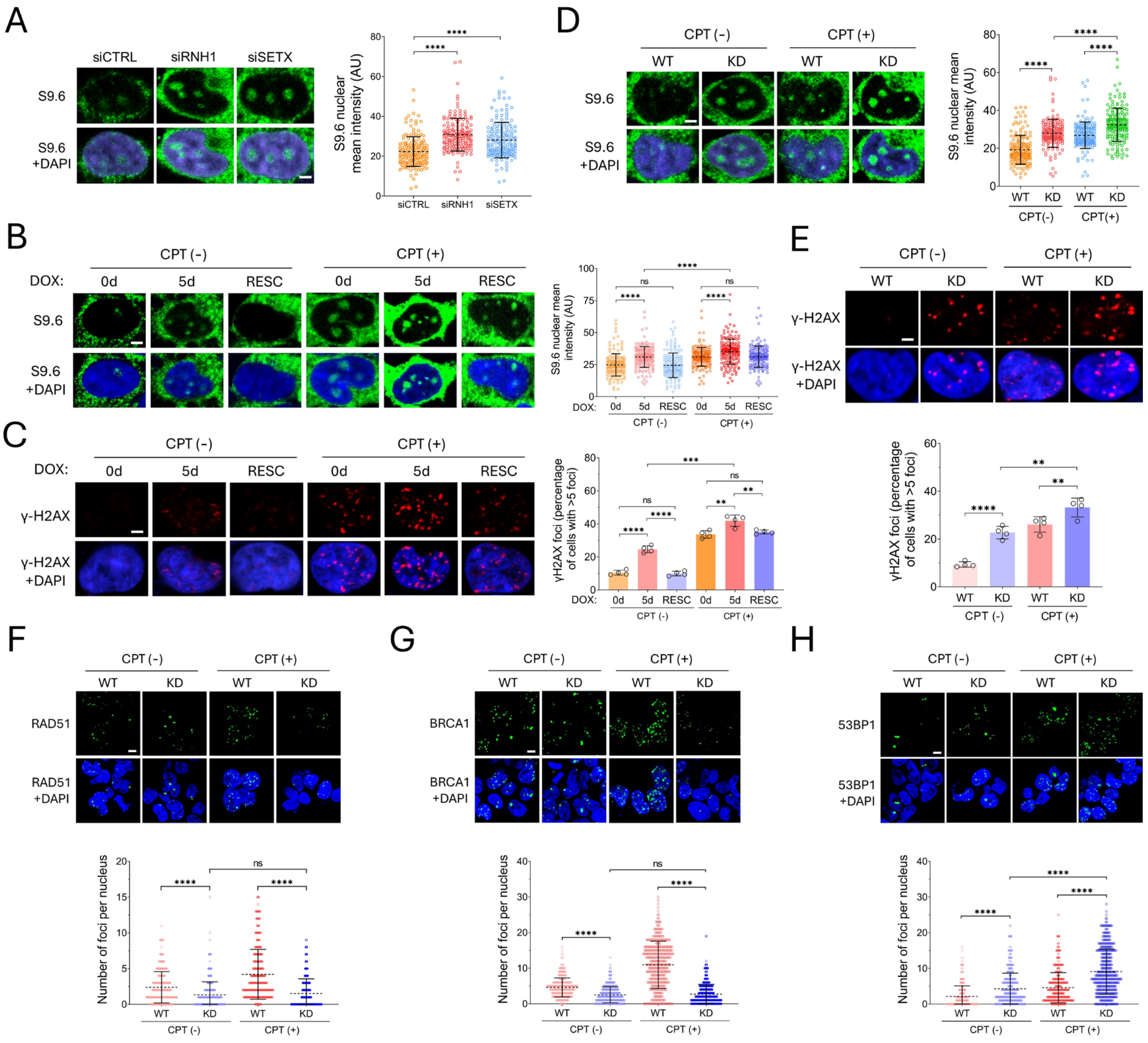


**Supplementary Figure 6: LSH deficiency impairs Rad51 filament formation at R-loop-induced DNA damage in PC3 and LNCaP cells (corresponding to Figure 4). A** Representative images of S9.6 immunostaining used to assess R-loop levels in PC3 cells treated with siRNA targeting RNH1 or SETX. The scale bar represents 5 μm. Data is presented as a scatter plot with mean ± s.d. (n = 4 biologically independent experiments). ****$p < 0.0001$, as determined by the One-way ANOVA with Tukey's multiple comparison test. **B** Representative images of S9.6 immunostaining to evaluate R-loop levels in PC3 Tet-on LSH-shRNA cells treated with DMSO or CPT (10 μM; 2 h). The cells were left untreated (Dox_0d), treated with for five days (Dox_5d) or subjected to a five-day doxycycline washout period (RESC). The scale bar represents 5 μm. Data is presented as a scatter plot with mean ± s.d. (n = 4 biologically independent experiments). ****$p < 0.0001$; n.s. indicates not significant, as determined by the One-way ANOVA with Tukey's multiple comparison test. **C** Representative images of γ-H2AX immunostaining and the percentage of cells with more than five γ-H2AX foci in PC3 Tet-on LSH-shRNA cells treated with DMSO or CPT (10 μM; 2 h). The cells were left untreated (Dox_0d), treated with for five days (Dox_5d) or subjected to a five-day doxycycline washout period (RESC). The scale bar represents 5 μm. Data are presented as a bar graph with mean ± s.d. (n = 4 biologically independent experiments). **p < 0.01; ***p < 0.001; ****p < 0.0001; n.s. indicates not significant, as determined by the One-way ANOVA with Tukey's multiple comparison test. **D** Representative images of S9.6 immunostaining to evaluate the R-loop levels in LSH WT and KD LNCaP cells treated with DMSO or CPT (10 μM; 2 h). The scale bar represents 5 μm. Data is presented as a scatter plot with mean ± s.d. (n = 4 biologically independent experiments). ****p < 0.0001, as determined by the two-tailed Mann–Whitney test. **E** Representative images of γ-H2AX immunostaining and the percentage of cells with more than five γ-H2AX foci in LSH WT and KD LNCaP cells treated with DMSO or CPT (10 μM; 2 h). The scale bar represents 5 μm. Data are presented as a bar graph with mean ± s.d. (n = 4 biologically independent experiments). **p < 0.01; ****p < 0.0001, as determined by the two-tailed Mann–Whitney test. **F-H** Representative immunofluorescence images showing RAD51 (**F**), BRCA1 (**G**), and 53BP1 (**H**) in LSH WT and KD LNCaP cells treated with DMSO or CPT (10 μM; 2 h). The scale bar represents 10 μm. The quantification of the number of RAD51, BRCA1, and 53BP1 foci per nucleus for each experimental condition was presented as a scatter plot with mean ± s.d. (n = 4 biologically independent experiments). ****p < 0.0001; n.s. indicates not significant, as determined by the two-tailed Mann–Whitney test.

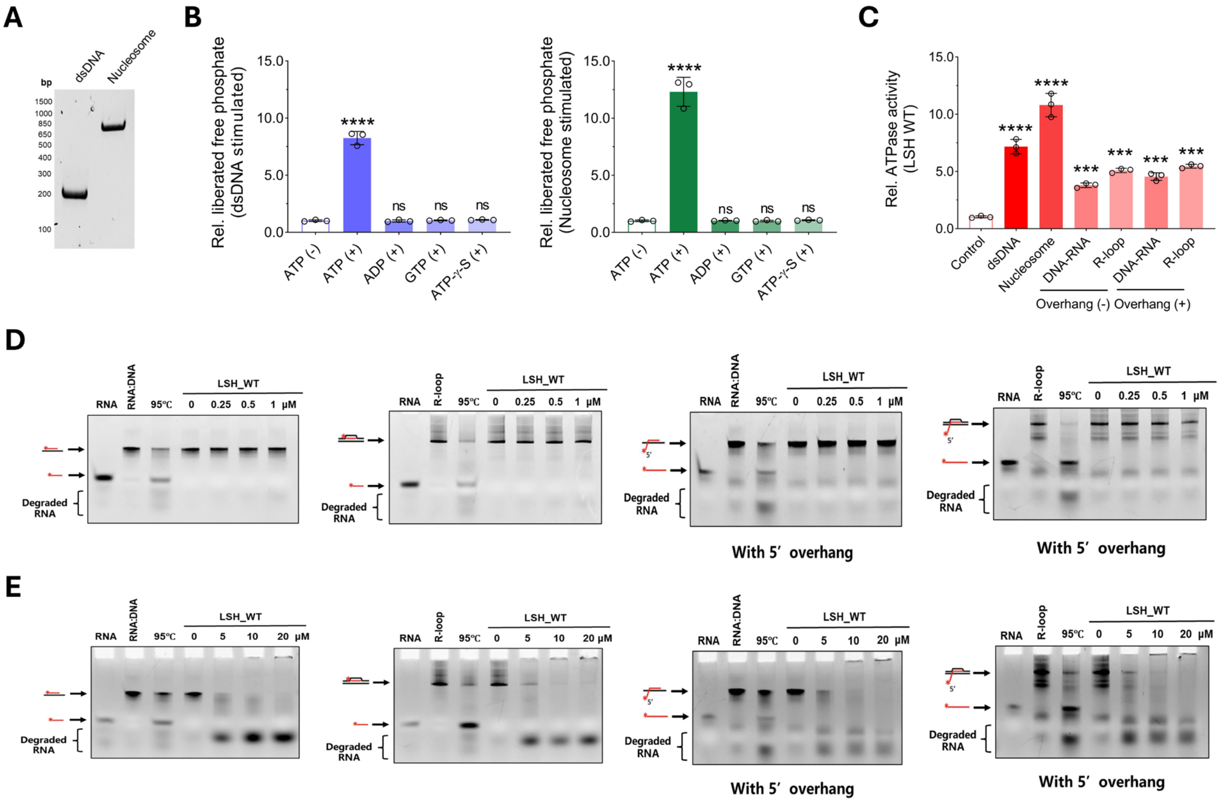


**Supplementary Figure 7: In vitro experiments revealed that LSH lacked the capacity to resolve either R-loops or RNA: DNA hybrids. A** Native PAGE and SYBR green staining showing the reconstituted mono-nucleosomes containing 208 bp double-stranded DNA (dsDNA) template. **B** LSH was ATP hydrolysis dependent, and ATP analogs including ADP, GTP, and ATP-γ-S failed to stimulate ATPase activity in the presence of dsDNA (left) or nucleosomes (right). Samples without ATP served as controls. The quantification of the liberated free phosphate was normalized to the controls. Data are presented as a bar graph with mean ± s.d. (n = 3 biologically independent experiments). ****p < 0.0001; n.s. indicates not significant, as determined by the two-tailed Mann–Whitney test. **C** Quantification of the relative ATPase activity stimulated by LSH_WT recombinant protein in the presence of dsDNA, nucleosomes, or various R-loops and RNA: DNA hybrids. Reactions without cofactors were served as controls. Data are presented as a bar graph with mean ± s.d. (n = 3 biologically independent experiments). ***p < 0.001; ****p < 0.0001, as determined by the two-tailed Mann–Whitney test. **D-E** In vitro unwinding assay indicated that LSH_WT recombinant protein, at low concentrations (**D**; 0 μM, 0.25 μM, 0.50 μM, and 1 μM) or at high concentrations (**E**; 0 μM, 5 μM, 10 μM, and 20 μM), lacked the capacity to unwind various R-loops (200 nM) or RNA: DNA hybrids (200 nM) in vitro, regardless of the presence of 5'-RNA overhangs. The RNA 5’-terminus was labeled with 6-FAM fluorescence; DNA is depicted in black, while RNA is shown in red. The samples were loaded onto a 15% polyacrylamide gel and visualized using the Cy2 channel.

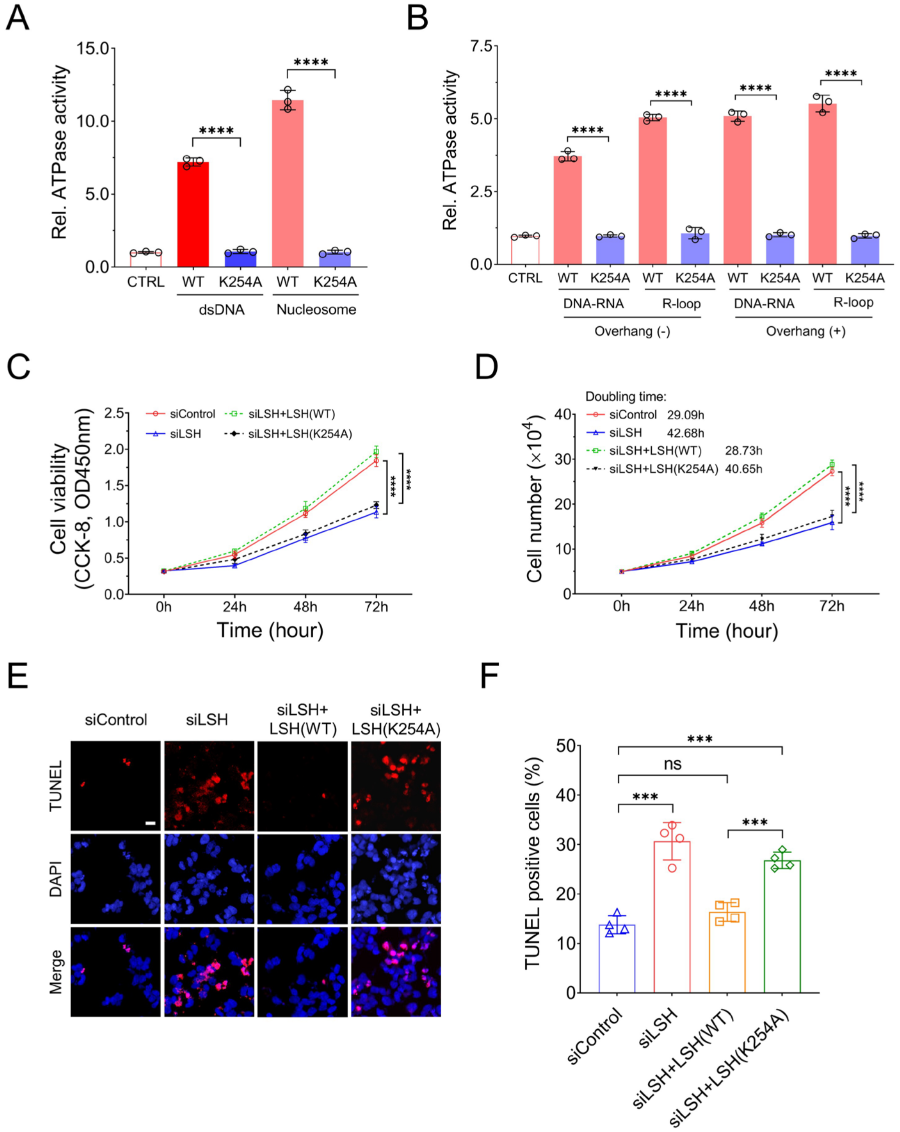


**Supplementary Figure 8: The mutation of LSH at the ATP-binding site is linked to cell proliferation and apoptosis in PC3 cells (corresponding to Figure 5). A** Comparisons of ATPase activity of LSH (WT) and LSH (K254A) recombinant protein in the presence of dsDNA or nucleosomes. Reactions without cofactors were served as controls. Data are presented as a bar graph with mean ± s.d. (n = 3 biologically independent experiments). ****$p < 0.0001$, as determined by the two-tailed Student's t test. **B** Comparisons of ATPase activity of LSH (WT) and LSH (K254A) recombinant protein in the presence of various R-loops and RNA: DNA hybrids. Reactions without cofactors were served as controls. Data are presented as a bar graph with mean ± s.d. (n = 3 biologically independent experiments). ****$p < 0.0001$, as determined by the two-tailed Student's t test. **C** Cell viability in PC3 cells was assessed using the CCK-8 assay at 0, 24, 48, and 72 hours. The cells overexpressing either exogenous LSH(WT)-Flag or LSH(K254A)-Flag were transfected with LSH 3' UTR siRNA or control siRNA. Data are presented as a curve chart with mean ± s.d. (n = 3 biologically independent experiments). ****$p < 0.0001$, as determined by the One-way ANOVA with Tukey's multiple comparison test. **D** Proliferation curves of PC3 cells treated as in panel C. The doubling time was calculated for all groups. Data are presented as a curve chart with mean ± s.d. (n = 3 biologically independent experiments). ****$p < 0.0001$, as determined by the One-way ANOVA with Tukey's multiple comparison test. **E-F** Representative images of apoptotic cells, analyzed by TUNEL assay (red), with nuclei counterstained by DAPI, are shown (**E**). The cells were treated as in panel C. The scale bar represents 20 μm. Data are presented as a bar graph showing the mean ± s.d. (**F**, n = 3 biologically independent experiments). ***$p < 0.001$; n.s. indicates not significant, as determined by the One-way ANOVA with Tukey's multiple comparison test. Source data are provided as a Source Data file.

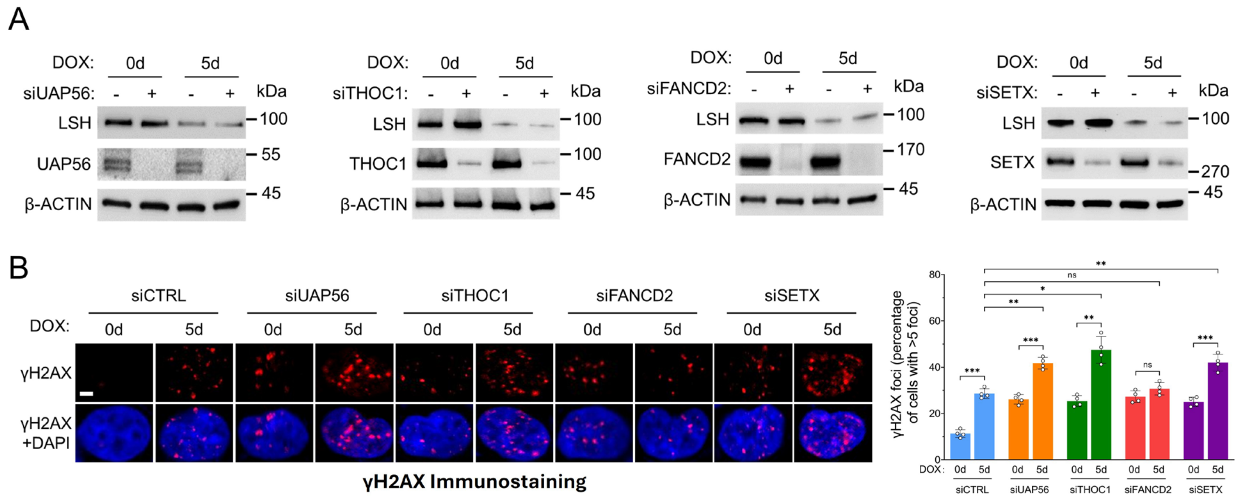


**Supplementary Figure 9: Analysis of DNA damage in relation to LSH and known R-loop-preventing factors (corresponding to Figure 6). A** Western blot analysis was performed on PC3 Tet-on LSH-shRNA cells following transient transfection with siRNAs targeting UAP56, THOC1, FANCD2, or SETX. The cells were either left untreated (DOX_0d) or treated with doxycycline for five days (DOX_5d). β-Actin was used as the loading control. **B** Representative images of γH2AX immunostaining in Tet-on LSH-shRNA PC3 cells following transient transfection with siRNAs targeting UAP56, THOC1, FANCD2, or SETX. The cells were either left untreated (DOX_0d) or treated with doxycycline for five days (DOX_5d). The scale bar represents 10 µm. The percentage of cells with more than five γ-H2AX foci are presented as a bar graph with mean ± s.d. (n = 4 biologically independent experiments). *$p < 0.05$; **$p < 0.01$; ***$p < 0.001$; n.s. indicates not significant, as determined by the two-tailed Mann–Whitney test. Source data are provided as a Source Data file.

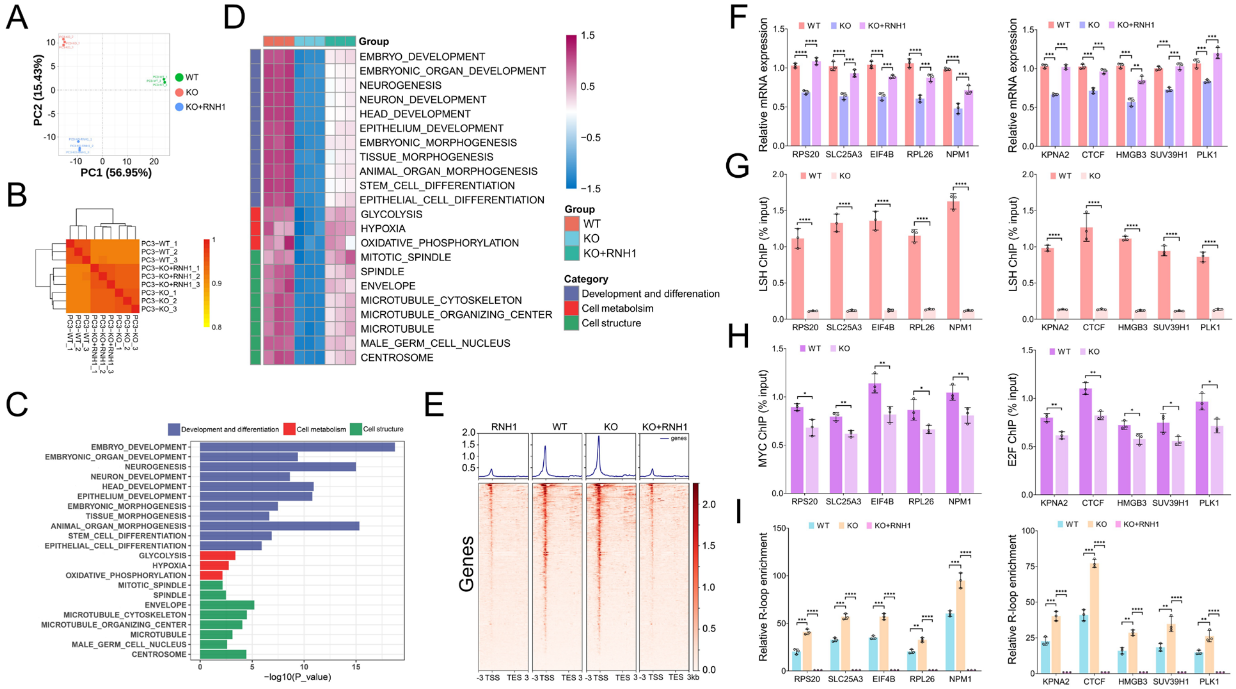


**Supplementary Figure 10: LSH-mediated R-loop accumulation is associated with the expression of MYC and E2F target genes (corresponding to Figure 7). A** Principal component analysis of the RNA-seq data was performed in LSH_WT, LSH_KO, and KO+RNH1 PC3 cells. Data represents n = 3 biologically independent experiments. **B** Unbiased clustering analysis of the RNA-seq data was performed in LSH_WT, LSH_KO, and KO+RNH1 PC3 cells. Data represents n = 3 biologically independent experiments. **C** Pathway analysis of the overlapping genes was performed, focusing on cell development and differentiation, cell metabolism, and cell structure. Data represents n = 3 biologically independent experiments. **D** Heat map displays the relative expression levels of overlapping genes in LSH_WT, LSH_KO, and KO+RNH1 PC3 cells, as outlined in the pathway analysis from panel C. Data are representative of n = 3 biologically independent experiments. **E** The average R-loop CUT&Tag signals across the gene body region, from TSS to TES sites, of overlapping genes were analyzed in LSH_WT, LSH_KO, and KO+RNH1 PC3 cells. Samples treated with RNH1 enzyme served as a negative control to confirm the specificity of the detected R-loop signals. Data are representative of n = 3 biologically independent experiments. TSS, transcription start site. TES, transcription end site. **F** RT-qPCR analysis of selected MYC (left panel) and E2F (right panel) target genes for LSH_WT, LSH_KO, and KO+RNH1 PC3 cells. Data are represented as a bar graph with mean ± s.d. (n = 3 biologically independent experiments). **p < 0.01; ***p < 0.001; ****p < 0.0001, as determined by the One-way ANOVA with Tukey's multiple comparison test. **G** ChIP–qPCR analysis of LSH at selected MYC (left panel) and E2F (right panel) target genes for LSH_WT and LSH_KO PC3 cells. Data are represented as a bar graph with mean ± s.d. (n = 3 biologically independent experiments).

****p < 0.0001, as determined by the two-tailed Mann–Whitney test. **H** ChIP–qPCR analysis of MYC (left panel) and E2F (right panel) at selected target genes for LSH_WT and LSH_KO PC3 cells. Data are represented as a bar graph with mean ± s.d. (n = 3 biologically independent experiments). *p < 0.05; **p < 0.01, as determined by the two-tailed Mann–Whitney test. **I** R-loop CUT&Tag–qPCR analysis of selected MYC (left panel) and E2F (right panel) target genes for LSH_WT, LSH_KO, and KO+RNH1 PC3 cells. Data are represented as a bar graph with mean ± s.d. (n = 3 biologically independent experiments). **p < 0.01; ***p < 0.001; ****p < 0.0001, as determined by the One-way ANOVA with Tukey's multiple comparison test. Source data are provided as a Source Data file.

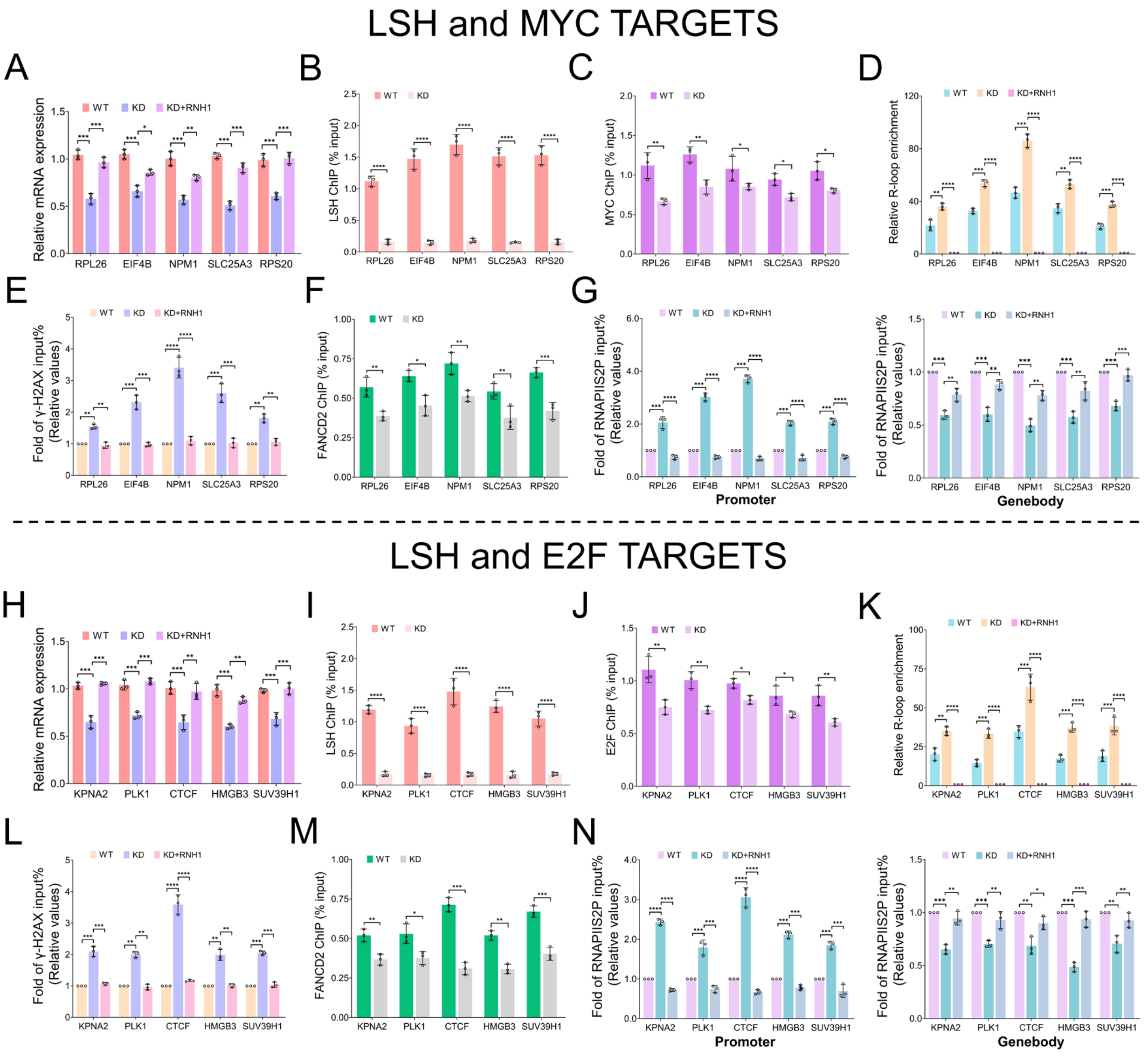


**Supplementary Figure 11: LSH-mediated R-loop accumulation is linked to MYC and E2F target genes expression through the modulation of RNA Polymerase II occupancy at promoter regions in LNCaP cells (corresponding to Figure 7). A** ChIP–qPCR analysis of LSH at the selected MYC target genes, as identified in the overlapping genes, was performed in LSH_WT and LSH_KD LNCaP cells. Data are represented as a bar graph with mean ± s.d. (n = 3 biologically independent experiments). *p < 0.05; **p < 0.01; ***p < 0.001, as determined by the One-way ANOVA with Tukey's multiple comparison test. **B** RT-qPCR analysis of the selected MYC target genes in LSH_WT and LSH_KD LNCaP cells. Data are represented as a bar graph with mean ± s.d. (n = 3 biologically independent experiments). ****p < 0.0001, as determined by the two-tailed Mann–Whitney test. **C** ChIP–qPCR analysis of MYC at the selected MYC target genes was performed in LSH_WT and LSH_KD LNCaP cells. Data are represented as a bar graph with mean ± s.d. (n = 3 biologically independent experiments). *p < 0.05; **p < 0.01, as determined by the two-tailed Mann–Whitney test. **D** R-loop CUT&Tag–qPCR analysis of the selected MYC target genes for LSH_WT, LSH_KD, and KD+RNH1 LNCaP cells. Data are represented as a bar graph with mean ± s.d. (n = 3 biologically independent experiments). $^{**}p < 0.01$; $^{***}p < 0.001$; $^{****}p < 0.0001$, as determined by the One-way ANOVA with Tukey's multiple comparison test. **E** ChIP–qPCR analysis of γ-H2AX at the selected MYC target genes for LSH_WT, LSH_KD, and KD+RNH1 LNCaP cells. Values were normalized to the WT group and presented as a bar graph with mean ± s.d. (n = 3 biologically independent experiments). $^{**}p < 0.01$; $^{***}p < 0.001$; $^{****}p < 0.0001$, as determined by the One-way ANOVA with Tukey's multiple comparison test. **F** ChIP–qPCR analysis of FANCD2 at the selected MYC target genes for LSH_WT and LSH_KD cells. Data are represented as a bar graph with mean ± s.d. (n = 3 biologically independent experiments). $^{*}p < 0.05$; $^{**}p < 0.01$; $^{***}p < 0.001$, as determined by the two-tailed Mann–Whitney test. **G** ChIP–qPCR analysis of RNAPIIS2P at the promoter and gene body regions of the selected MYC target genes for LSH_WT, LSH_KD, and KD+RNH1 LNCaP cells. Values were normalized to the WT group and presented as a bar graph with mean ± s.d. (n = 3 biologically independent experiments). $^{**}p < 0.01$; $^{***}p < 0.001$, as determined by the One-way ANOVA with Tukey's multiple comparison test. **H** ChIP–qPCR analysis of LSH at the selected E2F target genes, as identified in the overlapping genes, was performed in LSH_WT and LSH_KD LNCaP cells. Data are represented as a bar graph with mean ± s.d. (n = 3 biologically independent experiments). $^{**}p < 0.01$; $^{***}p < 0.001$, as determined by the One-way ANOVA with Tukey's multiple comparison test. **I** RT-qPCR analysis of the selected E2F target genes in LSH_WT and LSH_KD LNCaP cells. Data are represented as a bar graph with mean ± s.d. (n = 3 biologically independent experiments). $^{****}p < 0.0001$, as determined by the two-tailed Mann–Whitney test. **J** ChIP–qPCR analysis of E2F at the selected E2F target genes was performed in LSH_WT and LSH_KD LNCaP cells. Data are represented as a bar graph with mean ± s.d. (n = 3 biologically independent experiments). $^{*}p < 0.05$; $^{**}p < 0.01$, as determined by the two-tailed Mann–Whitney test. **K** R-loop CUT&Tag–qPCR analysis of the selected E2F target genes for LSH_WT, LSH_KD, and KD+RNH1 LNCaP cells. Data are represented as a bar graph with mean ± s.d. (n = 3 biologically independent experiments). $^{**}p < 0.01$; $^{***}p < 0.001$; $^{****}p < 0.0001$, as determined by the One-way ANOVA with Tukey's multiple comparison test. **L** ChIP–qPCR analysis of γ-H2AX at the selected E2F target genes for LSH_WT, LSH_KD, and KD+RNH1 LNCaP cells. Values were normalized to the WT group and presented as a bar graph with mean ± s.d. (n = 3 biologically independent experiments). $^{**}p < 0.01$; $^{***}p < 0.001$; $^{****}p < 0.0001$, as determined by the One-way ANOVA with Tukey's multiple comparison test. **M** ChIP–qPCR analysis of FANCD2 at the selected E2F target genes for LSH_WT and LSH_KD cells. Data are represented as a bar graph with mean ± s.d. (n = 3 biologically independent experiments). $^{*}p < 0.05$; $^{**}p < 0.01$; $^{***}p < 0.001$, as determined by the two-tailed Mann–Whitney test. **N** ChIP–qPCR analysis of RNAPIIS2P at the promoter and gene body regions of the selected E2F target genes for LSH_WT, LSH_KD, and KD+RNH1 LNCaP cells. Values were normalized to the WT group and presented as a bar graph with mean ± s.d. ($n = 3$ biologically independent experiments). **$p < 0.01$; ***$p < 0.001$; ****$p < 0.0001$, as determined by the One-way ANOVA with Tukey's multiple comparison test. Source data are provided as a Source Data file.

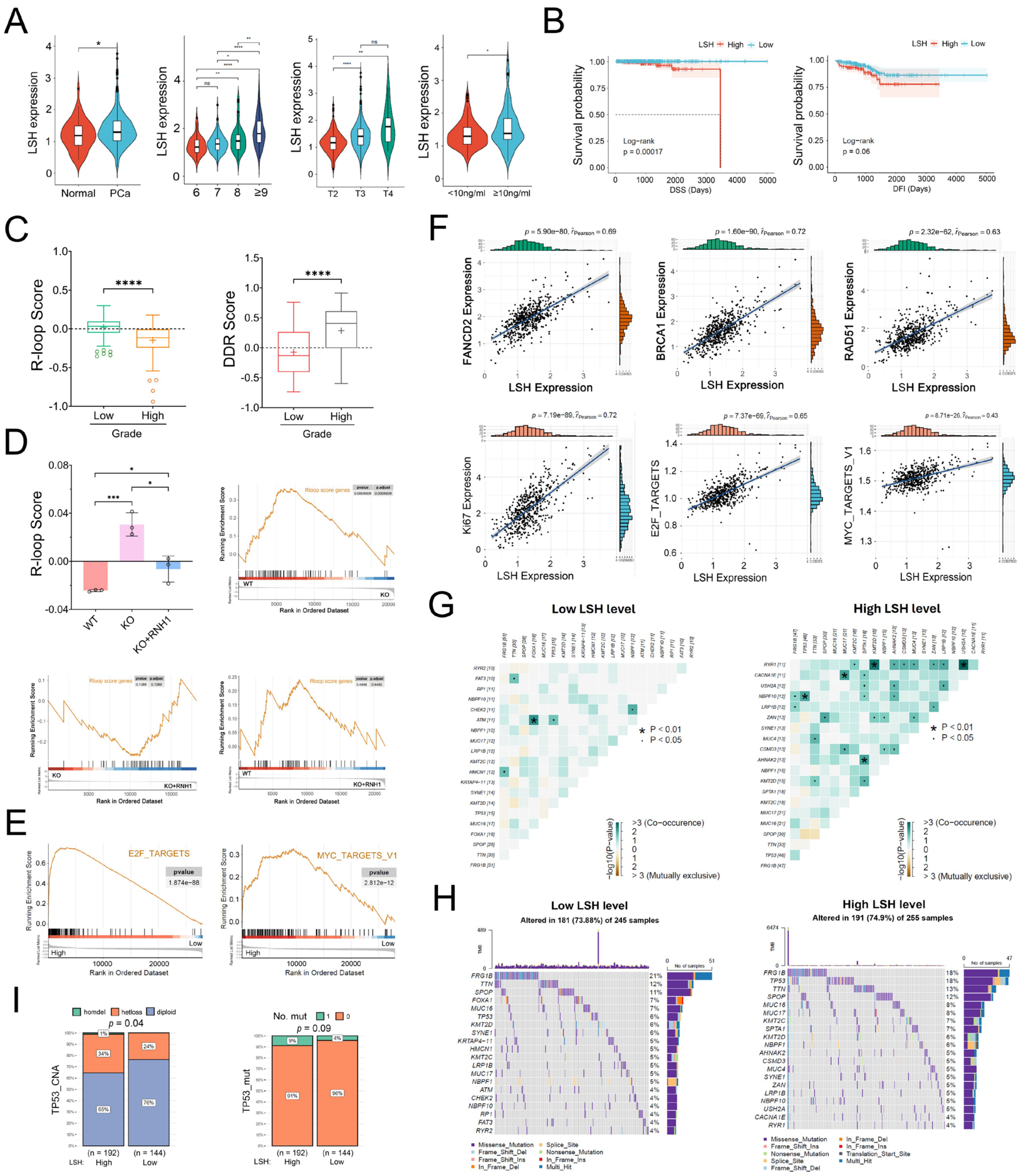


**Supplementary Figure 12: Clinical relevance of LSH expression in relation to R-loop accumulation in prostate cancer (corresponding to Figure 8). A** Comparisons of LSH expression are presented using violin plots across subgroups based on tissue type (normal tissue vs. prostate cancer), Gleason score, pathological stage, and PSA value utilizing prostate cancer TCGA RNA-seq data. Endpoints depict minimum and maximum values, quartiles depicted by thin black lines, median depicted by thick black lines. *p < 0.05; **p < 0.01; ****p < 0.0001; ns indicates not significant, as determined by the two-tailed Student's t test or the One-way ANOVA with Tukey's multiple comparison test. **B** High LSH expression was significantly associated with a shorter survival rate, as indicated by Disease-specific survival (DSS) and Disease-free interval (DFI) data. The p-values are shown in the panel. **C** Prostate cancer patients with high Gleason score exhibited lower R-loop scores and higher DDR scores. ****p < 0.0001, as determined by the two-tailed Student's t test. **D** The bar graph indicates that LSH KO cells exhibit higher R-loop scores compared to LSH WT cells, while KO+RNH1 cells partially restore the scores. Meanwhile, enrichment plots for genes associated with R-loop scores were generated. LSH KO cells exhibited a reduced expression of R-loop score-related genes compared to LSH WT cells, while KO+RNH1 cells showed a restoration in the enrichment of these genes. *p < 0.05; ***p < 0.001, as determined by the One-way ANOVA with Tukey's multiple comparison test. **E** Enrichment plots for E2F and MYC targets were generated using prostate cancer TCGA RNA-seq data, which was classified into high and low LSH expression groups. The p-values are shown in the panel. **F** Pearson correlations between LSH expression and the levels of FANCD2, BRCA1, RAD51, and Ki67 were analyzed in prostate cancer, along with correlations between LSH expression and E2F and MYC target genes. The p-values are shown in the panel. **G** Heatmap displaying mutually exclusive or co-occurring mutations among the top mutated genes in low and high LSH expression groups. *p < 0.01, • p < 0.05. **H** Oncoplots showing the top 20 mutated genes in low and high LSH expression groups using prostate cancer TCGA RNA-seq data. **I** Comparisons of mutations in TP53 gene in high or low LSH expression groups for prostate cancer patients. The p-values are shown in the panel. Significance was determined by the two-tailed Student's t test. Source data are provided as a Source Data file.

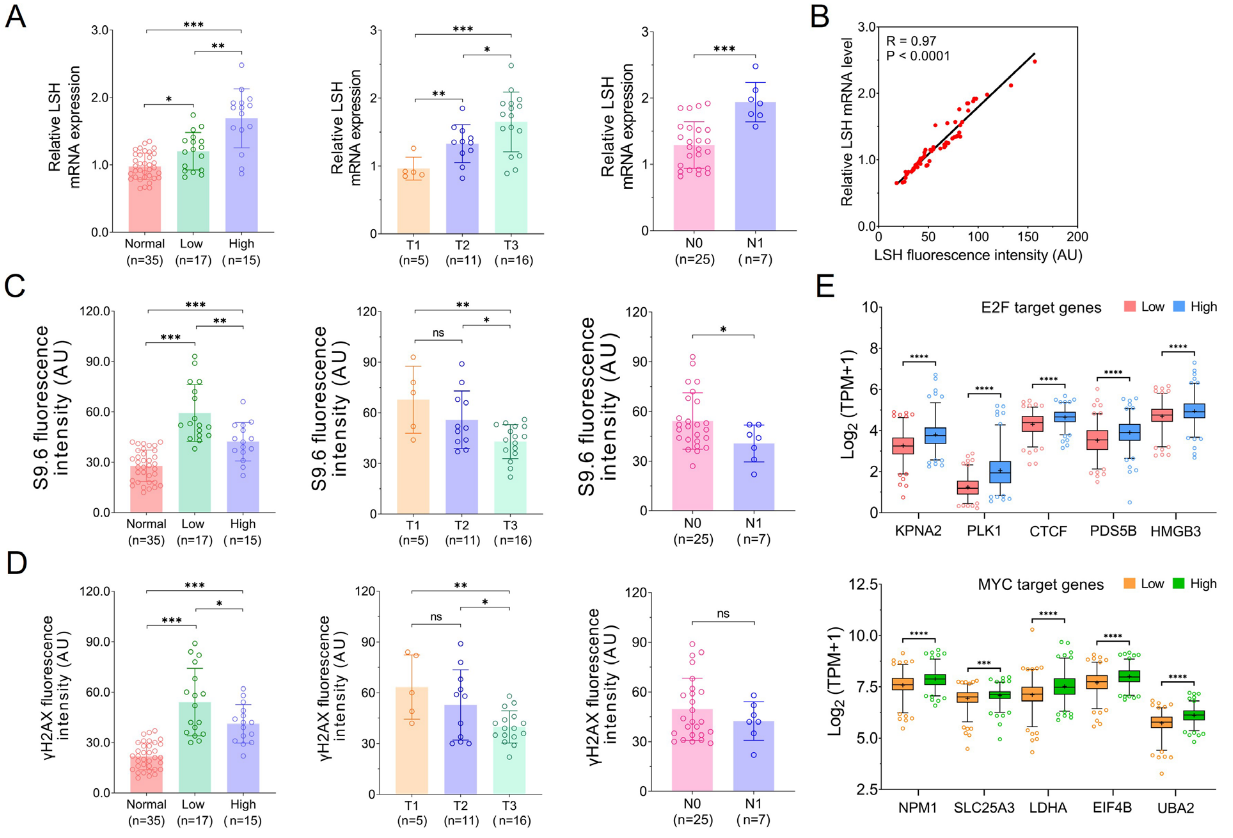


**Supplementary Figure 13: Clinical relevance of LSH mRNA expression, R-loop accumulation and DNA damage using real-world collected prostate cancer samples (corresponding to Figure 8). A** Quantification of relative LSH mRNA expression across subgroups based on the Gleason score (low vs. high), clinical stage (T) and lymph node involvement (N) using real-world collected prostate cancer samples. Data are presented as a bar graph with mean ± s.d. *$p < 0.05$; **$p < 0.01$; ***$p < 0.001$, as determined by the two-tailed Student's t test or the One-way ANOVA with Tukey's multiple comparison test. **B** Pearson correlations between LSH fluorescence intensity and relative LSH mRNA level. The R value and p-value are shown in the panel. AU denotes arbitrary units. **C** Quantification of R-loop accumulation analyzed by S9.6 fluorescence intensity across subgroups based on the Gleason score (low vs. high), clinical stage (T) and lymph node involvement (N) using real-world collected prostate cancer samples. Data are presented as a bar graph with mean ± s.d. *$p < 0.05$; **$p < 0.01$; ***$p < 0.001$; ns indicates not significant, as determined by the two-tailed Student's t test or the One-way ANOVA with Tukey's multiple comparison test. **D** Quantification of DNA damage analyzed by γ-H2AX immunostaining across subgroups based on the Gleason score (low vs. high), clinical stage (T) and lymph node involvement (N) using real-world collected prostate cancer samples. Data are presented as a bar graph with mean ± s.d. *$p < 0.05$, **$p < 0.01$ and ***$p < 0.001$; ns indicates not significant, as determined by the two-tailed Student's t test or the One-way ANOVA with Tukey's multiple comparison test. **E** Analysis of selected E2F and MYC target genes expression using prostate cancer RNA-seq data from TCGA database. Data was classified into low and high LSH expression groups. The E2F and MYC target genes expression are presented as a box plot, where the center line represents the median value, and the boxes and whiskers indicate the 25th to 75th and 5th to 95th percentiles, respectively. Outliers are shown as individual points. ***$p < 0.001$; ****$p < 0.0001$, as determined by the two-tailed Student's t test. Source data are provided as a Source Data file.

**Supplementary Table 1. siRNA sequences used in this study.**

| siRNA | Strand | Sequence (5’-3’) |
|---|---|---|
| siControl | Sense | UUCUCCGAACGUGUCACGUdTdT |
| | Anti-sense | ACGUGACACGUUCGGAGAAdTdT |
| siUAP56 | Sense | GUCUAUCAAGAAGGAUGAAdTdT |
| | Anti-sense | UUCAUCCUUCUUGAUAGACdTdT |
| siTHOC1 | Sense | GACGAGGAAGCUCCAACAAdTdT |
| | Anti-sense | UUGUUGGAGCUUCCUCGUCdTdT |
| siFANCD2 | Sense | GGCUUGACAGAGUUGUGGAdTdT |
| | Anti-sense | UCCACAACUCUGUCAAGCCdTdT |
| siSETX | Sense | GCAUGAGGUUUUAUGGGAAtt |
| | Anti-sense | UUCCCAUAAAACCUCAUGCtt |
| siRNaseH1 | Sense | GGAUGGAGAUGGACAUGAAtt |
| | Anti-sense | UUCAUGUCCAUCUCCAUCCtt |
| siLSH | Sense | GAACAAAGAAGUAUCCAUAUUtt |
| | Anti-sense | AAUAUGGAUACUUCUUUGUUCtt |

**Supplementary Table 2. Clinical characteristics of the real-world patients with prostate cancer included in the study.**

| Patient ID | Age | G1 | G2 | GS | Pathology T stage | Pathology N stage | Pathology M stage |
|---|---|---|---|---|---|---|---|
| 1 | 75 | 4 | 4 | 8 | 3 | 1 | 0 |
| 2 | 74 | 3 | 4 | 7 | 2 | 0 | 0 |
| 3 | 82 | 4 | 3 | 7 | 1 | 0 | 0 |
| 4 | 68 | 4 | 5 | 9 | 3 | 1 | 0 |
| 5 | 71 | 3 | 5 | 8 | 1 | 0 | 0 |
| 6 | 64 | 3 | 3 | 6 | 1 | 0 | 0 |
| 7 | 77 | 4 | 3 | 7 | 3 | 0 | 0 |
| 8 | 65 | 4 | 4 | 8 | 3 | 0 | 0 |
| 9 | 69 | 4 | 4 | 8 | 3 | 0 | 0 |
| 10 | 67 | 3 | 5 | 8 | 2 | 0 | 0 |
| 11 | 74 | 4 | 5 | 9 | 3 | 1 | 0 |
| 12 | 76 | 3 | 4 | 7 | 1 | 0 | 0 |
| 13 | 74 | 3 | 5 | 8 | 2 | 0 | 0 |
| 14 | 73 | 4 | 3 | 7 | 2 | 0 | 0 |
| 15 | 68 | 3 | 4 | 7 | 3 | 0 | 0 |
| 16 | 73 | 4 | 3 | 7 | 2 | 0 | 0 |
| 17 | 66 | 3 | 5 | 8 | 3 | 0 | 0 |
| 18 | 78 | 4 | 4 | 8 | 3 | 0 | 0 |
| 19 | 72 | 5 | 4 | 9 | 3 | 1 | 0 |
| 20 | 75 | 3 | 4 | 7 | 2 | 0 | 0 |
| 21 | 61 | 4 | 4 | 8 | 3 | 1 | 0 |
| 22 | 71 | 4 | 5 | 9 | 3 | 1 | 0 |
| 23 | 65 | 3 | 4 | 7 | 1 | 0 | 0 |
| 24 | 67 | 3 | 3 | 6 | 2 | 0 | 0 |
| 25 | 69 | 3 | 4 | 7 | 3 | 0 | 0 |
| 26 | 73 | 4 | 4 | 8 | 3 | 0 | 0 |
| 27 | 73 | 3 | 4 | 7 | 2 | 1 | 0 |
| 28 | 72 | 3 | 4 | 7 | 3 | 0 | 0 |
| 29 | 77 | 3 | 3 | 6 | 2 | 0 | 0 |
| 30 | 66 | 3 | 3 | 6 | 2 | 0 | 0 |
| 31 | 66 | 3 | 4 | 7 | 2 | 0 | 0 |
| 32 | 74 | 4 | 5 | 9 | 3 | 0 | 0 |

### Supplementary Table 3. Antibodies used in this study.

| Antibody | Company | Catalog # | Dilution for WB | Dilution for IF/PLA | CO-IP (μg) | ChIP/CUT&TAG (μg) |
|---|---|---|---|---|---|---|
| LSH | Cell signaling | 7998S | 1: 1000 | 1: 400 | 3 μg | 3 μg |
| γ-H2AX | Cell signaling | 9718S | | 1: 400 | | 3 μg |
| S9.6 | Sigma | MABE1095 | | 1: 400 | 3 μg | 3 μg |
| FANCD2 | Santa Cruz | sc-20022 | 1: 100 | 1: 50 | | 3 μg |
| BLM | Abcam | ab2179 | | 1: 400 | | |
| RNaseH1 | Santa Cruz | sc-376326 | 1: 100 | | | |
| SETX | Santa Cruz | sc-100319 | 1: 100 | | | |
| UAP56 | Proteintech | 14798-1-AP | 1: 500 | | | |
| THOC1 | Santa Cruz | sc-514123 | 1: 100 | | | |
| 53BP1 | Santa Cruz | sc-517281 | | 1: 100 | | |
| RAD51 | Santa Cruz | sc-377467 | | 1: 100 | | |
| BRCA1 | Santa Cruz | sc-6954 | | | | |
| PCNA | Santa Cruz | sc-56 | | 1: 200 | | |
| RNAPIIS2P | Abcam | ab193468 | | 1: 200 | | 3 μg |
| γ-RPA32 S4/8P | Bethyl laboratories | A300-245A | | 1: 200 | | |
| MYC | Santa Cruz | sc-40 | 1: 100 | | | 3 μg |
| E2F3 | Santa Cruz | sc-56665 | 1: 100 | | | 3 μg |
| LAMIN B1 | Abcam | ab16048 | 1: 500 | | | |
| Flag | Thermo Fisher | MA1-91878 | 1: 1000 | | | |
| V5 | Thermo Fisher | R960-25 | 1: 1000 | 1: 200 | | |
| β-ACTIN | Cell signaling | 4967S | 1: 1000 | | | |
| GAPDH | Cell signaling | 2118S | 1: 1000 | | | |
| Rabbit IgG | Millipore | 12-370 | | | | 3 μg |
| Mouse IgG | Millipore | 12-371 | | | | 3 μg |
| anti-rabbit IgG, HRP | Abcam | ab6721 | 1: 2500 | | | |
| anti-mouse IgG, HRP | Abcam | ab6728 | 1: 2500 | | | |
| anti-rabbit IgG, 488 | Abcam | ab150077 | | 1: 500 | | |
| anti-rabbit IgG, 594 | Abcam | ab150080 | | 1: 500 | | |
| anti-mouse IgG, 488 | Abcam | ab150113 | | 1: 500 | | |
| anti-mouse IgG, 594 | Abcam | ab150116 | | 1: 500 | | |

### Supplementary Table 4. Primers used in this study.

| Primer | Direction | Sequence | Application |
|---|---|---|---|
| FOXP4 | F | TTGGTGCACGTGGTTTTCTC | ChIP/CUT&TAG |
| | R | CCTAAAGCAGGTGCAGCAACT | |
| CDH8 | F | TGAGGACACAGTGAGAAGTTGATTG | ChIP/CUT&TAG |
| | R | CATGCCAGTGTGATCGGATTC | |
| TM4SF1 | F | TGGTTGAGGCTTTGAAAGACAGT | ChIP/CUT&TAG |
| | R | CAGATTGGAAGCTGTCCAGACA | |
| PRMT2 | F | GCATTAGCGCCACCCATTT | ChIP/CUT&TAG |
| | R | GGAAGCAGAATGATGACGTTTCT | |
| SAT-2 | F | CATCGAATGGAAATGAAAGGAGTC | ChIP/CUT&TAG |
| | R | ACCATTGGATGATTGCAGTCAA | |
| α-SAT | F | TCATTCCCACAAACTGCGTTG | ChIP/CUT&TAG |
| | R | TCCAACGAAGGCCACAAGA | |
| rRNA | F | GGATGCGTGCATTTATCAGA | ChIP/CUT&TAG |
| | R | GATCGGCCCGAGGTTATCTA | |
| Telomere | F | GTCCGTCCGTGAAATTGCG | ChIP/CUT&TAG |
| | R | GGTCCAAACGAGTCTCCGTC | |
| SNRPN | F | GCCAAATGAGTGAGGATGGT | ChIP/CUT&TAG |
| | R | TCCTCTCTGCCTGACTCCAT | |
| EGR1 | F | GAACGTTCAGCCTCGTTCTC | ChIP/CUT&TAG |
| | R | GGAAGGTGGAAGGAAACACA | |
| RPS20 | F | GCCACCCAGTCCTGATACCT | ChIP/CUT&TAG |
| | R | CAGCTCAGTTGACCATTCGG | |
| SLC25A3 | F | GCCTGTAATCCCAGCACATT | ChIP/CUT&TAG |
| | R | TCTTACCCTGCTCGCTTTGT | |
| EIF4B | F | AGTTGCTCCGGTTCAAACAC | ChIP/CUT&TAG |
| | R | AGAAGACAGGATGCGCAGTT | |
| RPL26 | F | AGATTCAGTTCTGGGTGGTGA | ChIP/CUT&TAG |
| | R | CAGGCCGAGCATTTAGAAAC | |
| NPM1 | F | AAGTCACCCGCTTTCTTTCA | ChIP/CUT&TAG |
| | R | TCGTTACCCCAAAGTTCAGG | |
| KPNA2 | F | TCCTGATGCTCTGTTGACCA | ChIP/CUT&TAG |
| | R | TTCAAATGCAGCAGAATTGC | |
| CTCF | F | TGGCCATGGAATTTCTCTTC | ChIP/CUT&TAG |
| | R | GACCGTTCCGTTGTGAGAAT | |
| HMGB3 | F | GTTGGCCCAAGGTACTCAAA | ChIP/CUT&TAG |
| | R | TTCTCTGCAGCAAAGCTCAA | |
| SUV39H1 | F | TCACTCTTGTGGGTTGGACA | ChIP/CUT&TAG |
| | R | AGGCAGGCTGTCATTCAGTT | |
| PLK1 | F | ATGGTGGTGCATGCTTGTAA | ChIP/CUT&TAG |

| | | | |
|---|---|---|---|
| | R | GGCAGGTCATAGCCAAATGT | |
| RPS20 | F | GACCAGTTCGAATGCCTACCA | RT |
| | R | AAGTGTACTGCTGGCCCTC | |
| SLC25A3 | F | GTGGCACAACACATACAGCA | RT |
| | R | AGTACGTTCAAAGCAGGCGA | |
| EIF4B | F | AGCAGAAGTAAGTCAGACCAGG | RT |
| | R | CTTGTAGGGGACTGCTGCTC | |
| RPL26 | F | GACTTCCGACCGAAGCAAGA | RT |
| | R | CATTAGCCTTTTCCCGCTGC | |
| NPM1 | F | AACGGTCAGTTTAGGGGCTG | RT |
| | R | GGAACCTTGCTACCACCTCC | |
| KPNA2 | F | TCCAAGCTACTCAAGCTGCC | RT |
| | R | CTATCGGGGGTGCAGGATTC | |
| CTCF | F | ACGCCAGTGTAGAAGTCAGC | RT |
| | R | ACAGCATCACAGTAACGGCA | |
| HMGB3 | F | GGCAAAGGCAGATAAAGTGC | RT |
| | R | CCACATCTCACCCAGCTTTT | |
| SUV39H1 | F | GGCAACATCTCCCACTTTGT | RT |
| | R | CAATACGGACCCGCTTCTTA | |
| PLK1 | F | AAGAGATCCCGGAGGTCCTA | RT |
| | R | GCTGCGGTGAATGGATATTT | |
| PDS5B | F | CAGTGGCCTGAGGAAAAGAG | RT |
| | R | GCGTCCTACACGGACATTTT | |
| LDHA | F | TGGCAGCCTTTTCCTTAGAA | RT |
| | R | CGCTTCCAATAACACGGTTT | |
| UBA2 | F | AATCAATGGCAGGGAACATT | RT |
| | R | TGACGTCGGCATTATTTGAA | |
| LSH | F | GAACCCAGGAGGAACGTCAA | RT |
| | R | ACTCCCTGATTAGACGGCAC | |
| GAPDH | F | GAGTCAACGGATTTGGTCGT | RT |
| | R | TTGATTTTGGAGGGATCTCG | |

**Supplementary Table 5. Oligonucleotides for EMSA used in this study.**

| siRNA | Sequence (5'-3') |
|---|---|
| DNA1 | 5'-CATTGCATATTTAAAACATGTTGGATCCCACGTTGCATGCT |
| | GATAGCCTACTAGAGCTGTATGAATTCAAATGACCTCTTATCAAGTGAC-3' |
| DNA2 | 5'-GTCACTTGATAAGAGGTCATTTGAATTCATGGCTTAGAGCT |
| | TAATTGCTGAATCTGGTGCTGGGATCCAACATGTTTTAAATATGCAATG-3' |
| RNA1 | 5'-6-FAM-GUGCUACGAUGCUAGUCGGGGAGUGC |
| | ACCAGAUUCAGCAAUUAAGCUCUAAGCC-3' |
| RNA2 | 5'-6-FAM-GCACCAGAUUCAGCAAUUAAGCUCUAAGCC-3' |
